# SLALOM: A Simple and Rapid Method for Enzymatic Synthesis of CRISPR-Cas9 sgRNA Libraries

**DOI:** 10.1101/2020.07.21.175117

**Authors:** Joshua D Yates, Robert C Russell, H Joseph Yost, Jonathon T Hill

## Abstract

CRISPR-Cas9 sgRNA libraries have transformed functional genetic screening and have enabled innovative CRISPR-based methods, such as the visualization of chromatin dynamics in living cells. These libraries have the potential to be applied to a vast number of biological systems and aid in the development of new technologies, but their synthesis is hindered by the cost, time requirements, and technical difficulty of current sgRNA library generation methods. Here, we describe SLALOM—a rapid enzymatic method for generating robust, variant-matched sgRNA libraries from any source of DNA in under 3 hours. This method utilizes a custom sgRNA scaffold sequence and a novel method for detaching oligonucleotides from solid supports using a strand displacing polymerase. Using this method, we have constructed libraries targeting the *E. coli* genome and the transcriptome of developing zebrafish hearts, demonstrating its potential to expand the reach of CRISPR technology and facilitate methods requiring custom sgRNA libraries.

## INTRODUCTION

Although originally harnessed for editing single genomic loci^1,2^, the *Streptococcus pyogenes* CRISPR-Cas9 system has since been expanded to a number of high-throughput molecular techniques utilizing oligonucleotide libraries encoding tens of thousands of unique single-guide RNAs (sgRNAs) to target a large number of genetic loci^3,4^. These genome-wide libraries have typically been cloned into lentiviral vectors for forward-genetic screening cells in culture^3,5–8^. However, more recent studies have also adapted them for novel techniques, such as chromosome painting^9,10^ and gene regulatory analysis^11^, expanding the potential for sgRNA library-based methods.

A major hurdle in the application of the CRISPR sgRNA libraries is in generating the library itself. Designing and chemically synthesizing a custom sgRNA library using microarray-based or similar technologies^12^ can be expensive and time consuming and requires detailed sequence information and specialized bioinformatics tools. Because they are based on a reference sequence, chemically synthesized libraries can also differ significantly from the actual subject of the study, limiting their application to a few model organisms with well-assembled genomes and low rates of DNA sequence polymorphism. To address these issues, several enzymatic approaches have been developed to create libraries from various DNA inputs^10,13–15^. Using an enzymatic approach eliminates the need for subject-specific genomic sequence data, greatly diminishes the number of sequence mismatches between the library and subject, and can be synthesized at a fraction of the cost and time of chemically synthesized libraries. However, these methods can be difficult to carry out, require large amounts of DNA, or result in libraries where most of the sgRNAs are non-functional.

Here, we describe a streamlined enzymatic sgRNA library generation method that produces high quality libraries from small amounts of input material. By designing a custom sgRNA sequence containing a restriction endonuclease recognition sequence within the repeat:anti-repeat duplex, we were able to develop a method that requires just two consecutive sets of restriction digests and ligations. Additionally, our method constructs the library on the surface of magnetic beads—minimizing the loss of material and simplifying purification between steps. This method, which we have named SLALOM (sgRNA Library Assembly by Ligation On Magnetic beads), can be carried out in a few hours and with less than 1 μg of DNA or cDNA as input. Using SLALOM, we have generated sgRNA libraries from *E. coli* genomic DNA and normalized cDNA from developing zebrafish hearts and show that libraries generated by this method are effective *in vivo* through a fluorescent knockout screen in cell culture. Our data show that SLALOM is a simple, rapid, cost-effective method for generating sgRNA libraries to be used in a wide range of biological systems and techniques.

## RESULTS

### Method Design

To overcome the drawbacks of current enzymatic sgRNA library generation methods, we developed the SLALOM method (outlined in Figure 1, complete protocol in Supplemental Note 1). First, the restriction enzyme HpaII (CCGG), which recognizes a DNA sequence containing a PAM, is used to fragment the DNA (Step 1). HpaII was selected because its short palindromic recognition sequence ensures high coverage and cuts adjacent to PAMs on both the forward and reverse strands^10^. Next, an adapter is ligated to the ends of the digested fragments (Step 2). This adapter contains a custom sgRNA scaffold sequence modified to incorporate an MmeI recognition sequence and a long 5’ overhang. After ligation, the products are immobilized on magnetic beads by hybridization of the long single-stranded overhang to a complementary sequence affixed to the beads. After washing, the ligated DNA is digested by MmeI, which cleaves at a fixed distance from its recognition sequence^16^, leaving 18/19 bp of the ligated DNA sequence to form the sgRNA spacer (Step 3). After another wash, the fragments are ligated with a second adapter containing a promoter region (e.g. T7 for *in vitro* transcription or U6 for *in vivo* transcription) (Step 4). Finally, the beads are washed, and the completed library detached and repaired using a strand-displacing polymerase (Step 5).

**Figure 1.**
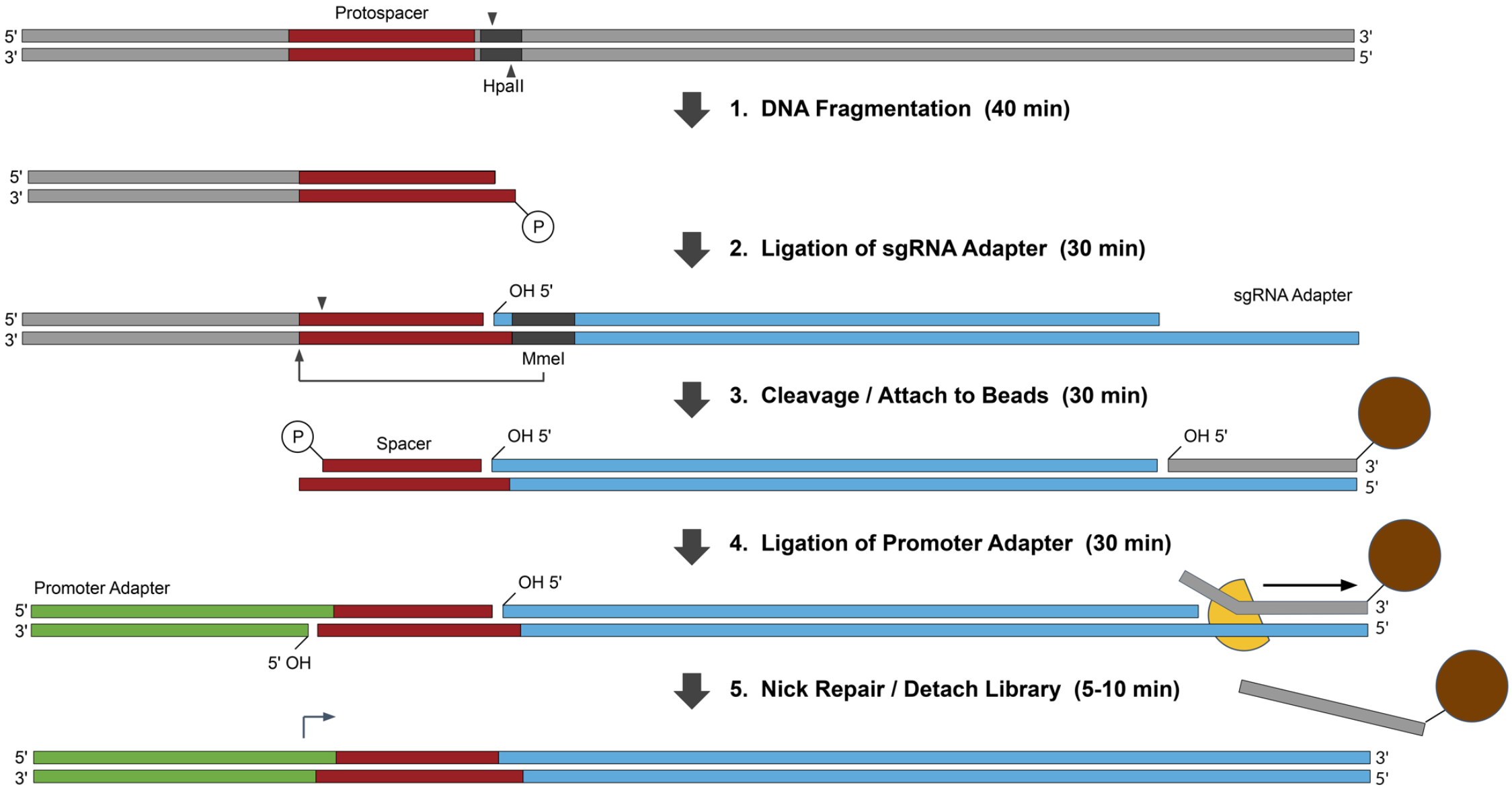
Schematic of the sgRNA Library Synthesis Method. Each step of the process is represented by a black arrow. Restriction enzyme binding sites are shown in black. Magnetic beads are represented as orange circles. (Step 1) DNA from an arbitrary source is fragmented by a restriction enzyme such as HpaII (CCGG) that contains the canonical PAM (NGG) in its recognition sequence. (Step 2) An adapter (blue) encoding a modified sgRNA sequence is ligated to the DNA fragments. The adapter is unphosphorylated, contains an MmeI recognition sequence, has a two base-pair, single-stranded overhang compatible with the fragmented DNA, and a long single-stranded overhang capable of hybridizing to a single-stranded oligo. (Step 3) A single-stranded capture oligonucleotide attached to a magnetic bead at the 3’ end is used to immobilize the adapter to magnetic beads through hybridization and excess DNA is washed away before digestion with MmeI to capture the spacer sequence. (Step 4) An unphosphorylated adapter containing the T7 promoter sequence (green) and a 3’ overhang containing all possible dinucleotides is ligated to the cleaved spacer sequence. (Step 5) The nicked library is detached from the beads by a strand displacing polymerase (yellow) and nicks are simultaneously repaired by the same mechanism. The detached double stranded oligonucleotides constitute an sgRNA library containing spacers targeting the DNA source and can be transcribed *in vitro* or cloned into plasmids for downstream applications.

### Modifications to the sgRNA Sequence

The method described here relies on a dual-role adapter that can function as a binding site for MmeI and as a template for transcription of the sgRNA scaffold, thus requiring modifications to the Cas9 binding region of the sgRNA without disrupting endonuclease activity. A previously reported crystal structure of Cas9 in complex with the sgRNA revealed that the REC1 domain of Cas9 interacts with the repeat:anti-repeat duplex of the sgRNA^17^. Functional experiments in this same study showed that some mutations to the sequence of the duplex are permitted, while others can diminish or completely inhibit the Cas9 activity. It was therefore unclear if an MmeI recognition sequence could be incorporated into the repeat sequence without impairing the function of the Cas9 complex.

Based on this information, we incorporated the MmeI recognition sequence (TCCRAC) into the repeat sequence of the sgRNA at position 21 (directly after the spacer region) by making two mutations, U24G and U25G, in the repeat sequence. We also made corresponding mutations, A46C and A47C, in the anti-repeat sequence to maintain the repeat:antirepeat stem loop structure (Figure 2A). Digestion of a 1000 bp DNA fragment *in vitro* with Cas9 and the wild type or modified sgRNA showed cleavage efficiencies similar to or better than the unmodified sgRNA with both an 18 bp and 20 bp spacer, indicating that endonuclease activity of the protein was retained (Figure 2B, lanes 4 and 5).

**Figure 2.**
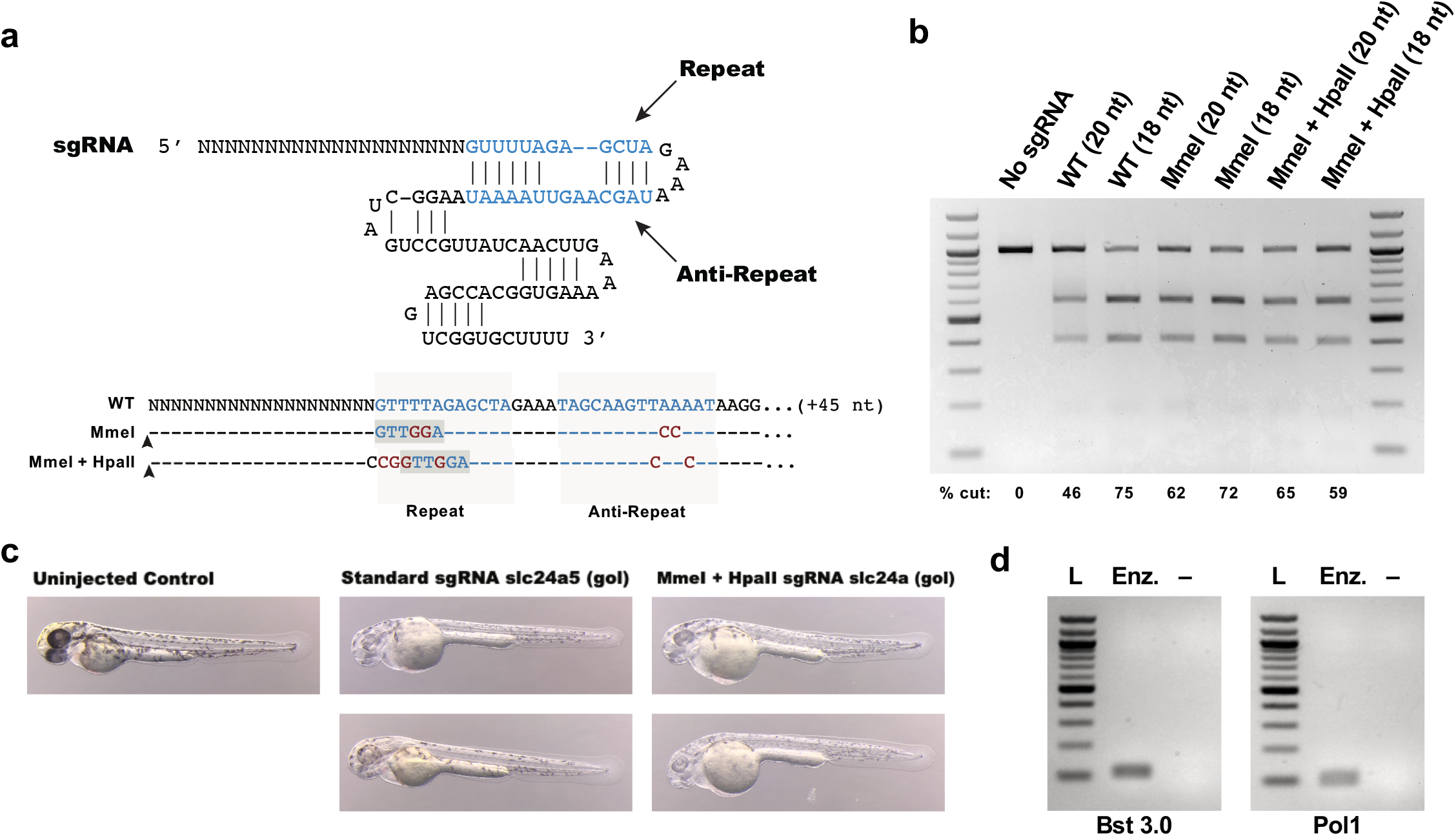
Incorporation of the MmeI Recognition Sequence into the Repeat:Anti-Repeat Duplex and Elution with a Strand Displacing Polymerase. A) Modifications were made to the Repeat:Anti-Repeat Duplex of the sgRNA sequence to insert a single MmeI site that cleaves 18-20 bp upstream of the sgRNA recognition sequence. Mutations are shown in red. The MmeI binding sites are shown in dark gray boxes. The MmeI cut site is represented with a triangle. B) *in vitro* Cas9 digestion of a 1000 bp DNA fragment with each sgRNA variant. C) Injection of zebrafish embryos with Cas9 and the sgRNA variant targeting the *gol* gene compared to an uninjected control (minimum n per pool = 150) D) Detachment of the sgRNA scaffold from magnetic beads using Bst 3.0. E) Detachment of the sgRNA scaffold from magnetic beads using Pol1.

The enzymes used to fragment the DNA prior to ligation with this adapter also produce short 3’ overhangs not compatible with the wildtype Cas9 sgRNA sequence. Therefore, we next engineered additional mutations to allow the sgRNA adapter sequence to hybridize with the fragmented ends, eliminating the need for removal of the overhangs and increasing ligation efficiency. This was accomplished by shifting the MmeI recognition sequence to position 23 and modifying the sgRNA sequence by making mutations G21C, T22G, T23G, and A26G in the repeat sequence and T45C, A48C and A49C in the anti-repeat sequence. Moving the MmeI site to this position resulted in guide sequences of 18 and 19 bp (Figure 2A) instead of the standard 20 bp. However, digestion of the 1000 bp DNA fragment *in vitro* using an sgRNA with these modifications again showed cleavage at the predicted location with similar efficiency to that of the unmodified sgRNA using either a 20 bp or 18 bp protospacer (Figure 2B, lanes 6 and 7).

To confirm that sgRNAs with these modifications function *in vivo*, we tested an sgRNA with these modifications and a spacer sequence targeting the zebrafish pigment gene *gol* (slc24a5)^18^. Injection of single-cell zebrafish embryos with the modified sgRNA showed levels of pigment loss at 48 hpf comparable to an unmodified sgRNA targeting the same spacer (Figure 2C). Together, these results demonstrate that modified sgRNAs can be used to guide Cas9 editing *in vitro* and *in vivo*.

### Detaching Oligonucleotides with a Strand Displacing Polymerase

We also sought to simplify purification and handling of the library during construction by incorporating magnetic bead purification into the SLALOM workflow. Constructing the library on the surface of magnetic beads allows for rapid purification between steps without disrupting the function of restriction enzyme or T4 DNA ligase activity^19^. To immobilize the modified sgRNA adapter on the beads, a biotinylated ssDNA oligo was designed to hybridize with the 3’ end of the modified sgRNA adapter and a corresponding 19 base overhang was designed into the 3’ end of the adapter. The ssDNA oligos were then attached to streptavidin-coated paramagnetic beads during the SLALOM protocol. Elution was accomplished by incubating the beads with a strand-displacing polymerase, which elongates the 3’ end of an oligo while displacing the pre-existing strand, detaching the final library product from the beads (See Figure 1). Two strand displacing polymerases were tested: Bst 3.0, which is most efficient at temperatures near the melting temperature of the bead-attached oligo, and Pol I, which is active at room temperature and contains 5’-3’ exonuclease activity, which completely digests the oligo being displaced. Oligo hybridization and elution assays showed that both enzymes were highly efficient (Figure 2), thus providing a simple technique to irreversibly detach oligonucleotides from solid supports while simultaneously repairing the nicks caused by using unphosphorylated substrates in the earlier ligation steps.

### GFP Library and Fluorescent Knockout Screen

To assess functionality of a SLALOM generated library in cell culture, we used SLALOM to create a library targeting the GFP coding region in GFP-LC3 Hela cells^20^. The GFP construct in these cells contains four HpaII sites (CCGG) (Figure 3A). We PCR amplified the GFP coding region (716 bp) to use as the SLALOM DNA input. In this iteration, the sgRNA adapter was also modified to use a longer stem-loop, shown to potentially increase sgRNA efficiency^9^, and both adapters incorporated Esp3I restriction sites for cloning into the LentiCRISPRv2 plasmid^21^. Library sequencing confirmed complete coverage of the 8 expected sgRNAs (Supplemental Figure 1). The cloned library was then packaged into lentivirus and used to transfect HeLa GFP-LC3 cells. Sorting of the transfected and untransfected cells by FACS showed that approximately 15% of the transfected cells lost GFP expression (Figure 3B, Supplemental Figure 2), comparable to previous studies looking at CRISPR efficiency^22^. Sequencing of the incorporated spacers and GFP mutations in each pool showed that all eight potential spacers could be found in the GFP-negative pool and mutations occurred at all four sites (Figure 3C). However, it was not possible to identify the mutation rate of each individual sgRNA, as many of the common mutations deleted both cut sites within a pair (Figure 3D).

**Figure 3.**
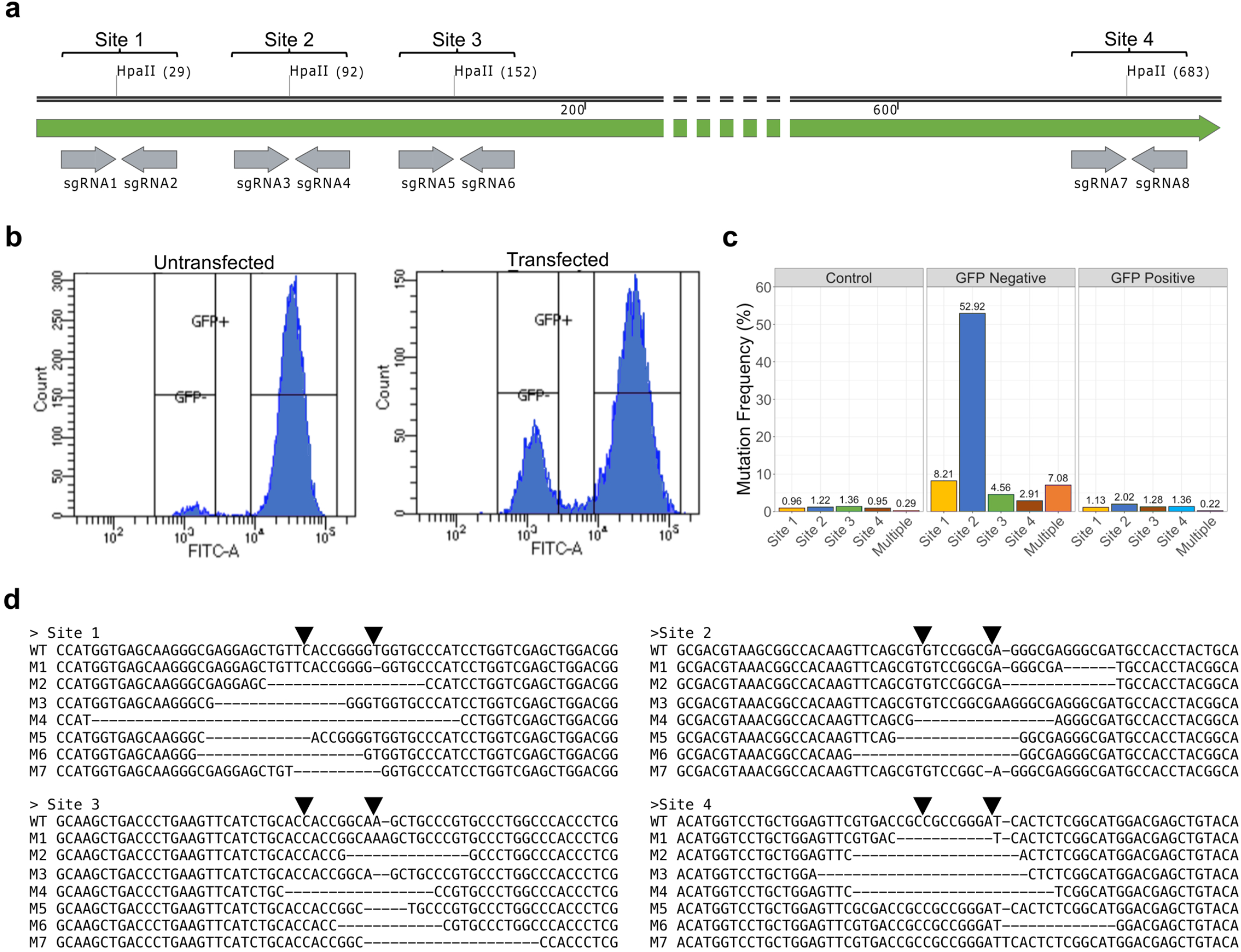
GFP Knockout Screen. A) Genomic DNA was isolated from HeLa cells stably expressing GFP-LC3. The four HpaII sites within the GFP ORF and the predicted sgRNAs are labeled. B) FACS sorting of transfected and untransfected cells. GFP fluorescence is on the x-axis and counts are on the y-axis. C) Mutation frequency at each of the HpaII sites based on high-throughput sequencing of the GFP locus. D) The most common variants for each of the four HpaII sites.

After establishing that all of the sgRNAs were active, we next analyzed their relative mutation efficiency. Relative enrichment of each sgRNA in the GFP minus pool was determined by normalizing the fractional counts from the sgRNA spacer sequencing results of the GFP minus pool to their fractional counts in the GFP positive pool (Figure 4A). It should be noted that this analysis is not completely analogous to enrichment analyses in genome-wide screens, as there is no background gene set for normalization, but does show the relative efficiency of each sgRNA in this library. These data showed that one sgRNA, a 19 bp sequence targeting the negative strand of site 2, was more active than the others. There were also two sequences that appeared to be significantly less effective than the average, an 18 bp sgRNA targeting site 3 and an 18 bp sgRNA targeting site 4.

**Figure 4.**
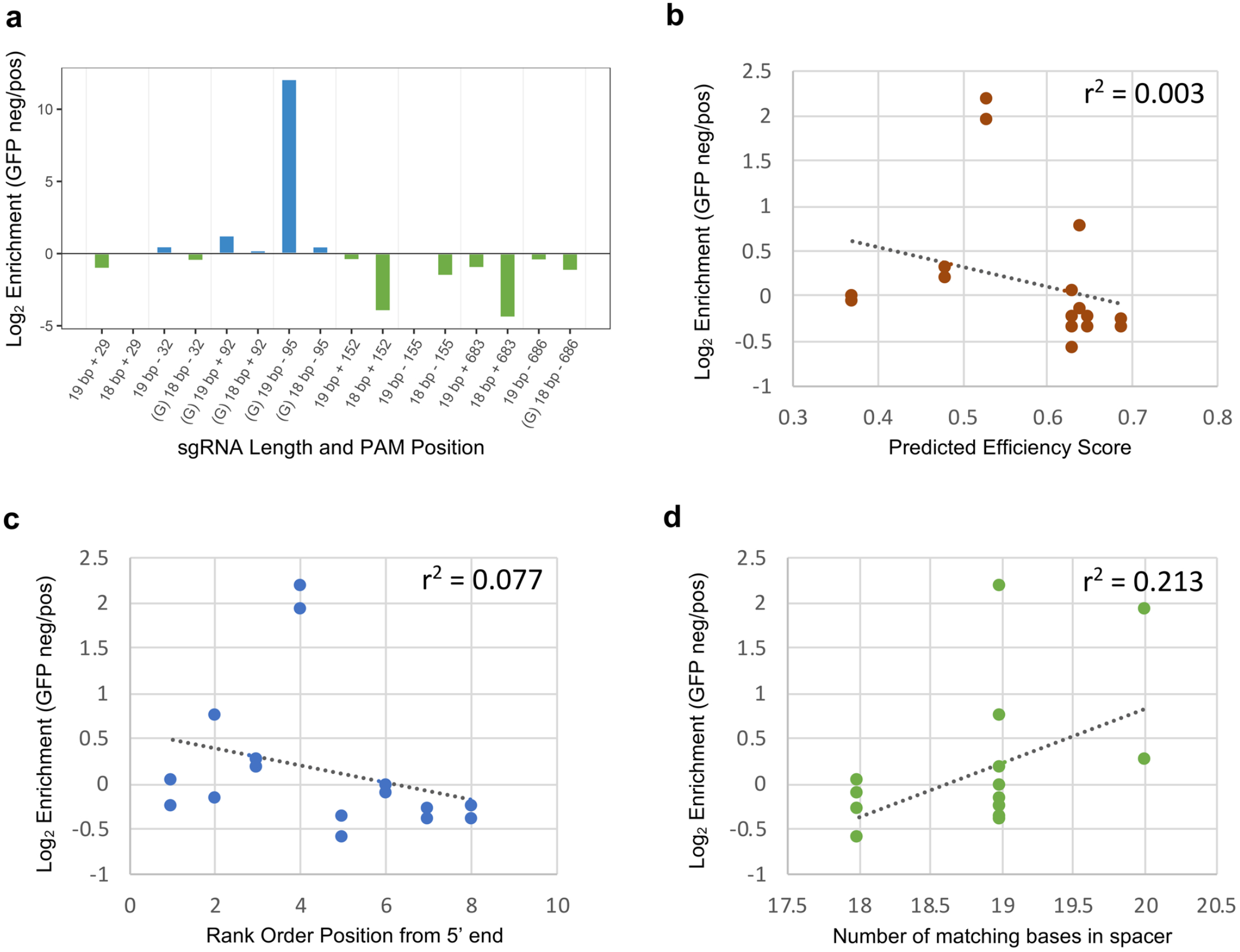
Relative Efficiency of sgRNAs in the GFP Library. A) Relative efficiency of each sgRNA measured by fold enrichment in the GFP-minus HeLa cell pool compared to the GFP-positive pool. B) Comparison of relative efficiency in the screen vs. predicted efficiency scores^23^ for the locus. C) Comparison of relative efficiency in the screen vs. location in the GFP coding region. D) Comparison of relative efficiency vs. length of the match between the spacer and protospacer.

We initially hypothesized that this variance may be due to the fact that SLALOM does not select for sgRNAs based on predicted efficiency. However, enrichment in the GFP-negative pool did not show strong correlation with predicted efficiency score^23^ (Figure 4B, r^2^=0.003), position in the gene (Figure 4C, r^2^=0.077) or length of the match (Figure 4D, r^2^=0.213). Overall, these results show that the library can effectively target multiple sites and generate a measurable enrichment of mutations causing a specific phenotype of interest, despite not using bioinformatics analysis during the design the library. Furthermore, digestion with HpaII is expected to generate an average of 13 sgRNAs per gene, much higher than current synthesized libraries, providing additional opportunities to include highly efficient guides.

### E. coli and Looping Zebrafish Heart Libraries

To test the method on a more complex DNA substrate, we isolated genomic DNA from *E. coli* MG1655 and created an sgRNA library using SLALOM (Figure 5A). The E. coli genome contains 24,311 HpaII sites. Because SLALOM produces two sgRNAs, one on the plus strand and one on the minus strand, for each site, we predicted a full-coverage library would contain 48,622 sgRNA templates. High-throughput sequencing of the spacers was then conducted to characterize the resulting sgRNA library. Sequencing results showed that the most common spacer lengths were 18 and 19 bases (Figure 5B) and were evenly represented in the library (Figure 5C). Finally, sites in the library covered 91.6% of the predicted HpaII sites and that 98.3% of the reads represented a PAM-adjacent spacer (Figure 5D), indicating that SLALOM can be used to create high quality sgRNA libraries from a high molecular weight DNA input.

**Figure 5.**
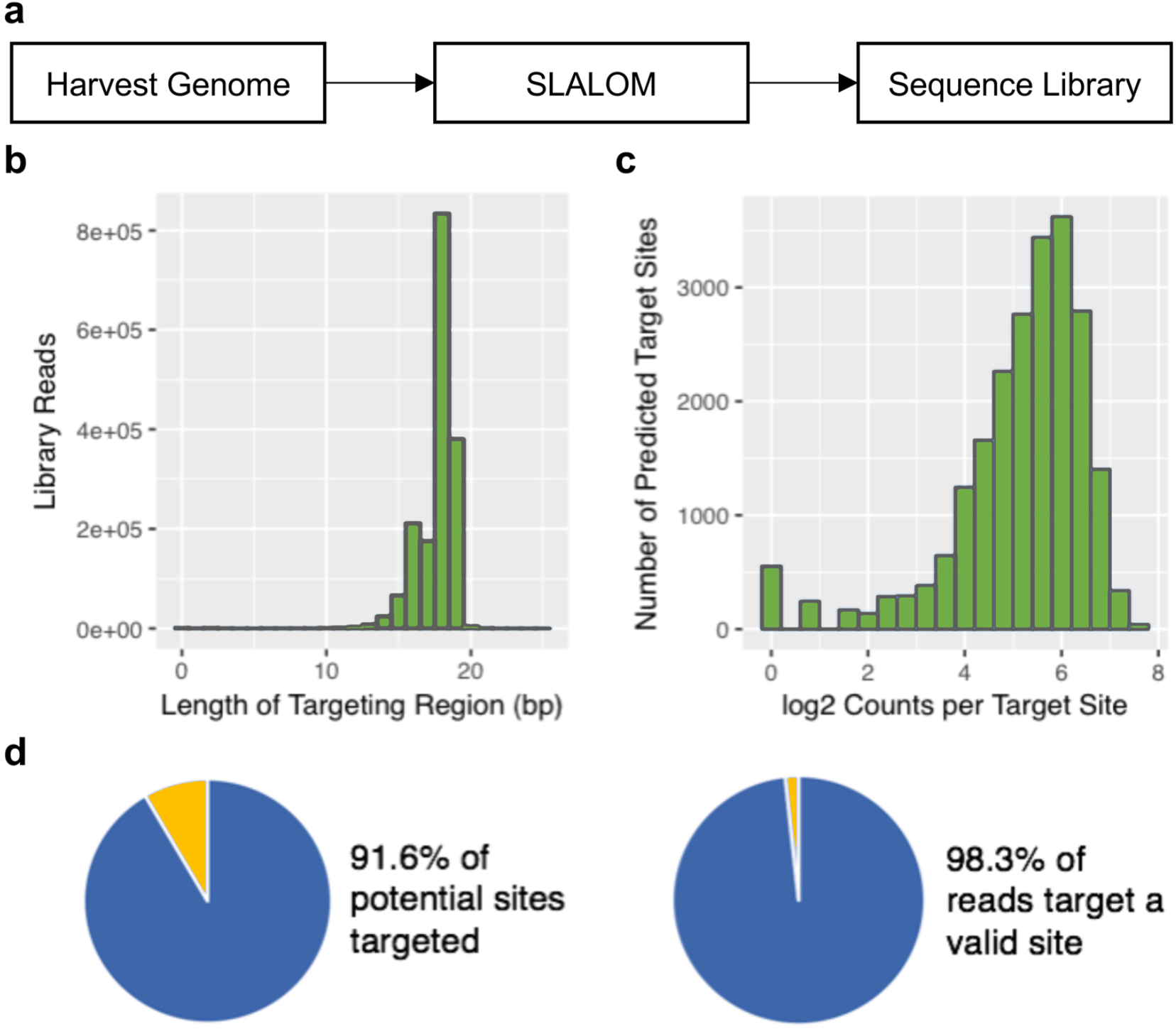
High-throughput sequencing of a SLALOM-generated sgRNA library to the E. coli MG1655 genome. A) Schematic diagram of the library generation process. B) Histogram showing the distribution of spacer lengths within the library. C) Histogram showing representation of each sgRNA target in the library. D) Analysis showing the coverage and purity of the library.

A clear advantage of SLALOM over chemically synthesized libraries is the ability to create tissue or cell-type specific libraries without prior information on the genes expressed in those cells. Thus, we next tested the ability of SLALOM to create an sgRNA library targeting all of—but only—the genes expressed in the developing zebrafish heart (Figure 6A). Zebrafish hearts were isolated from 48 hours post-fertilization (hpf) embryos for mRNA isolation. Because genes are expressed at various levels, we next created normalized zebrafish heart cDNA by subtractive hybridization^24^. The normalized cDNA library was confirmed by qPCR (Figure 6B) and used as input for SLALOM.

**Figure 6.**
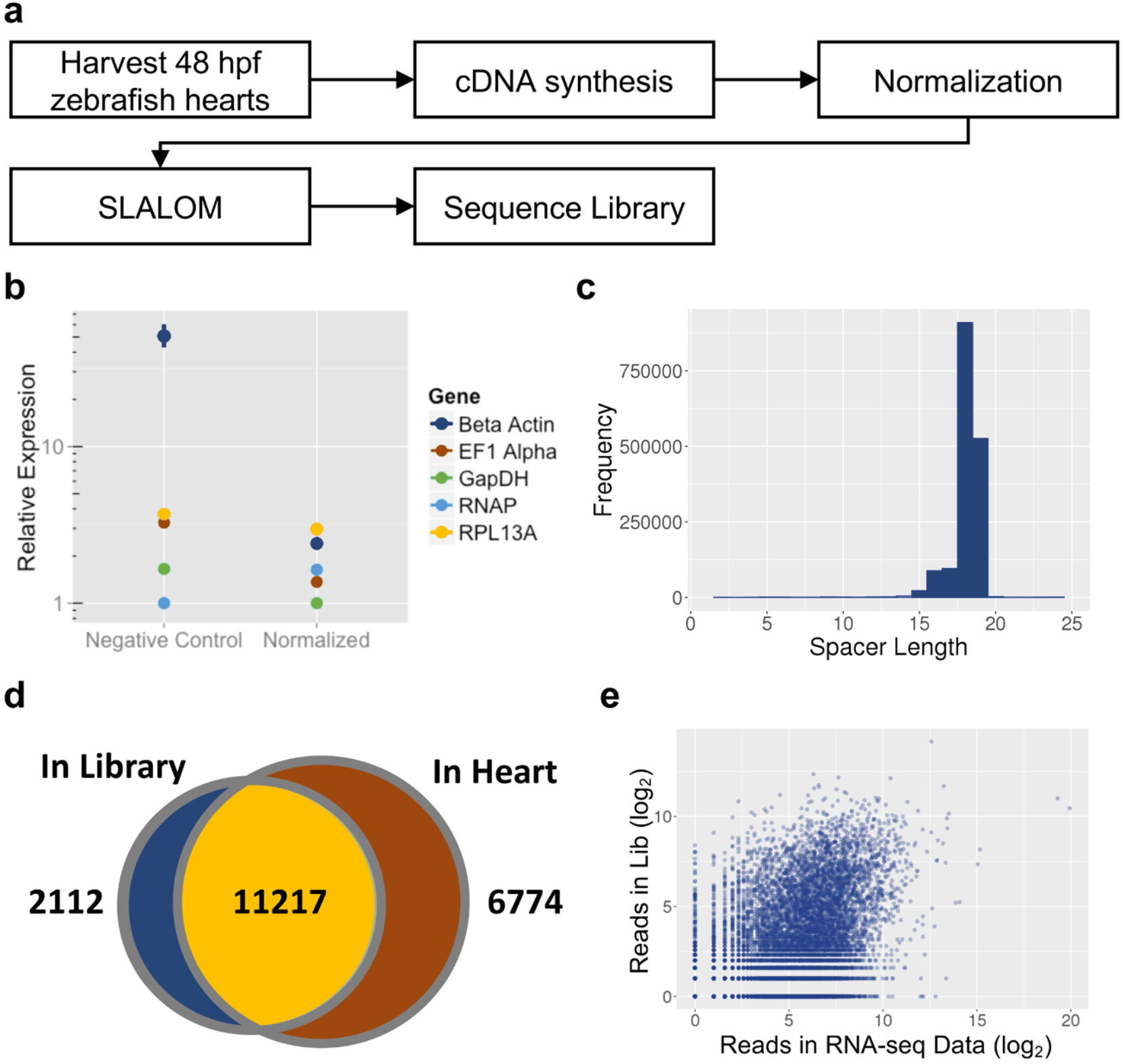
High-throughput sequencing of a SLALOM-generated sgRNA library to normalized cDNA extracted from a 48 hpf zebrafish heart. A) Schematic diagram of the library generation process. B) Results of qPCR analysis on five genes with varying expression levels in the endogenous tissue before and after cDNA normalization. C) Histogram showing the distribution of spacer lengths within the library. D) Venn diagram of the genes targeted by at least one spacer in the library compared to genes detected in the 48 hpf zebrafish heart by RNA-seq. A threshold of 1 read per gene was used to designate both genes and spacers as “detected”. E) Comparison of the total number of spacer reads vs. expression level in an RNA-seq dataset for 48 hpf zebrafish hearts.

Similar to the *E. coli* genome library, sequencing validation of the zebrafish heart library found spacers that were predominately 18 or 19 bp long (Figure 6C). As sgRNAs in this library were expected to only target expressed genes, we compared the library to a set of genes detected in an existing zebrafish heart RNA-seq dataset^25^. These data showed the library targets at least 11,217 of the genes expressed in the heart (Figure 6D). Representation in the library is not correlated with gene expression levels in the heart, confirming that cDNA normalization was effective (Figure 6E). Thus, SLALOM can be used to target an sgRNA library to a specific tissue.

## DISCUSSION

Generation of sgRNA libraries by chemical synthesis can be expensive, slow, and require significant bioinformatic analysis of a known genome to identify suitable spacers. Because of these limitations, synthesized libraries have been largely limited to a few well-established model organisms with low DNA sequence polymorphism rates (such as inbred mice and human cell lines) and complete genomes. However, these libraries cover only a small fraction of the biological systems where they could be used. Wider adoption of CRISPR library methods in non-traditional organisms and the development of new CRISPR-based technologies require the development of more attainable library generation methods.

One alternative to chemical synthesis is enzymatic processing of DNA inputs into sgRNAs. Several such methods have been proposed. Lane et al.^10^ digested DNA with restriction enzymes containing a PAM in their recognition sequences and after ligation of a temporary adapter used the restriction enzyme MmeI to capture the spacer before removing this adapter and attaching an adapter containing the sgRNA sequence. This method resulted in an E. coli genome library where 44% of the sequenced reads represented functional guides and 51% of the predicted spacers were included. In contrast, the *E. coli* library presented here achieved results of 98.3% and 91.6%, respectively, for these same analyses. Arakawa, et al.^13^ also used a restriction enzyme that cleaves outside of its recognition sequence to capture the spacer but selected for PAMs by creating a cDNA library using a semi-random hexamer containing the dinucleotide CC. This resulted in a library where 77.6% of the spacers were adjacent to the PAM. In contrast, Cheng et al.^14^ and Köferle, et al.^15^ did not select for PAMs but produced libraries from completely random fragments of a DNA sample. As a result, Cheng et al. produced dense libraries targeting practically all valid PAMs and Köferle, et al. produced libraries where 23-30% of the spacers were adjacent to PAMs and the spacer lengths distributed at about 27 bp. However, both libraries contained many spacers that did not target PAM sequences. Finally, library generation and screening have been combined in prokaryotes by transfecting the input DNA directly into bacteria with a hyper-stimulated CRISPR system^26^. Thus, there was a wide range of issues in the previous approaches, including complexity, reliability, difficulty, time requirements, input amounts, and library quality and density.

SLALOM has several advantages over these methods, as it requires fewer steps and smaller amounts of starting material while yielding sgRNA libraries in which almost all of the final output contains functional guide sequences. Similar to the Lane et al. method described above, SLALOM takes advantage of a restriction enzyme to determine the PAM, as this method makes it possible to use any source of DNA input (e.g. genomic DNA, cDNA, or PCR products) and to tune the density of the library by choosing enzymes with different recognition sequence lengths and/or combining multiple restriction enzymes. However, unlike the Lane et al. method, SLALOM uses a two-base overhang incorporated into the adapter sequence, eliminating the need for mung-bean nuclease blunting of the digested fragments. Several previously reported methods^10,13,14^ also temporarily ligate an intermediate adapter containing a recognition sequence for a restriction enzyme that cleaves outside of this sequence to capture approximately 20 bp of the fragment before removing the adapter. In SLALOM, this restriction enzyme site is directly incorporated within the sgRNA template, eliminating the need for an intermediate adapter. Using a single adapter also allows the reaction steps to occur while attached to a magnetic bead, making purification between steps fast and simple and reducing material loss.

While SLALOM can already make high quality libraries, it also serves as a platform to expand and diversify gene-targeting methods. For example, other restriction enzymes could be used to create different sets of sgRNAs or in combination to increase library density. There might also be ways to improve the mutation frequency, such as using a Type IIS enzyme with a longer reach to generate sgRNAs that are 20-21 instead of 18-19 bases long. Based on previous studies, this would likely increase cutting efficiency but could also decrease specificity^27,28^. Finally, we made libraries using genomic DNA and normalized cDNA, but other sources of the DNA input, such as pull downs to obtain DNA fragments bound by various proteins, could be used as starting material. For example, conducting an RNA-PolII pulldown could make the representation of genes actively transcribed in the tissue of interest more consistent than normalized cDNA while still limiting the library to active genes, and pulldowns of other DNA-binding proteins could also be used to examine subsets of enhancers or other non-coding regions, especially if combined with repressor- or activator-bound Cas9 variants^5,29–31^.

One potential application of SLALOM-generated sgRNA libraries is in forward-genetic screening. CRISPR screening in cell culture is already well established and has been shown to be more effective than chemical mutagenesis or siRNA-based methods^8,32^. However, current sgRNA libraries are generally synthesized to target all of the genes in the genome, which reduces the number of times each gene can be targeted and increases the number of cells without mutations in active genes, limiting the rate of gene discovery. However, enzymatic sgRNA generation allows the library to be tailored to the genes expressed in the tissue and at the time point of interest. For example, the zebrafish heart library reported here contains only sgRNAs targeting the exons of genes expressed during heart looping morphogenesis. Thus, this method will restrict mutations to genes involved in heart development, greatly improving the rate of gene discovery and potentially expanding the number of biological systems that can be screened using CRISPR technology beyond cell culture. Similar applications of SLALOM to novel experimental designs and organisms will greatly increase the reach of CRISPR/Cas9 technology in a wide range of fields.

## EXPERIMENTAL PROCEDURES

### Oligonucleotides and Bead Preparation

*In vitro* transcription of sgRNAs was accomplished by annealing and extending pairs of oligos (see Supplemental Table 1 for oligos ordered from Integrated DNA Technologies (Coralville, USA) or Eurofins (Louisville, KY)) by incubating at 66°C for 20 minutes to produce double-stranded DNA templates containing a T7 promoter sequence, spacer sequence, and sgRNA scaffold sequence and column purified. Either the MEGAscript™ T7 Transcription Kit (Thermo Fisher Scientific) or the HiScribe™ T7 Quick High Yield RNA Synthesis Kit (New England Biolabs, Ipswich, USA) was used to transcribe the templates. In either case, 210-300 ng of the template was used and incubated at 37°C for 1-2 hours. DNase I was then added, and the reaction incubated for an additional 15 minutes at 37°C. The sgRNA was then purified using either the RNA Clean & Concentrator-5 Kit (Zymo Research, Irvine, USA) or by phenol-chloroform extraction and ethanol precipitation.

Capture beads were prepared by resuspending 50 μL of streptavidin magnetic beads (New England Biolabs, Ipswich, USA) in 20 μL containing 100 pmol of the biotinylated oligo and incubating at room temperature for 15 minutes. The beads are then washed twice by resuspending the beads in 50-100 μL of 1x Cutsmart^®^ buffer. The ability of the biotinylated oligo to bind to the magnetic beads was similar in both the recommended binding buffer as well as CutSmart^®^ buffer.

### *In vitro* Digestion with Cas9

*In vitro* digestion of DNA fragments was accomplished by adding 200 ng of DNA, 70 pmol Cas9 Nuclease, S. pyogenes (1,000 nM) (New England Biolabs, Ipswich, USA), and 100 ng of the sgRNA in NEBuffer 3.1 (New England Biolabs, Ipswich, USA) for 20 minutes at 37°C and 10 minutes at 65°C, followed by adding 1 μL of proteinase K (800 units/ml) (NEB) and incubating at room temperature for 10 minutes before being run on a 1.7% agarose gel with a 100 bp ladder for 35 minutes.

DNA adapters were prepared by resuspending complimentary oligos at a final concentration of 10 μM of each in CutSmart^®^ buffer (New England Biolabs, Ipswich, USA). The reaction was then heated to 98°C for 2 minutes then set to ramp from 85°C to 65°C for 1 hour and finally from 65°C to 8°C for 30 minutes.

### SLALOM Library Construction Method

A complete protocol for the SLALOM method can be found in Supplementary Note 1. Briefly, DNA containing about 10 pmol of recognition sites for HpaII (CCGG) was added to a 50 μL reaction with a final concentration of 1x CutSmart^®^ buffer and 10 units of HpaII. The reaction was incubated at 37°C for 20 minutes and heat inactivated at 80°C for 20 minutes. 2,000 units of T4 DNA ligase, 10 pmol sgRNA adapter, and 50% PEG 6000 w/v were added to the reaction at room temperature to bring the reaction to a total volume of 75 μL and 7.5% PEG. The reaction was then incubated at room temperature for 20 min. The reaction was mixed with capture beads for 15 minutes and the beads washed twice by exchanging the buffer for 50 μL 1x CutSmart^®^ buffer. The beads were then resuspended in 50 μL of 2 units MmeI and 50 µM S-adenosylmethionine (SAM) in 1x CutSmart^®^ buffer and incubated at room temperature for 20 min while keeping the beads in solution by occasionally pipetting. Beads were washed as described above and resuspended in 50 μL of 1x CutSmart^®^ containing 2,000 units of T4 and 30 pmol of the T7 adapter in 1x CutSmart^®^. The reaction was incubated for 20 minutes at room temperature. The beads were washed and resuspended in 50 μL of 1x CutSmart^®^ containing 10 units of DNA Polymerase 1 and 200 µM dNTPs in 1x CutSmart^®^ and incubated for 5-10 minutes at RT. A DNA Clean and Concentrate column (Zymo Research, Irvine, USA) was used to purify the final library.

### Zebrafish Embryo Injections

Zebrafish (*Danio rerio*) embryos were collected and injected at the single-cell stage with Cas9:sgRNA Ribonucleoprotein (RNP) by incubating Cas9 (Integrated DNA Technologies, Coralville, USA) with sgRNA in a 300 mM KCl solution for 5 minutes before injecting approximately 1 nl of solution into each zygote. Injected embryos were kept at 28.5°C and after 2 days were visually examined for the extent of pigmentation. Photos were taken using an Olympus SZX16 microscope.

### Fluorescent Knockout Screening

GFP-LC3 HeLa cells stably expressing GFP were cultured in D10 medium (Dulbecco’s modified Eagle’s medium (DMEM), Fetal bovine serum (10%), L-glutamine, penicillin, Streptomycin) at 37°C with 5% CO_2_ and passaged every 3 days to maintain growth conditions. Genomic DNA from these cells was isolated by collecting cells by centrifugation at 1000 g for 5 min and resuspension in 1x RIPA buffer containing Pronase at 37 °C for 10-15 min followed by phenol chloroform extraction and ethanol precipitation. Genomic DNA was used as a template to amplify the GFP coding region using the GFP forward and reverse primers to generate a 716 bp fragment with 4 HpaII (CCGG) sites. This fragment was used as a substrate for SLALOM and cloned into the lentiCRISPRv2 plasmid (Addgene, cat. no. 52961). The library digested with Esp3I and purified using a DNA Clean & Concentrator-5 kit (Zymo Research, Irvine, USA). The plasmid was digested with Esp3I and NheI and the resulting 12,895 bp fragment was gel extracted. Ligation using T4 DNA ligase of the two fragments was followed by transformation into NEB^®^ Stable Competent *E. coli* cells (New England Biolabs, Ipswich, USA) where 100 μL was plated and 900 μL was used for an overnight culture in LB with 100 mg/ml Ampicillin. The overnight culture was used to inoculate a 500 mL LB with the antibiotic and the plasmid DNA was isolated using NucleoBond Xtra Midi EF (Macherey-Nagel, Dueren, Germany). Sequence flanking the spacer sequences were amplified and were sequenced. Following validation of the library, it was packaged into lentivirus by VectorBuilder (Chicago, USA).

To calculate the multiplicity of infection (MOI), two 6 well dishes were seeded at approximately 20% confluency and allowed to attach to the plates. Cells from the first dish were transduced with increasing amounts of the virus in media containing 6 μg/mL Polybrene. After 2 days of growth, media was replaced with media containing 8 μg/mL Puromycin for two additional days until all cells in the control well had died and the surviving cells were counted, and the ratio was used to calculate a curve. Cells infected at an estimtaed MOI of 0.3 and sub-cultured to obtain enough cells for sorting. After 10 days, cells were sorted based on fluorescence using a FACS Aria Fusion and the sorted cells were cultured separately. Genomic DNA from these two populations as well as from uninfected cells was isolated and the GFP coding region was sequenced. The spacers from these two populations were also sequenced.

### E. coli Genomic DNA Isolation

Genomic DNA from *Escherichia coli* MG1655 was isolated using the PowerLyzer^®^ UltraClean^®^ Microbial DNA Isolation Kit from Qiagen (Hilden, Germany). Bacterial cells were harvested and lysed using glass microbeads. Final product was run on a gel and a clear band of high molecular weight was observed. The genomic DNA was used as a substrate for SLALOM and the resulting library was sequenced.

### Developing Zebrafish Heart mRNA Extraction

Zebrafish hearts were isolated as published previously^33^. Briefly, about 200 Tg(myl7:GFP) zebrafish embryos were collected at 48 hours post-fertilization and placed in media containing tricaine. Embryos were resuspended in L-15 media with 10% FBS and the tissue of the embryos was disrupted by passing them through a 19G needle. Intact hearts were isolated under a fluorescent microscope with a pipette and after being washed were resuspended and homogenized in TRI-Reagent^®^ (Zymo Research, Irvine, USA). Zebrafish heart mRNA was then purified using the Direct-zol™ RNA Kit (Zymogen Research, Irvine, USA).

### cDNA Library Synthesis and Normalization

The SMART^®^ cDNA Library Construction Kit (Clontech) was used to reverse transcribe the zebrafish heart RNA. An alternative oligonucleotide that lacked an HpaII binding site was used in place of the 5’ oligo provided by the kit to prevent gRNA creation to the adapters in the cDNA library (Supplementary Table 1). The cDNA library was then normalized using the Trimmer-2 cDNA normalization kit (Evrogen, Moscow, Russia).

### High-throughput Sequencing and Data Analysis

High throughput sequencing of the libraries was carried out at the University of Utah Shared Genomics Resource on an Illumina HiSeq 2500. PhiX DNA was added to each library before sequencing to allow for more diversity during the initial reading process. Custom scripts were generated to extract the targeting region of each sgRNA template and analyze the data (Supplemental Notes 2 and 3). Because the length of the library was 128 bp, but the read length was only 50 bp, about half of the reads were discarded because they did not contain the spacer. The extracted spacer sequences were then aligned to either the Escherichia coli MG1655 genome or the zebrafish GRCz11 genome. Statistical analysis was then used to determine the coverage of each library.

Confirmation of the GFP mutation rates was conducted by high-throughput sequencing using the Genewiz Amplicon-EZ service (South Plainfield, USA) and analyzed using custom scripts in R based on the Cris.py pipeline^34^ (Supplemental Notes 4 and 5). Briefly, reads containing complete PCR products were selected and searched for a 30 bp sequence spanning each HpaII site. Reads were categorized by whether they contained wild type sequences at each site. Reads with identified mutations at each site were then tabulated to find the mutation frequency at the site and the most common mutations. Reads with large deletions that spanned multiple HpaII sites were categorized separately as “Multiple”.

## Supporting information

Supplemental Note 1: SLALOM Protocol

## DISCLOSURES

### IACUC Approval

All animal studies were approved by the Brigham Young University Institutional Animal Care and Use Committeee under protocol number 18-0704.

### Data availability

All high-throughput sequencing data used in this study are available through the Sequence Read Archive (Project ID: PRJNA642300).

## Acknowledgments

The Hela cells used here were a generous gift from Joshua L. Anderson. We would also like to thank Marc Hansen, Brent Nielsen, and Jeffery Barrow for their critical reading of the manuscript.

---

**Supplemental Figure 1.**
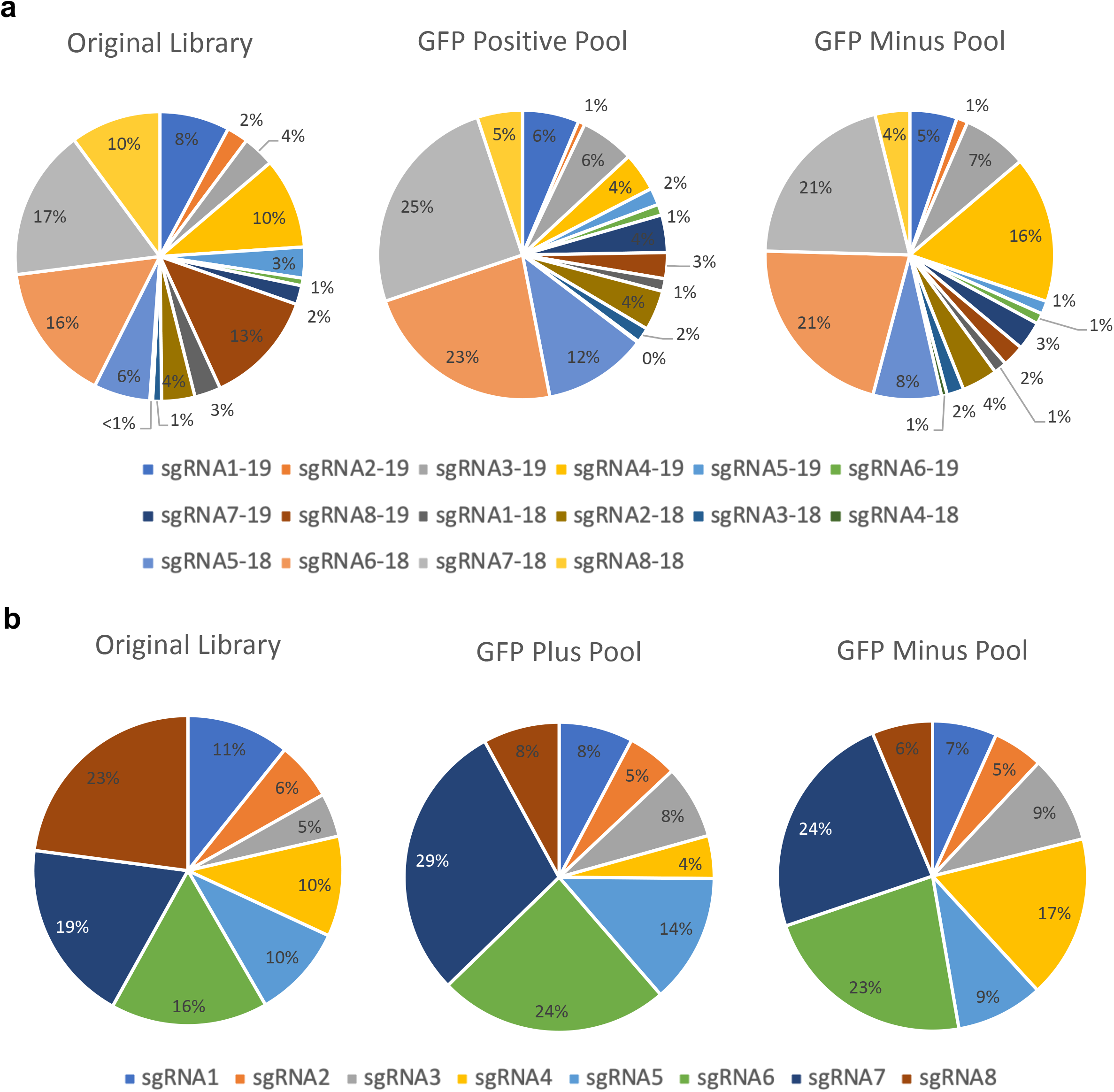
Relative representation of sgRNAs. The overall library, GFP Positive Cells after FACS sorting, and GFP Minus Pool after FACS sorting were sequenced to identify the relative representation of each potential sgRNA.

**Supplemental Figure 2.**
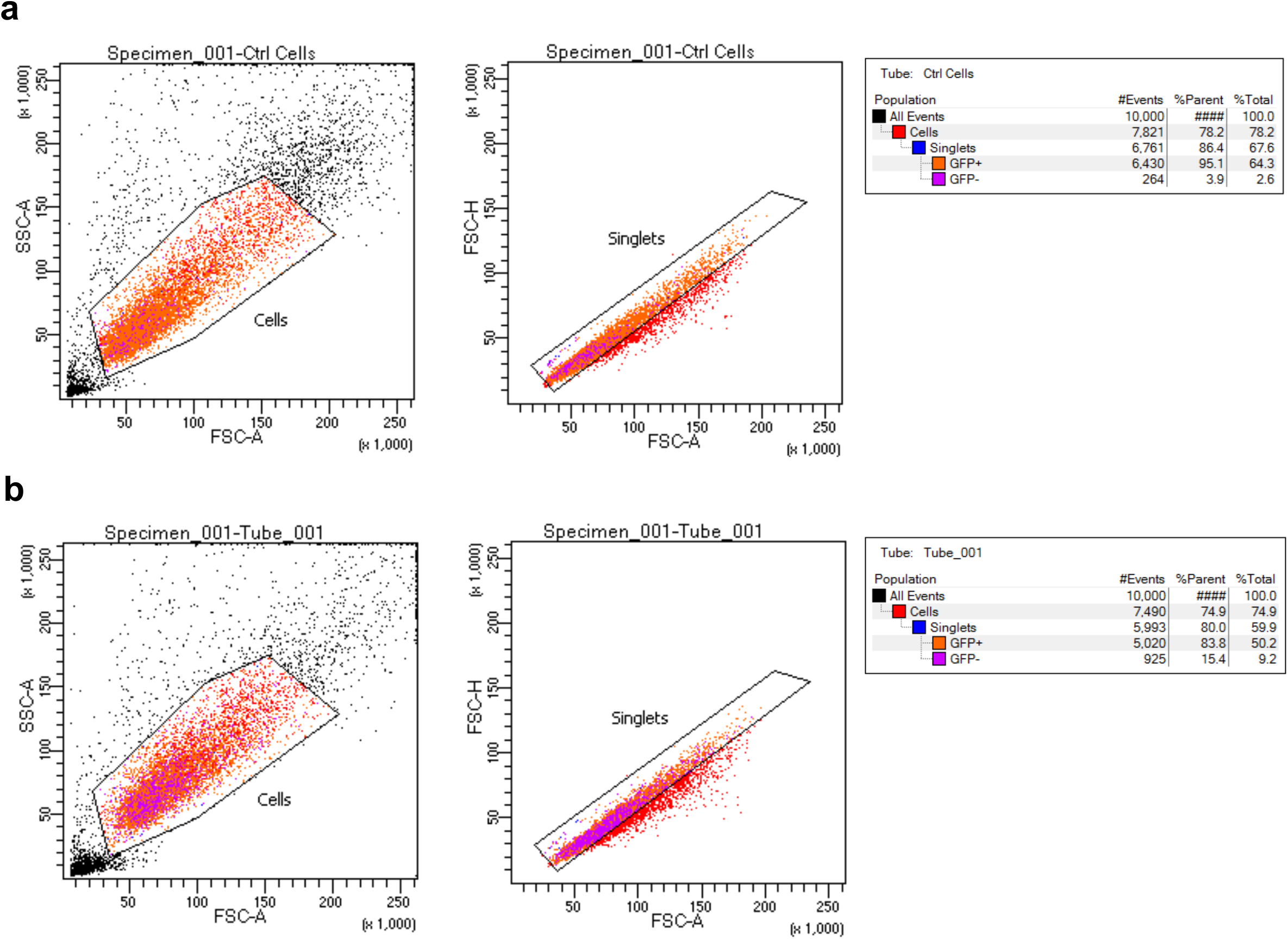
Raw FACS Results. A) Raw FACS results for live cell gating for untransfected cells. B) Raw FACS results for live cell gating for transfected cells. FSC-A = Forward Scatter Area, FSC-H = Forward Scatter Height, SSC-A = Side Scatter Area.

## REFERENCES

1. Jinek, M. et al. A programmable dual-RNA-guided DNA endonuclease in adaptive bacterial immunity. Science (80-.). 337, 816–821 (2012).

2. Cong, L. et al. Multiplex genome engineering using CRISPR/Cas systems. Science (80-.). 339, 819–823 (2013).

3. Shalem, O. et al. Genome-scale CRISPR-Cas9 knockout screening in human cells. Science 343, (2014).

4. Wang, T., Wei, J. J., Sabatini, D. M. & Lander, E. S. Genetic screens in human cells using the CRISPR-Cas9 system. Science (80-.). 343, 80–84 (2014).

5. Sanson, K. R. et al. Optimized libraries for CRISPR-Cas9 genetic screens with multiple modalities. Nat. Commun. 9, 5416 (2018).

6. Korkmaz, G. et al. Functional genetic screens for enhancer elements in the human genome using CRISPR-Cas9. Nat. Biotechnol. 34, 192 (2016).

7. Viswanatha, R., Li, Z., Hu, Y. & Perrimon, N. Pooled genome-wide CRISPR screening for basal and context-specific fitness gene essentiality in Drosophila cells. Elife 7, (2018).

8. Koike-Yusa, H., Li, Y., Tan, E. P., Velasco-Herrera, M. D. C. & Yusa, K. Genome-wide recessive genetic screening in mammalian cells with a lentiviral CRISPR-guide RNA library. Nat. Biotechnol. 32, 267–273 (2014).

9. Chen, B. et al. Dynamic imaging of genomic loci in living human cells by an optimized CRISPR/Cas system. Cell 155, 1479–1491 (2013).

10. Lane, A. B. et al. Enzymatically Generated CRISPR Libraries for Genome Labeling and Screening. Dev. Cell 34, 373–378 (2015).

11. Klann, T. S. et al. CRISPR-Cas9 epigenome editing enables high-throughput screening for functional regulatory elements in the human genome. Nat. Biotechnol. 35, 561–568 (2017).

12. Kosuri, S. & Church, G. M. Large-scale de novo DNA synthesis: Technologies and applications. Nature Methods vol. 11 499–507 (2014).

13. Arakawa, H. A method to convert mRNA into a gRNA library for CRISPR/Cas9 editing of any organism. Sci. Adv. 2, e1600699 (2016).

14. Cheng, J. et al. A Molecular Chipper technology for CRISPR sgRNA library generation and functional mapping of noncoding regions. Nat. Commun. 7, (2016).

15. Köferle, A. et al. CORALINA: a universal method for the generation of gRNA libraries for CRISPR-based screening. BMC Genomics 17, 917 (2016).

16. Morgan, R. D., Dwinell, E. A., Bhatia, T. K., Lang, E. M. & Luyten, Y. A. The MmeI family: type II restriction-modification enzymes that employ single-strand modification for host protection. Nucleic Acids Res. 37, 5208–21 (2009).

17. Nishimasu, H. et al. Crystal Structure of Cas9 in Complex with Guide RNA and Target DNA. Cell 156, 935–949 (2014).

18. Burger, A. et al. Maximizing mutagenesis with solubilized CRISPR-Cas9 ribonucleoprotein complexes. Development 143, 2025–37 (2016).

19. Pengpumkiat, S., Koesdjojo, M., Rowley, E. R., Mockler, T. C. & Remcho, V. T. Rapid synthesis of a long double-stranded oligonucleotide from a single-stranded nucleotide using magnetic beads and an oligo library. PLoS One 11, e0149774 (2016).

20. Mizushima, N., Yoshimori, T. & Levine, B. Methods in Mammalian Autophagy Research. Cell vol. 140 313–326 (2010).

21. Sanjana, N. E., Shalem, O. & Zhang, F. Improved vectors and genome-wide libraries for CRISPR screening. Nat. Methods 11, 783–784 (2014).

22. Jin, J. et al. An improved strategy for CRISPR/Cas9 gene knockout and subsequent wildtype and mutant gene rescue. PLoS One 15, 1–21 (2020).

23. Kim, H. K. et al. SpCas9 activity prediction by DeepSpCas9, a deep learning–based model with high generalization performance. Sci. Adv. 5, eaax9249 (2019).

24. Diatchenko, L. et al. Suppression subtractive hybridization: A method for generating differentially regulated or tissue-specific cDNA probes and libraries. Proc. Natl. Acad. Sci. U. S. A. 93, 6025–6030 (1996).

25. Hill, J. T. et al. Heart morphogenesis gene regulatory networks revealed by temporal expression analysis. Development 144, 3487–3498 (2017).

26. Jiang, W., Oikonomou, P. & Tavazoie, S. Comprehensive Genome-wide Perturbations via CRISPR Adaptation Reveal Complex Genetics of Antibiotic Sensitivity. Cell 180, 1002–1017.e31 (2020).

27. Dagdas, Y. S., Chen, J. S., Sternberg, S. H., Doudna, J. A. & Yildiz, A. A conformational checkpoint between DNA binding and cleavage by CRISPR-Cas9. Sci. Adv. 3, eaao0027 (2017).

28. Fu, Y., Sander, J. D., Reyon, D., Cascio, V. M. & Joung, J. K. Improving CRISPR-Cas nuclease specificity using truncated guide RNAs. Nat. Biotechnol. 32, 279–284 (2014).

29. Horlbeck, M. A. et al. Compact and highly active next-generation libraries for CRISPR-mediated gene repression and activation. Elife 5, (2016).

30. Gilbert, L. A. et al. Genome-Scale CRISPR-Mediated Control of Gene Repression and Activation. (2014) doi:10.1016/j.cell.2014.09.029.

31. Konermann, S. et al. Genome-scale transcriptional activation by an engineered CRISPR-Cas9 complex. Nature 517, 583–588 (2015).

32. Chen, S. et al. Genome-wide CRISPR screen in a mouse model of tumor growth and metastasis. Cell 160, 1246–1260 (2015).

33. Burns, C. G. & MacRae, C. A. Purification of hearts from zebrafish embryos. Biotechniques 40, 274–282 (2006).

34. Connelly, J. P. & Pruett-Miller, S. M. CRIS.py: A Versatile and High-throughput Analysis Program for CRISPR-based Genome Editing. Sci. Rep. 9, 1–8 (2019).

