## Supplemental Note 1: SLALOM Protocol for "SLALOM: A Simple and Rapid Method for Enzymatic Synthesis of CRISPR-Cas9 sgRNA Libraries"

SLALOM: A Rapid Method for Enzymatically Generating sgRNAs

**Capture bead preparation (100 pmol capture)**

Aliquot 50 µl of streptavidin magnetic beads into a 1.5 mL microcentrifuge tube.
Wash twice and Resuspend in 100 pmol biotinylated oligo.

| | |
|---|---|
| 17 µl | H20 |
| 2 µl | 10x cutsmart buffer |
| 1 µl | biotinylated oligo (100 µM) |

- Incubate at 20°C for 15 minutes.
- Add 30 µl of 1x cutsmart buffer.
- Wash twice and place in a new tube.

To wash beads: apply magnet, discard supernatant, resuspend in 50 µl of 1x cutsmart buffer by gently pipetting, apply magnet, discard supernatant, resuspend 50 µl of 1x cutsmart buffer.

**Adapter preparation (10 µM)**

| | |
|---|---|
| 35 µl | H20 |
| 5 ul | 10x cutsmart buffer |
| 5 ul | Top oligo (100 uM) |
| 5 ul | Bottom oligo (100 uM) |

- Hybridize oligos

| | |
|---|---|
| 98°C | 2:00 |
| 87°C - 67°C | 40:00 |
| 67°C - 27°C | 20:00 |
| 8°C | hold |

**PEG 6000 (50% w/v)**

| | |
|---|---|
| 25 g | PEG 6000 |
| 27 mL | Nuclease Free H20 (up to 50 mL) |

- heat at 50°C and stir with stir bar
- Filter sterilize

Notes:

- Drop dialysis can increase the efficiency of the first ligation step.
- Use fresh SAM.

**1. HpaII digestion, combine components in a clean 0.2 ml PCR tube:**

| | |
|---|---|
| x µl | H20 (up to 50 µl) |
| x µl | Input DNA (> 20 pmol cut sites) |
| 5 µl | 10x cutsmart buffer |
| 1 µl | HpaII |

Note:
If the number of HpaII sites in a DNA sample can be calculated it is ideal, if not the amount of input DNA required for genomic DNA is usually in the range of 1-2 µg.

- Incubate at 37°C for 20 min, 80°C for 20 min

**2. Scaffold adapter ligation, add the following components to the tube:**

| | |
|---|---|
| 2 µl | 10x cutsmart buffer |
| 1 µl | 10 µM scaffold adapter (10 pmol) |
| 7 µl | 10 mM ATP |
| 1 µl | T4 DNA Ligase (2,000,000 units/mL) |
| 10 µl | 50% PEG 6000 |

- Incubate at room temperature for 30 minutes.
- Resuspend capture beads in the previous solution (about 70 µl).
- Incubate at room temperature for 15 minutes.
- Wash the beads twice with 1x cutsmart buffer and place in a new tube.

**3. MmeI digestion, resuspend beads in:**

| | |
|---|---|
| 43 µl | H20 |
| 5 µl | 10x Cutsmart buffer |
| 1 µl | 2.5 mM SAM |
| 1 µl | MmeI |

- Incubate at 37°C for 30 minutes.
- Wash twice and place in a new tube.

**4. T7 adapter ligation, resuspend beads in:**

| | |
|---|---|
| 29 µl | H20 |
| 5 µl | 10x Cutsmart buffer |
| 3 µl | 10 uM T7 adapter (30 pmol) |
| 5 µl | 10 mM ATP |
| 1 µl | T4 Ligase (2,000,000 units/mL) |
| 7 µl | 50% PEG 6000 |

- Incubate at room temperature for 30 minutes.
- Wash twice and place in a new tube.

**5. Bst elution and nick repair, resuspend beads in:**

| | |
|---|---|
| 43 µl | H20 |
| 5 µl | 10x Cutsmart buffer |
| 1 µl | 10 mM dNTPs |
| 1 µl | DNA polymerase I (10,000 units/mL) |

- Incubate at room temperature for 10 minutes.
- Collect solution and purify using a PCR purification column.

- PCR amplification can be accomplished using a standard Phusion 2 step PCR with 50 ng of library as a template - [98°C, 64°C for 10 seconds each] x 30
