## Supplemental Table 1 for "SLALOM: A Simple and Rapid Method for Enzymatic Synthesis of CRISPR-Cas9 sgRNA Libraries"

Supplemental Table 1: SLALOM oligos

**Anneal and Extend**

>scaffold_wt_R
AAAAGCACCGACTCGGTGCCACTTTTTCAAGTTGATAACGGACTAGCCTTATTTTAACTTGCTATTTCTAGCTCTAAAAC

>t7_GFP20_wt_F
GAAATTAATACGACTCACTATAGGGCGAGGAGCTGTTCACCGGTTTTAGAGCTAGAAATAGCAAGTTAAAATAAGGCTAGTCCG

>t7_GFP18_wt_F
GAAATTAATACGACTCACTATAGCGAGGAGCTGTTCACCGGTTTTAGAGCTAGAAATAGCAAGTTAAAATAAGGCTAGTCCG

>scaffold_mod4_R
AAAAGCACCGACTCGGTGCCACTTTTTCAAGTTGATAACGGACTAGCCTTATTGGAACTTGCTATTTCTAGCTCTCCAAC

>t7_GFP20_mod4_F
GAAATTAATACGACTCACTATAGGGCGAGGAGCTGTTCACCGGTTGGAGAGCTAGAAATAGCAAGTTCCAATAAGGC

>t7_GFP18_mod4_F
GAAATTAATACGACTCACTATAGCGAGGAGCTGTTCACCGGTTGGAGAGCTAGAAATAGCAAGTTCCAATAAGGC

>scaffold_mod7_R
AAAAGCACCGACTCGGTGCCACTTTTTCAAGTTGATAACGGACTAGCCTTAGGTTGACTTGCTATTTCTAGCTCCAACCG

>t7_GFP20_mod7_F
GAAATTAATACGACTCACTATAGGGCGAGGAGCTGTTCACCGCGGTTGGAGCTAGAAATAGCAAGTCAACCTAAG

>t7_GFP20_mod7_R
GAAATTAATACGACTCACTATAGCGAGGAGCTGTTCACCGCGGTTGGAGCTAGAAATAGCAAGTCAACCTAAG

**1000 bp cmlc:GFP**

>cmlc_600
GACATTTTTCCTGCCATCCTGTCTTAGGCTG

>GFP_400
TCCAGCTTGTGCCCCAGGATGTTG

**E coli & Zebrafish Libraries**

>T7_promoter_adapter_top_eco_zeb
CGGCGAAATTAATACGACTCACTATAGNN

>T7_promoter_adapter_bottom_eco_zeb
CTATAGTGAGTCGTATTAATTTCGCCG

>Scaffold_adapter_top_eco_zeb
CGGTTGGAGCTAGAAATAGCAAGTCAACCTAAGGCTAGTCCGTTATCAACTTGAAAAAGTGGCAC

>Scaffold_adapter_bottom_eco_zeb
AAAAAAAGCACCGACTCGGTGCCACTTTTTCAAGTTGATAACGGACTAGCCTTAGGTTGACTTGCTATTTCTAGCTCCAAC

>Capture_oligo_eco_zeb
CGAGTCGGTGCTTTTTTTGCGCATGC/3BioTEG/

>PCR_primer_Forward_eco_zeb
CGGCGAAATTAATACGACTCACTATAG

>PCR_primer_Reverse_eco_zeb
AAAAAAAGCACCGACTCGG

**GFP Library**

>U6_promoter_adapter_top_lentiCRISPRv2
GCGAAATTAATACGACTCACTATAGCGTCTCACACCGNN

>U6_promoter_adapter_bottom_lentiCRISPRv2
CGGTGTGAGACGCTATAGTGAGTCGTATTAATTTCGC

>Scaffold_adapter_top_lentiCRISPRv2
CGGTTGGAGCTATGCTGGAAACAGCATAGCAAGTCAACCTAAGGCTAGTCCGTTATCAACTTGAAAAAGTGGCACCGAGTCGGTGC
TTTTTTTG

>Scaffold_adapter_bottom_lentiCRISPRv2
GACCTACGTCTCGCTAGCAAAAAAAGCACCGACTCGGTGCCACTTTTTCAAGTTGATAACGGACTAGCCTTAGGTTGACTTGCTAT
GCTGTTTCCAGCATAGCTCCAAC

>Capture_oligo_lentiCRISPRv2
CTAGCGAGACGTAGGTCGCGCATGC/3BioTEG/

>PCR_primer_Forward_lentiCRISPRv2
GCGAAATTAATACGACTCACTATAGCGTCTCAC

>PCR_primer_Reverse_lentiCRISPRv2
GACCTACGTCTCGCTAGCAAAAAAAGCAC
