## Supplemental Note 2 for "SLALOM: A Simple and Rapid Method for Enzymatic Synthesis of CRISPR-Cas9 sgRNA Libraries": SupplementalNote2.html

Analysis of E. coli Library


Code 

- Show All Code
- Hide All Code
- Download Rmd

### Analysis of E. coli Library

###### Jonathon Hill

We have generated and sequenced a SLOLAM library targeting the E. coli genome. This will show us the ability of the method to create complex libraries and is comparable to the Lane et al. method for enzymatic sgRNA library generation, as they also made a library to E. coli. This analysis will build on Josh’s code that pulled the sequences and lengths of the spacers in the library.

#### Setup


```
library(Rsubread)

# Make genome index
buildindex("MG1655", './MG1655_genome/MG1655_complete_genome.fasta')
```

#### Align


```
library(Rsubread)
library(Rsamtools)
align(
  index = 'MG1655_genome/MG1655',
  readfile1 = 'fastq/e_coli_captured_sequence.fastq',
  output_file = 'bam/e_coli_captured_sequence.BAM',
  type = "dna",
  indels = 0,
  maxMismatches = 2,
  unique = TRUE,
  TH1 = 1,
  nthreads = 8
)
```


```
        ==========     _____ _    _ ____  _____  ______          _____  
        =====         / ____| |  | |  _ \|  __ \|  ____|   /\   |  __ \ 
          =====      | (___ | |  | | |_) | |__) | |__     /  \  | |  | |
            ====      \___ \| |  | |  _ <|  _  /|  __|   / /\ \ | |  | |
              ====    ____) | |__| | |_) | | \ \| |____ / ____ \| |__| |
        ==========   |_____/ \____/|____/|_|  \_\______/_/    \_\_____/
       Rsubread 2.0.1

//================================= setting ==================================\\
||                                                                            ||
|| Function      : Read alignment (DNA-Seq)                                   ||
|| Input file    : e_coli_captured_sequence.fastq                             ||
|| Output file   : e_coli_captured_sequence.BAM (BAM)                         ||
|| Index name    : MG1655                                                     ||
||                                                                            ||
||                    ------------------------------------                    ||
||                                                                            ||
||                               Threads : 8                                  ||
||                          Phred offset : 33                                 ||
||                             Min votes : 1 / 10                             ||
||                        Max mismatches : 2                                  ||
||                      Max indel length : 0                                  ||
||            Report multi-mapping reads : no                                 ||
|| Max alignments per multi-mapping read : 1                                  ||
||                                                                            ||
\\============================================================================//

//================ Running (23-Jun-2020 16:37:53, pid=24540) =================\\
||                                                                            ||
|| Check the input reads.                                                     ||
|| The input file contains base space reads.                                  ||
|| Initialise the memory objects.                                             ||
|| Estimate the mean read length.                                             ||
|| The range of Phred scores observed in the data is [14,38]                  ||
|| Create the output BAM file.                                                ||
|| Check the index.                                                           ||
|| Init the voting space.                                                     ||
|| Global environment is initialised.                                         ||
|| Load the 1-th index block...                                               ||
|| The index block has been loaded.                                           ||
|| Start read mapping in chunk.                                               ||
||    2% completed, 0.0 mins elapsed, rate=109.8k reads per second            ||
||    8% completed, 0.0 mins elapsed, rate=106.3k reads per second            ||
||   15% completed, 0.0 mins elapsed, rate=108.7k reads per second            ||
||   22% completed, 0.1 mins elapsed, rate=107.3k reads per second            ||
||   28% completed, 0.1 mins elapsed, rate=105.8k reads per second            ||
||   35% completed, 0.1 mins elapsed, rate=106.7k reads per second            ||
||   42% completed, 0.1 mins elapsed, rate=106.3k reads per second            ||
||   48% completed, 0.1 mins elapsed, rate=105.0k reads per second            ||
||   55% completed, 0.1 mins elapsed, rate=104.3k reads per second            ||
||   62% completed, 0.1 mins elapsed, rate=104.0k reads per second            ||
||   70% completed, 0.2 mins elapsed, rate=93.7k reads per second             ||
||   73% completed, 0.2 mins elapsed, rate=91.1k reads per second             ||
||   76% completed, 0.2 mins elapsed, rate=88.8k reads per second             ||
||   80% completed, 0.2 mins elapsed, rate=87.0k reads per second             ||
||   83% completed, 0.2 mins elapsed, rate=85.5k reads per second             ||
||   86% completed, 0.2 mins elapsed, rate=84.1k reads per second             ||
||   90% completed, 0.3 mins elapsed, rate=82.8k reads per second             ||
||   93% completed, 0.3 mins elapsed, rate=81.7k reads per second             ||
||   96% completed, 0.3 mins elapsed, rate=80.5k reads per second             ||
||                                                                            ||
||                           Completed successfully.                          ||
||                                                                            ||
\\====================================    ====================================//

//================================   Summary =================================\\
||                                                                            ||
||                 Total reads : 1,394,595                                    ||
||                      Mapped : 1,106,316 (79.3%)                            ||
||             Uniquely mapped : 1,106,316                                    ||
||               Multi-mapping : 0                                            ||
||                                                                            ||
||                    Unmapped : 288,279                                      ||
||                                                                            ||
||                      Indels : 0                                            ||
||                                                                            ||
||                Running time : 0.3 minutes                                  ||
||                                                                            ||
\\============================================================================//

                      e.coli.captured.sequence.BAM
Total_reads                                1394595
Mapped_reads                               1106316
Uniquely_mapped_reads                      1106316
Multi_mapping_reads                              0
Unmapped_reads                              288279
Indels                                           0
```


```
sortBam(
  file = 'bam/e_coli_captured_sequence.BAM',
  destination = 'bam/e_coli_captured_sequence.sorted'
)
```


```
[1] "bam/e_coli_captured_sequence.sorted.bam"
```


```
indexBam('bam/e_coli_captured_sequence.sorted.bam')
```


```
      bam/e_coli_captured_sequence.sorted.bam 
"bam/e_coli_captured_sequence.sorted.bam.bai"
```

#### Assign Features

We will borrow the Rsubread package featureCounts() function to assign reads to HpaII sites. All sites in the E. coli genome are compiled in a fake gtf file.


```
library(Rsubread)
counts = featureCounts(
  files = 'bam/e_coli_captured_sequence.sorted.bam',
  annot.ext = 'e_coli_HpaII.GTF',
  GTF.featureType = "HpaII",
  minMQS = 20,
  primaryOnly = TRUE,
  ignoreDup = TRUE,
  isGTFAnnotation = TRUE,
  nthreads = 8)
```


```
        ==========     _____ _    _ ____  _____  ______          _____  
        =====         / ____| |  | |  _ \|  __ \|  ____|   /\   |  __ \ 
          =====      | (___ | |  | | |_) | |__) | |__     /  \  | |  | |
            ====      \___ \| |  | |  _ <|  _  /|  __|   / /\ \ | |  | |
              ====    ____) | |__| | |_) | | \ \| |____ / ____ \| |__| |
        ==========   |_____/ \____/|____/|_|  \_\______/_/    \_\_____/
       Rsubread 2.0.1

//========================== featureCounts setting ===========================\\
||                                                                            ||
||             Input files : 1 BAM file                                       ||
||                           o e_coli_captured_sequence.sorted.bam            ||
||                                                                            ||
||              Annotation : e_coli_HpaII.GTF (GTF)                           ||
||      Dir for temp files : .                                                ||
||                 Threads : 8                                                ||
||                   Level : meta-feature level                               ||
||              Paired-end : no                                               ||
||      Multimapping reads : counted                                          ||
||     Multiple alignments : primary alignment only                           ||
|| Multi-overlapping reads : not counted                                      ||
||   Min overlapping bases : 1                                                ||
||        Duplicated Reads : ignored                                          ||
||                                                                            ||
\\============================================================================//

//================================= Running ==================================\\
||                                                                            ||
|| Load annotation file e_coli_HpaII.GTF ...                                  ||
||    Features : 24311                                                        ||
||    Meta-features : 24311                                                   ||
||    Chromosomes/contigs : 1                                                 ||
||                                                                            ||
|| Process BAM file e_coli_captured_sequence.sorted.bam...                    ||
||    Single-end reads are included.                                          ||
||    Total alignments : 1394595                                              ||
||    Successfully assigned alignments : 1084767 (77.8%)                      ||
||    Running time : 0.01 minutes                                             ||
||                                                                            ||
|| Write the final count table.                                               ||
|| Write the read assignment summary.                                         ||
||                                                                            ||
\\============================================================================//
```


```
counts$stat
```


```
                          Status e.coli.captured.sequence.sorted.bam
1                       Assigned                             1084767
2            Unassigned_Unmapped                              288279
3           Unassigned_Read_Type                                   0
4           Unassigned_Singleton                                   0
5      Unassigned_MappingQuality                                 726
6             Unassigned_Chimera                                   0
7      Unassigned_FragmentLength                                   0
8           Unassigned_Duplicate                                   0
9        Unassigned_MultiMapping                                   0
10          Unassigned_Secondary                                   0
11           Unassigned_NonSplit                                   0
12         Unassigned_NoFeatures                               18781
13 Unassigned_Overlapping_Length                                   0
14          Unassigned_Ambiguity                                2042
```


```
# turn genecounts into dataframe, add log2
genecounts = counts$counts
genecounts = data.frame(feature = rownames(genecounts), counts = genecounts[TRUE])
genecounts$log2 = log2(genecounts$counts)
head(genecounts)
```


```
    feature counts     log2
1 HpaII_583    109 6.768184
2 HpaII_679     93 6.539159
3 HpaII_740     22 4.459432
4 HpaII_784     63 5.977280
5 HpaII_874     11 3.459432
6 HpaII_911      0     -Inf
```


```
# remove 0
genecounts = subset(genecounts, genecounts$counts > 0)
```

#### Plot Results


```
library(ggplot2)
color = "#E69F00"
color = "#4186C4"
# histogram of coverage
ggplot(genecounts, aes(x = log2) ) +
  geom_histogram(
    binwidth = 0.4,
    color = color,
    fill = color,
    alpha = 0.2) +
  xlab('log2 Counts per Target Site') +
  ylab('Number of Predicted Target Sites')
```


```
# histogram of spacer length
targeter_length = read.csv(file = 'e_coli_length.csv')
head(targeter_length)
```


```
  count length
1 count     25
2 count     25
3 count     23
4 count     23
5 count     22
6 count     22
```


```
ggplot(targeter_length, aes(x = length) ) +
  geom_histogram(
    binwidth = 1,
    color = color,
    fill = color,
    alpha = 0.2) +
  xlab('Length of Targeting Region (bp)') +
  ylab('Library Reads')
```

### Identify Site Coverage and Library Purity


```
library(GenomicAlignments)
library(GenomicFeatures)
library(GenomicRanges)
gal <- readGAlignments("bam/e_coli_captured_sequence.sorted.bam") 
hpaII_sites <- read.table("e_coli_HpaII.GTF")
hpaII_pos <- GRanges(seqnames = hpaII_sites$V1, 
                     ranges = IRanges(start = hpaII_sites$V4, end = hpaII_sites$V5))
unmatched_reads <- subsetByOverlaps(gal,hpaII_pos, ignore.strand = TRUE, invert = TRUE)
unmatched_hpaII <- subsetByOverlaps(hpaII_pos, gal, ignore.strand = TRUE, invert = TRUE)
unmatched_unique_reads <- reduce(GRanges(unmatched_reads), ignore.strand = TRUE)
df <- data.frame(group = c("Percent of sites targeted", "Percent not targeted"), value = c(sum(counts$counts > 0) / nrow(counts$counts), 1 - sum(counts$counts > 0) / nrow(counts$counts)))
df
```


```
                      group     value
1 Percent of sites targeted 0.9164987
2      Percent not targeted 0.0835013
```


```
cat("\nFraction of reads not at a valid site: ", 
length(unmatched_reads)/(sum(counts$counts) + length(unmatched_reads)))
```


```
Fraction of reads not at a valid site:  0.01716035
```


#### Session Information


```
sessionInfo()
```
