## Supplemental Note 3 for "SLALOM: A Simple and Rapid Method for Enzymatic Synthesis of CRISPR-Cas9 sgRNA Libraries": SupplementalNote3.html

Heart Transcriptome Library Analysis


### Heart Transcriptome Library Analysis

###### Jonathon Hill

#### Setup

```
library(Rsubread)
library(Rsamtools)
```

```
## Loading required package: GenomeInfoDb
```

```
## Loading required package: BiocGenerics
```

```
## Loading required package: parallel
```

```
## 
## Attaching package: 'BiocGenerics'
```

```
## The following objects are masked from 'package:parallel':
## 
##     clusterApply, clusterApplyLB, clusterCall, clusterEvalQ,
##     clusterExport, clusterMap, parApply, parCapply, parLapply,
##     parLapplyLB, parRapply, parSapply, parSapplyLB
```

```
## The following objects are masked from 'package:stats':
## 
##     IQR, mad, sd, var, xtabs
```

```
## The following objects are masked from 'package:base':
## 
##     anyDuplicated, append, as.data.frame, basename, cbind, colnames,
##     dirname, do.call, duplicated, eval, evalq, Filter, Find, get, grep,
##     grepl, intersect, is.unsorted, lapply, Map, mapply, match, mget,
##     order, paste, pmax, pmax.int, pmin, pmin.int, Position, rank,
##     rbind, Reduce, rownames, sapply, setdiff, sort, table, tapply,
##     union, unique, unsplit, which, which.max, which.min
```

```
## Loading required package: S4Vectors
```

```
## Loading required package: stats4
```

```
## 
## Attaching package: 'S4Vectors'
```

```
## The following object is masked from 'package:base':
## 
##     expand.grid
```

```
## Loading required package: IRanges
```

```
## Loading required package: GenomicRanges
```

```
## Loading required package: Biostrings
```

```
## Loading required package: XVector
```

```
## 
## Attaching package: 'Biostrings'
```

```
## The following object is masked from 'package:base':
## 
##     strsplit
```

```
library(VennDiagram)
```

```
## Loading required package: grid
```

```
## Loading required package: futile.logger
```

```
library(ggplot2)

reference_genome = "GRCz11"
gtf_file = "/home/jth54/GRCz11/Danio_rerio.GRCz11.93.chr.gtf"
```

#### Spacer Length

```
data <- readLines("Heart/fastq/fish_heart_captured_sequence.fastq")
seqs <- data[seq(2, length(data), 4)]

ggplot(data.frame(seqs = nchar(seqs)), aes(x = seqs) ) +
  geom_histogram(
    binwidth = 1,
    color = "royalblue4",
    fill = "royalblue4",
    alpha = .3) +
  xlab('Spacer Length') +
  ylab('Frequency') +
  scale_x_continuous(limits = c(1,25)) +
  theme(text = element_text(size = 24))
```

```
## Warning: Removed 2 rows containing missing values (geom_bar).
```

#### Align

```
subjunc(index = reference_genome,
        readfile1 = 'Heart/fastq/fish_heart_captured_sequence.fastq',
        output_file = 'Heart/bam/fish_heart_captured_sequence.BAM',
        nthreads = 8,
        TH1 = 1,
        useAnnotation = TRUE,
        annot.ext = gtf_file,
        isGTF = TRUE)
```

```
## 
##         ==========     _____ _    _ ____  _____  ______          _____  
##         =====         / ____| |  | |  _ \|  __ \|  ____|   /\   |  __ \ 
##           =====      | (___ | |  | | |_) | |__) | |__     /  \  | |  | |
##             ====      \___ \| |  | |  _ <|  _  /|  __|   / /\ \ | |  | |
##               ====    ____) | |__| | |_) | | \ \| |____ / ____ \| |__| |
##         ==========   |_____/ \____/|____/|_|  \_\______/_/    \_\_____/
##        Rsubread 2.0.1
## 
## //================================= setting ==================================\\
## ||                                                                            ||
## || Function      : Read alignment + Junction detection (RNA-Seq)              ||
## || Input file    : fish_heart_captured_sequence.fastq                         ||
## || Output file   : fish_heart_captured_sequence.BAM (BAM)                     ||
## || Index name    : GRCz11                                                     ||
## ||                                                                            ||
## ||                    ------------------------------------                    ||
## ||                                                                            ||
## ||                               Threads : 8                                  ||
## ||                          Phred offset : 33                                 ||
## ||                             Min votes : 1 / 14                             ||
## ||                        Max mismatches : 3                                  ||
## ||                      Max indel length : 5                                  ||
## ||            Report multi-mapping reads : yes                                ||
## || Max alignments per multi-mapping read : 1                                  ||
## ||                           Annotations : Danio_rerio.GRCz11.93.chr.gtf  ... ||
## ||                                                                            ||
## \\============================================================================//
## 
## //================ Running (24-Jun-2020 10:57:44, pid=12013) =================\\
## ||                                                                            ||
## || Check the input reads.                                                     ||
## || The input file contains base space reads.                                  ||
## || Initialise the memory objects.                                             ||
## || Estimate the mean read length.                                             ||
## || The range of Phred scores observed in the data is [14,38]                  ||
## || Create the output BAM file.                                                ||
## || Check the index.                                                           ||
## || Init the voting space.                                                     ||
## || Load the annotation file.                                                  ||
## || 481079 annotation records were loaded.                                     ||
## ||                                                                            ||
## || Chromosomes/contigs in index but not in annotation :                       ||
## ||    KN150665.1                                                              ||
## ||    KN150549.1                                                              ||
## ||    KN149996.1                                                              ||
## ||    KZ116059.1                                                              ||
## ||    KN150350.1                                                              ||
## ||    KN150403.1                                                              ||
## ||    KN150181.1                                                              ||
## ||    KN150234.1                                                              ||
## ||    KN149850.1                                                              ||
## ||    KN149903.1                                                              ||
## ||    KN150065.1                                                              ||
## ||    KN150118.1                                                              ||
## ||    KZ115966.1                                                              ||
## ||    KN150593.1                                                              ||
## ||    KN150646.1                                                              ||
## ||    KN148828.2                                                              ||
## ||    KN150477.1                                                              ||
## ||    KN150500.1                                                              ||
## ||    KN149977.2                                                              ||
## ||    KZ116010.1                                                              ||
## ||    KN150331.1                                                              ||
## ||    KN150162.1                                                              ||
## ||    KN150215.1                                                              ||
## ||    KN149715.1                                                              ||
## ||    KN150690.1                                                              ||
## ||    KN150574.1                                                              ||
## ||    KN150458.1                                                              ||
## ||    KN150289.1                                                              ||
## ||    KN149958.1                                                              ||
## ||    KN150312.1                                                              ||
## ||    KN149789.1                                                              ||
## ||    KN150090.1                                                              ||
## ||    KN149812.1                                                              ||
## ||    KN147651.2                                                              ||
## ||    KZ115991.1                                                              ||
## ||    KN150671.1                                                              ||
## ||    KN150555.1                                                              ||
## ||    KN150608.1                                                              ||
## ||    KN150386.1                                                              ||
## ||    KZ116065.1                                                              ||
## ||    KN150439.1                                                              ||
## ||    KN149939.1                                                              ||
## ||    KN150240.1                                                              ||
## ||    KN150071.1                                                              ||
## ||    KN149740.1                                                              ||
## ||    KN150008.1                                                              ||
## ||    KN147632.2                                                              ||
## ||    KZ115972.1                                                              ||
## ||    KN150652.1                                                              ||
## ||    KN150483.1                                                              ||
## ||    KN150536.1                                                              ||
## ||    KZ116046.1                                                              ||
## ||    KN149983.1                                                              ||
## ||    KN149867.1                                                              ||
## ||    KN150221.1                                                              ||
## ||    KN149698.1                                                              ||
## ||    KN150052.1                                                              ||
## ||    KN149721.1                                                              ||
## ||    KN150105.1                                                              ||
## ||    KZ115953.1                                                              ||
## ||    KN150580.1                                                              ||
## ||    KN150633.1                                                              ||
## ||    KN150464.1                                                              ||
## ||    KN150517.1                                                              ||
## ||    KZ116027.1                                                              ||
## ||    KN149795.1                                                              ||
## ||    KN150179.1                                                              ||
## ||    KN149848.1                                                              ||
## ||    KN150202.1                                                              ||
## ||    KN150033.1                                                              ||
## ||    KN149702.1                                                              ||
## ||    KN150561.1                                                              ||
## ||    KN150614.1                                                              ||
## ||    KN150392.1                                                              ||
## ||    KN150445.1                                                              ||
## ||    KN150276.1                                                              ||
## ||    KN149892.1                                                              ||
## ||    KN150329.1                                                              ||
## ||    KZ116008.1                                                              ||
## ||    KN149829.1                                                              ||
## ||    KN150130.1                                                              ||
## ||    KN150014.1                                                              ||
## ||    KN150711.1                                                              ||
## ||    KN150542.1                                                              ||
## ||    KZ116052.1                                                              ||
## ||    KN150426.1                                                              ||
## ||    KN149873.1                                                              ||
## ||    KN150257.1                                                              ||
## ||    KN149926.1                                                              ||
## ||    KN149757.1                                                              ||
## ||    KN150111.1                                                              ||
## ||    KZ115989.1                                                              ||
## ||    KN150669.1                                                              ||
## ||    KN150470.1                                                              ||
## ||    KN150523.1                                                              ||
## ||    KN150354.1                                                              ||
## ||    KN149970.1                                                              ||
## ||    KZ116033.1                                                              ||
## ||    KN150407.1                                                              ||
## ||    KN150185.1                                                              ||
## ||    KN149854.1                                                              ||
## ||    KN150238.1                                                              ||
## ||    KN149907.1                                                              ||
## ||    KN149685.1                                                              ||
## ||    KN149738.1                                                              ||
## ||    KN150620.1                                                              ||
## ||    KN150451.1                                                              ||
## ||    KN150504.1                                                              ||
## ||    KN150282.1                                                              ||
## ||    KZ116014.1                                                              ||
## ||    KN149951.1                                                              ||
## ||    KN150335.1                                                              ||
## ||    KN149782.1                                                              ||
## ||    KN150166.1                                                              ||
## ||    KN150219.1                                                              ||
## ||    KN149835.1                                                              ||
## ||    KN149719.1                                                              ||
## ||    KN150020.1                                                              ||
## ||    KN150578.1                                                              ||
## ||    KN150601.1                                                              ||
## ||    KN150432.1                                                              ||
## ||    KN150263.1                                                              ||
## ||    KN149932.1                                                              ||
## ||    KN150094.1                                                              ||
## ||    KN150147.1                                                              ||
## ||    KN149763.1                                                              ||
## ||    KN149816.1                                                              ||
## ||    KZ115995.1                                                              ||
## ||    KN150675.1                                                              ||
## ||    KN150559.1                                                              ||
## ||    KN150360.1                                                              ||
## ||    KN150413.1                                                              ||
## ||    KN150191.1                                                              ||
## ||    KN149860.1                                                              ||
## ||    KN150244.1                                                              ||
## ||    KN149913.1                                                              ||
## ||    KN149691.1                                                              ||
## ||    KN150128.1                                                              ||
## ||    KN147636.1                                                              ||
## ||    KZ115976.1                                                              ||
## ||    KN150656.2                                                              ||
## ||    KN150709.1                                                              ||
## ||    KN150487.1                                                              ||
## ||    KN150510.1                                                              ||
## ||    KZ116020.1                                                              ||
## ||    KN150341.1                                                              ||
## ||    KN150172.1                                                              ||
## ||    KN149841.1                                                              ||
## ||    KN150225.1                                                              ||
## ||    KN150056.1                                                              ||
## ||    KN149725.1                                                              ||
## ||    KZ115957.1                                                              ||
## ||    KN150468.1                                                              ||
## ||    KN150299.1                                                              ||
## ||    KN149968.1                                                              ||
## ||    KZ116001.1                                                              ||
## ||    KN150322.1                                                              ||
## ||    KN150206.1                                                              ||
## ||    KN149822.1                                                              ||
## ||    KN150037.1                                                              ||
## ||    KN149706.1                                                              ||
## ||    KN150565.1                                                              ||
## ||    KN150396.1                                                              ||
## ||    KN150449.1                                                              ||
## ||    KN149896.1                                                              ||
## ||    KN149949.1                                                              ||
## ||    KN150250.1                                                              ||
## ||    KN150303.1                                                              ||
## ||    KN150081.1                                                              ||
## ||    KN149750.1                                                              ||
## ||    KN150134.1                                                              ||
## ||    KN149803.1                                                              ||
## ||    KN150018.1                                                              ||
## ||    KN147642.2                                                              ||
## ||    KZ115982.1                                                              ||
## ||    KN150662.1                                                              ||
## ||    KN150493.1                                                              ||
## ||    KN150546.1                                                              ||
## ||    KZ116056.1                                                              ||
## ||    KN149993.1                                                              ||
## ||    KN150400.1                                                              ||
## ||    KN150231.1                                                              ||
## ||    KN149900.1                                                              ||
## ||    KN150062.1                                                              ||
## ||    KN150115.1                                                              ||
## ||    KZ115963.1                                                              ||
## ||    KN150643.1                                                              ||
## ||    KN150527.1                                                              ||
## ||    KN150358.1                                                              ||
## ||    KZ116037.1                                                              ||
## ||    KN149974.1                                                              ||
## ||    KN150189.1                                                              ||
## ||    KN150212.1                                                              ||
## ||    KN149689.2                                                              ||
## ||    KN150043.1                                                              ||
## ||    KN150571.1                                                              ||
## ||    KN150455.1                                                              ||
## ||    KN150508.1                                                              ||
## ||    KN150286.1                                                              ||
## ||    KZ116018.1                                                              ||
## ||    KN149955.1                                                              ||
## ||    KN149786.1                                                              ||
## ||    KN149839.1                                                              ||
## ||    KN150140.2                                                              ||
## ||    KN150024.1                                                              ||
## ||    KN150698.1                                                              ||
## ||    KN150552.1                                                              ||
## ||    KN150605.1                                                              ||
## ||    KZ116062.1                                                              ||
## ||    KN150383.1                                                              ||
## ||    KN150436.1                                                              ||
## ||    KN149883.1                                                              ||
## ||    KN150267.1                                                              ||
## ||    KN149936.1                                                              ||
## ||    KN150098.1                                                              ||
## ||    KN149767.1                                                              ||
## ||    KN150121.1                                                              ||
## ||    KN150005.1                                                              ||
## ||    KZ115999.1                                                              ||
## ||    KN150679.1                                                              ||
## ||    KN149980.1                                                              ||
## ||    KN150364.1                                                              ||
## ||    KZ116043.1                                                              ||
## ||    KN150195.1                                                              ||
## ||    KN149864.1                                                              ||
## ||    KN150248.1                                                              ||
## ||    KN149917.1                                                              ||
## ||    KN150079.1                                                              ||
## ||    KN149695.1                                                              ||
## ||    KN149748.1                                                              ||
## ||    KN150102.1                                                              ||
## ||    KZ115950.1                                                              ||
## ||    KN150630.1                                                              ||
## ||    KN150514.1                                                              ||
## ||    KN150292.1                                                              ||
## ||    KN150345.1                                                              ||
## ||    KN149961.1                                                              ||
## ||    KZ116024.1                                                              ||
## ||    KN149792.1                                                              ||
## ||    KN150176.1                                                              ||
## ||    KN150229.1                                                              ||
## ||    KN149729.1                                                              ||
## ||    KN150030.1                                                              ||
## ||    KN150588.1                                                              ||
## ||    KZ116005.1                                                              ||
## ||    KN150326.1                                                              ||
## ||    KN150157.1                                                              ||
## ||    KN149826.1                                                              ||
## ||    KN150011.1                                                              ||
## ||    KN150685.1                                                              ||
## ||    KN150254.1                                                              ||
## ||    KN150307.1                                                              ||
## ||    KN149923.1                                                              ||
## ||    KN150085.1                                                              ||
## ||    KN149754.1                                                              ||
## ||    KN149807.1                                                              ||
## ||    KZ115986.1                                                              ||
## ||    KN150666.1                                                              ||
## ||    KN150520.1                                                              ||
## ||    KN149997.1                                                              ||
## ||    KZ116030.1                                                              ||
## ||    KN150351.1                                                              ||
## ||    KN150404.1                                                              ||
## ||    KN150182.1                                                              ||
## ||    KN149851.1                                                              ||
## ||    KN149904.1                                                              ||
## ||    KN150066.1                                                              ||
## ||    KN149682.1                                                              ||
## ||    KN150119.1                                                              ||
## ||    KN149735.1                                                              ||
## ||    KZ115967.1                                                              ||
## ||    KN150594.1                                                              ||
## ||    KN150647.1                                                              ||
## ||    KN150478.1                                                              ||
## ||    KN150501.1                                                              ||
## ||    KN149978.1                                                              ||
## ||    KN150332.1                                                              ||
## ||    KZ116011.1                                                              ||
## ||    KN150216.1                                                              ||
## ||    KN149832.1                                                              ||
## ||    KN150047.1                                                              ||
## ||    KN149716.1                                                              ||
## ||    KN150691.1                                                              ||
## ||    KZ115948.1                                                              ||
## ||    KN150575.1                                                              ||
## ||    KN150628.1                                                              ||
## ||    KN150459.1                                                              ||
## ||    KN149959.1                                                              ||
## ||    KN150260.1                                                              ||
## ||    KN150313.1                                                              ||
## ||    KN150144.1                                                              ||
## ||    KN149760.1                                                              ||
## ||    KN149813.1                                                              ||
## ||    KN150028.1                                                              ||
## ||    KN147652.2                                                              ||
## ||    KZ115992.1                                                              ||
## ||    KN150556.1                                                              ||
## ||    KN150609.1                                                              ||
## ||    KN150387.1                                                              ||
## ||    KZ116066.1                                                              ||
## ||    KN149887.1                                                              ||
## ||    KN150241.1                                                              ||
## ||    KN150072.1                                                              ||
## ||    KN149741.1                                                              ||
## ||    KN150125.1                                                              ||
## ||    KN150009.1                                                              ||
## ||    KZ115973.1                                                              ||
## ||    KN150653.1                                                              ||
## ||    KN150484.1                                                              ||
## ||    KN150537.1                                                              ||
## ||    KZ116047.1                                                              ||
## ||    KN150368.1                                                              ||
## ||    KN149984.1                                                              ||
## ||    KN150199.1                                                              ||
## ||    KN150222.1                                                              ||
## ||    KN149722.1                                                              ||
## ||    KN150106.1                                                              ||
## ||    KZ115954.1                                                              ||
## ||    KN150581.1                                                              ||
## ||    KN150634.1                                                              ||
## ||    KN150465.1                                                              ||
## ||    KN150518.1                                                              ||
## ||    KN150296.1                                                              ||
## ||    KZ116028.1                                                              ||
## ||    KN149965.1                                                              ||
## ||    KN150349.1                                                              ||
## ||    KN149796.1                                                              ||
## ||    KN149849.1                                                              ||
## ||    KN150150.1                                                              ||
## ||    KN150203.1                                                              ||
## ||    KN150034.1                                                              ||
## ||    KN149703.1                                                              ||
## ||    KN150562.1                                                              ||
## ||    KN150615.1                                                              ||
## ||    KN150393.1                                                              ||
## ||    KN150446.1                                                              ||
## ||    KN149893.1                                                              ||
## ||    KZ116009.1                                                              ||
## ||    KN149946.1                                                              ||
## ||    KN150300.1                                                              ||
## ||    KN149777.1                                                              ||
## ||    KN150131.1                                                              ||
## ||    KN149800.1                                                              ||
## ||    KN150015.1                                                              ||
## ||    KN150689.1                                                              ||
## ||    KN150490.1                                                              ||
## ||    KN150543.1                                                              ||
## ||    KN150374.1                                                              ||
## ||    KZ116053.1                                                              ||
## ||    KN149990.1                                                              ||
## ||    KN149874.1                                                              ||
## ||    KN150258.1                                                              ||
## ||    KN149927.1                                                              ||
## ||    KN150089.1                                                              ||
## ||    KN149758.1                                                              ||
## ||    KN150112.1                                                              ||
## ||    KZ115960.1                                                              ||
## ||    KN150640.1                                                              ||
## ||    KN150471.1                                                              ||
## ||    KN150355.1                                                              ||
## ||    KN149971.1                                                              ||
## ||    KZ116034.1                                                              ||
## ||    KN150408.1                                                              ||
## ||    KN150186.1                                                              ||
## ||    KN149855.1                                                              ||
## ||    KN149908.1                                                              ||
## ||    KN149686.1                                                              ||
## ||    KN149739.1                                                              ||
## ||    KN150040.1                                                              ||
## ||    KN150598.1                                                              ||
## ||    KN150621.1                                                              ||
## ||    KN150452.1                                                              ||
## ||    KN150505.1                                                              ||
## ||    KN150283.1                                                              ||
## ||    KZ116015.1                                                              ||
## ||    KN150336.1                                                              ||
## ||    KN149783.1                                                              ||
## ||    KN150167.1                                                              ||
## ||    KN149836.1                                                              ||
## ||    KN150021.1                                                              ||
## ||    KN150579.1                                                              ||
## ||    KN150602.1                                                              ||
## ||    KN150380.1                                                              ||
## ||    KN150264.1                                                              ||
## ||    KN149880.1                                                              ||
## ||    KN149933.1                                                              ||
## ||    KN150317.1                                                              ||
## ||    KN150095.1                                                              ||
## ||    KN149764.1                                                              ||
## ||    KN150148.1                                                              ||
## ||    KN149817.1                                                              ||
## ||    KN150002.1                                                              ||
## ||    KZ115996.1                                                              ||
## ||    KN150676.1                                                              ||
## ||    KN150530.1                                                              ||
## ||    KZ116040.1                                                              ||
## ||    KN150361.1                                                              ||
## ||    KN150414.1                                                              ||
## ||    KN149861.1                                                              ||
## ||    KN149914.1                                                              ||
## ||    KN150076.1                                                              ||
## ||    KN149692.1                                                              ||
## ||    KN149745.1                                                              ||
## ||    KN147637.2                                                              ||
## ||    KZ115977.1                                                              ||
## ||    KN150488.1                                                              ||
## ||    KN150511.1                                                              ||
## ||    KN149988.1                                                              ||
## ||    KZ116021.1                                                              ||
## ||    KN150342.1                                                              ||
## ||    KN150173.1                                                              ||
## ||    KN150226.1                                                              ||
## ||    KN150057.1                                                              ||
## ||    KZ115958.1                                                              ||
## ||    KN150585.1                                                              ||
## ||    KN150469.1                                                              ||
## ||    KN149969.1                                                              ||
## ||    KN150270.1                                                              ||
## ||    KZ116002.1                                                              ||
## ||    KN150323.1                                                              ||
## ||    KN149770.1                                                              ||
## ||    KN150154.1                                                              ||
## ||    KN149823.1                                                              ||
## ||    KN150038.1                                                              ||
## ||    KN149707.2                                                              ||
## ||    KN150682.1                                                              ||
## ||    KN150619.1                                                              ||
## ||    KN150397.1                                                              ||
## ||    KN150420.1                                                              ||
## ||    KN149897.1                                                              ||
## ||    KN150251.1                                                              ||
## ||    KN150082.1                                                              ||
## ||    KN149751.1                                                              ||
## ||    KN149804.1                                                              ||
## ||    KN150019.2                                                              ||
## ||    KZ115983.1                                                              ||
## ||    KN150663.1                                                              ||
## ||    KN150494.1                                                              ||
## ||    KN150547.1                                                              ||
## ||    KZ116057.1                                                              ||
## ||    KN150401.1                                                              ||
## ||    KN149878.1                                                              ||
## ||    KN150232.1                                                              ||
## ||    KN149901.1                                                              ||
## ||    KN150063.1                                                              ||
## ||    KN149732.1                                                              ||
## ||    KZ115964.1                                                              ||
## ||    KN150591.1                                                              ||
## ||    KN150644.1                                                              ||
## ||    KN150475.1                                                              ||
## ||    KN150528.1                                                              ||
## ||    KZ116038.1                                                              ||
## ||    KN149859.1                                                              ||
## ||    KN150160.1                                                              ||
## ||    KN150213.1                                                              ||
## ||    KN150044.1                                                              ||
## ||    KN149713.1                                                              ||
## ||    KN147552.2                                                              ||
## ||    KN150572.1                                                              ||
## ||    KN150625.1                                                              ||
## ||    KN150509.1                                                              ||
## ||    KN150287.1                                                              ||
## ||    KN149956.1                                                              ||
## ||    KZ116019.1                                                              ||
## ||    KN150310.1                                                              ||
## ||    KN149787.1                                                              ||
## ||    KN150141.1                                                              ||
## ||    KN149810.1                                                              ||
## ||    KN150025.1                                                              ||
## ||    KN148038.2                                                              ||
## ||    KN150699.1                                                              ||
## ||    KN150553.1                                                              ||
## ||    KN150606.1                                                              ||
## ||    KZ116063.1                                                              ||
## ||    KN150384.1                                                              ||
## ||    KN150437.1                                                              ||
## ||    KN150268.1                                                              ||
## ||    KN149884.1                                                              ||
## ||    KN149937.1                                                              ||
## ||    KN149768.1                                                              ||
## ||    KN150122.1                                                              ||
## ||    KN150006.1                                                              ||
## ||    KZ115970.1                                                              ||
## ||    KN150650.1                                                              ||
## ||    KN150703.1                                                              ||
## ||    KN150481.1                                                              ||
## ||    KN150534.1                                                              ||
## ||    KN150365.1                                                              ||
## ||    KN149981.1                                                              ||
## ||    KZ116044.1                                                              ||
## ||    KN150418.1                                                              ||
## ||    KN150196.1                                                              ||
## ||    KN150249.1                                                              ||
## ||    KN149865.1                                                              ||
## ||    KN149696.2                                                              ||
## ||    KN149749.1                                                              ||
## ||    KN150103.1                                                              ||
## ||    KZ115951.1                                                              ||
## ||    KN150631.1                                                              ||
## ||    KN150462.1                                                              ||
## ||    KN150515.1                                                              ||
## ||    KN150293.1                                                              ||
## ||    KZ116025.1                                                              ||
## ||    KN149793.1                                                              ||
## ||    KN150177.1                                                              ||
## ||    KN149846.1                                                              ||
## ||    KN150200.1                                                              ||
## ||    KN150031.1                                                              ||
## ||    KN149700.1                                                              ||
## ||    KN150589.1                                                              ||
## ||    KN150390.1                                                              ||
## ||    KN150443.1                                                              ||
## ||    KN149890.1                                                              ||
## ||    KN150274.1                                                              ||
## ||    KZ116006.1                                                              ||
## ||    KN150327.1                                                              ||
## ||    KN149943.1                                                              ||
## ||    KN149774.1                                                              ||
## ||    KN150158.1                                                              ||
## ||    KN149827.1                                                              ||
## ||    KN150012.1                                                              ||
## ||    KN150540.1                                                              ||
## ||    KZ116050.1                                                              ||
## ||    KN150371.1                                                              ||
## ||    KN149871.1                                                              ||
## ||    KN150255.2                                                              ||
## ||    KN150308.1                                                              ||
## ||    KN149924.1                                                              ||
## ||    KN150086.1                                                              ||
## ||    KN149755.1                                                              ||
## ||    KZ115987.1                                                              ||
## ||    KN150667.1                                                              ||
## ||    KN150521.1                                                              ||
## ||    KN149998.1                                                              ||
## ||    KZ116031.1                                                              ||
## ||    KN150352.2                                                              ||
## ||    KN150405.1                                                              ||
## ||    KN150183.1                                                              ||
## ||    KN150236.1                                                              ||
## ||    KN149852.1                                                              ||
## ||    KN149905.1                                                              ||
## ||    KN150067.1                                                              ||
## ||    KN149683.1                                                              ||
## ||    KZ115968.1                                                              ||
## ||    KN150595.1                                                              ||
## ||    KN150479.1                                                              ||
## ||    KN150502.1                                                              ||
## ||    KN149979.1                                                              ||
## ||    KN150280.1                                                              ||
## ||    KZ116012.1                                                              ||
## ||    KN150333.1                                                              ||
## ||    KN150164.1                                                              ||
## ||    KN149780.1                                                              ||
## ||    KN150217.1                                                              ||
## ||    KN150048.1                                                              ||
## ||    KN149717.1                                                              ||
## ||    KZ115949.1                                                              ||
## ||    KN150576.1                                                              ||
## ||    KN150629.1                                                              ||
## ||    KN150430.1                                                              ||
## ||    KN150314.1                                                              ||
## ||    KN150092.1                                                              ||
## ||    KN150145.1                                                              ||
## ||    KN149761.1                                                              ||
## ||    KN149814.1                                                              ||
## ||    KN150029.1                                                              ||
## ||    KZ115993.1                                                              ||
## ||    KN150557.1                                                              ||
## ||    KN150388.1                                                              ||
## ||    KZ116067.1                                                              ||
## ||    KN150411.1                                                              ||
## ||    KN149888.1                                                              ||
## ||    KN150242.1                                                              ||
## ||    KN150073.1                                                              ||
## ||    KN149742.1                                                              ||
## ||    KN150126.1                                                              ||
## ||    KZ115974.1                                                              ||
## ||    KN150654.1                                                              ||
## ||    KN150485.1                                                              ||
## ||    KN150538.1                                                              ||
## ||    KZ116048.1                                                              ||
## ||    KN150369.1                                                              ||
## ||    KN150170.1                                                              ||
## ||    KN150054.1                                                              ||
## ||    KN149723.1                                                              ||
## ||    KZ115955.1                                                              ||
## ||    KN150582.1                                                              ||
## ||    KN150635.1                                                              ||
## ||    KN150466.1                                                              ||
## ||    KN150519.1                                                              ||
## ||    KN150297.1                                                              ||
## ||    KZ116029.1                                                              ||
## ||    KN150320.1                                                              ||
## ||    KN149797.1                                                              ||
## ||    KN150204.1                                                              ||
## ||    KN150035.1                                                              ||
## ||    KN150616.1                                                              ||
## ||    KN150394.1                                                              ||
## ||    KN150278.1                                                              ||
## ||    KN150301.1                                                              ||
## ||    KN149778.1                                                              ||
## ||    KN150132.1                                                              ||
## ||    KN149801.1                                                              ||
## ||    KN150016.1                                                              ||
## ||    KZ115980.1                                                              ||
## ||    KN150713.1                                                              ||
## ||    KN150491.1                                                              ||
## ||    KN150544.1                                                              ||
## ||    KN149991.1                                                              ||
## ||    KZ116054.1                                                              ||
## ||    KN150375.1                                                              ||
## ||    KN150428.1                                                              ||
## ||    KN150259.1                                                              ||
## ||    KN149928.1                                                              ||
## ||    KN150060.1                                                              ||
## ||    KN150113.1                                                              ||
## ||    KZ115961.1                                                              ||
## ||    KN150641.1                                                              ||
## ||    KN150472.1                                                              ||
## ||    KN150525.1                                                              ||
## ||    KZ116035.1                                                              ||
## ||    KN150187.1                                                              ||
## ||    KN149856.1                                                              ||
## ||    KN149909.1                                                              ||
## ||    KN150210.1                                                              ||
## ||    KN149687.1                                                              ||
## ||    KN150041.2                                                              ||
## ||    KN149710.1                                                              ||
## ||    KN150599.1                                                              ||
## ||    KN150622.1                                                              ||
## ||    KN150453.1                                                              ||
## ||    KN150506.1                                                              ||
## ||    KN150284.1                                                              ||
## ||    KZ116016.1                                                              ||
## ||    KN150337.1                                                              ||
## ||    KN149953.1                                                              ||
## ||    KN150168.1                                                              ||
## ||    KN149784.1                                                              ||
## ||    KN149837.1                                                              ||
## ||    KN150022.1                                                              ||
## ||    KN150550.1                                                              ||
## ||    KN150603.1                                                              ||
## ||    KZ116060.1                                                              ||
## ||    KN150434.1                                                              ||
## ||    KN149881.1                                                              ||
## ||    KN150265.1                                                              ||
## ||    KN150318.1                                                              ||
## ||    KN149765.1                                                              ||
## ||    KN149818.1                                                              ||
## ||    KZ115997.1                                                              ||
## ||    KN150677.1                                                              ||
## ||    KN150700.2                                                              ||
## ||    KZ116041.1                                                              ||
## ||    KN150362.1                                                              ||
## ||    KN150193.1                                                              ||
## ||    KN149862.1                                                              ||
## ||    KN150246.1                                                              ||
## ||    KN149915.1                                                              ||
## ||    KN149693.1                                                              ||
## ||    KN150077.1                                                              ||
## ||    KN149746.1                                                              ||
## ||    KN150100.1                                                              ||
## ||    KZ115978.1                                                              ||
## ||    KN150489.1                                                              ||
## ||    KN149989.1                                                              ||
## ||    KN150290.1                                                              ||
## ||    KZ116022.1                                                              ||
## ||    KN150343.1                                                              ||
## ||    KN150174.1                                                              ||
## ||    KN149790.1                                                              ||
## ||    KN150227.1                                                              ||
## ||    KN149843.1                                                              ||
## ||    KN150058.1                                                              ||
## ||    KN149727.1                                                              ||
## ||    KZ115959.1                                                              ||
## ||    KN150586.1                                                              ||
## ||    KN150639.1                                                              ||
## ||    KN150440.1                                                              ||
## ||    KN150271.1                                                              ||
## ||    KN149940.1                                                              ||
## ||    KZ116003.1                                                              ||
## ||    KN150324.1                                                              ||
## ||    KN150155.1                                                              ||
## ||    KN149771.1                                                              ||
## ||    KN149824.1                                                              ||
## ||    KN150039.1                                                              ||
## ||    KN149708.1                                                              ||
## ||    KN150683.1                                                              ||
## ||    KN150567.1                                                              ||
## ||    KN150398.1                                                              ||
## ||    KN150421.1                                                              ||
## ||    KN150252.1                                                              ||
## ||    KN149921.1                                                              ||
## ||    KN150083.1                                                              ||
## ||    KN150136.1                                                              ||
## ||    KN149805.1                                                              ||
## ||    KZ115984.1                                                              ||
## ||    KN150664.1                                                              ||
## ||    KN150495.1                                                              ||
## ||    KN150548.1                                                              ||
## ||    KZ116058.1                                                              ||
## ||    KN149995.1                                                              ||
## ||    KN150379.1                                                              ||
## ||    KN149879.1                                                              ||
## ||    KN150180.1                                                              ||
## ||    KN150233.1                                                              ||
## ||    KN150064.1                                                              ||
## ||    KN149680.1                                                              ||
## ||    KN149733.1                                                              ||
## ||    KN150117.1                                                              ||
## ||    KZ115965.1                                                              ||
## ||    KN150592.1                                                              ||
## ||    KN150645.1                                                              ||
## ||    KN150476.1                                                              ||
## ||    KN150529.1                                                              ||
## ||    KN149976.1                                                              ||
## ||    KZ116039.1                                                              ||
## ||    KN150330.1                                                              ||
## ||    KN150161.1                                                              ||
## ||    KN150214.1                                                              ||
## ||    KN150045.2                                                              ||
## ||    KN149714.1                                                              ||
## ||    KN150573.1                                                              ||
## ||    KN150626.1                                                              ||
## ||    KN149957.1                                                              ||
## ||    KN150311.1                                                              ||
## ||    KN149788.1                                                              ||
## ||    KN149811.1                                                              ||
## ||    KZ115990.1                                                              ||
## ||    KN150670.1                                                              ||
## ||    KN150554.1                                                              ||
## ||    KN150607.1                                                              ||
## ||    KN150385.1                                                              ||
## ||    KZ116064.1                                                              ||
## ||    KN149885.1                                                              ||
## ||    KN150269.1                                                              ||
## ||    KN149938.1                                                              ||
## ||    KN149769.1                                                              ||
## ||    KN150070.1                                                              ||
## ||    KN150123.1                                                              ||
## ||    KN150007.1                                                              ||
## ||    KZ115971.1                                                              ||
## ||    KN150651.1                                                              ||
## ||    KN150535.1                                                              ||
## ||    KZ116045.1                                                              ||
## ||    KN150366.1                                                              ||
## ||    KN149982.1                                                              ||
## ||    KN150419.1                                                              ||
## ||    KN150197.1                                                              ||
## ||    KN149866.1                                                              ||
## ||    KN149919.1                                                              ||
## ||    KN150220.1                                                              ||
## ||    KN149697.1                                                              ||
## ||    KN150051.1                                                              ||
## ||    KN149720.1                                                              ||
## ||    KN150104.1                                                              ||
## ||    KZ115952.1                                                              ||
## ||    KN150632.1                                                              ||
## ||    KN150516.1                                                              ||
## ||    KZ116026.1                                                              ||
## ||    KN150347.1                                                              ||
## ||    KN149794.1                                                              ||
## ||    KN149847.1                                                              ||
## ||    KN150201.1                                                              ||
## ||    KN150032.1                                                              ||
## ||    KN149701.1                                                              ||
## ||    KN150560.1                                                              ||
## ||    KN150613.1                                                              ||
## ||    KN150444.1                                                              ||
## ||    KN150275.1                                                              ||
## ||    KZ116007.1                                                              ||
## ||    KN150328.1                                                              ||
## ||    KN149944.1                                                              ||
## ||    KN150159.1                                                              ||
## ||    KN149775.2                                                              ||
## ||    KN149828.1                                                              ||
## ||    KN150013.1                                                              ||
## ||    KN150687.1                                                              ||
## ||    KN148869.2                                                              ||
## ||    KN150710.1                                                              ||
## ||    KN150541.1                                                              ||
## ||    KZ116051.1                                                              ||
## ||    KN150372.1                                                              ||
## ||    KN150425.1                                                              ||
## ||    KN149872.1                                                              ||
## ||    KN150309.1                                                              ||
## ||    KN149925.1                                                              ||
## ||    KN149756.1                                                              ||
## ||    KN149809.1                                                              ||
## ||    KN150110.1                                                              ||
## ||    KZ115988.1                                                              ||
## ||    KN150668.1                                                              ||
## ||    KN150499.1                                                              ||
## ||    KN150522.1                                                              ||
## ||    KN149999.1                                                              ||
## ||    KZ116032.1                                                              ||
## ||    KN150353.1                                                              ||
## ||    KN150406.1                                                              ||
## ||    KN150184.1                                                              ||
## ||    KN150237.1                                                              ||
## ||    KN149853.1                                                              ||
## ||    KN149906.1                                                              ||
## ||    KN150068.1                                                              ||
## ||    KN149684.1                                                              ||
## ||    KN149737.1                                                              ||
## ||    KZ115969.1                                                              ||
## ||    KN150596.1                                                              ||
## ||    KN150649.1                                                              ||
## ||    KN150450.1                                                              ||
## ||    KN150503.1                                                              ||
## ||    KN149950.1                                                              ||
## ||    KZ116013.1                                                              ||
## ||    KN150334.1                                                              ||
## ||    KN150165.1                                                              ||
## ||    KN150218.1                                                              ||
## ||    KN149834.1                                                              ||
## ||    KN150049.1                                                              ||
## ||    KN149718.1                                                              ||
## ||    KN150577.1                                                              ||
## ||    KN150431.1                                                              ||
## ||    KN149931.1                                                              ||
## ||    KN150315.1                                                              ||
## ||    KN150093.1                                                              ||
## ||    KN150146.1                                                              ||
## ||    KN149762.1                                                              ||
## ||    KN149815.1                                                              ||
## ||    KN150000.1                                                              ||
## ||    KZ115994.1                                                              ||
## ||    KN150558.1                                                              ||
## ||    KN150389.1                                                              ||
## ||    KN150412.1                                                              ||
## ||    KN149889.1                                                              ||
## ||    KN150190.1                                                              ||
## ||    KN150243.2                                                              ||
## ||    KN149912.1                                                              ||
## ||    KN150074.1                                                              ||
## ||    KN149690.1                                                              ||
## ||    KN149743.1                                                              ||
## ||    KN150127.1                                                              ||
## ||    KZ115975.1                                                              ||
## ||    KN150655.1                                                              ||
## ||    KN150708.1                                                              ||
## ||    KN150486.1                                                              ||
## ||    KZ116049.1                                                              ||
## ||    KN150340.1                                                              ||
## ||    KN150171.1                                                              ||
## ||    KN150224.1                                                              ||
## ||    KN149840.1                                                              ||
## ||    KN150055.1                                                              ||
## ||    KN149724.1                                                              ||
## ||    KZ115956.1                                                              ||
## ||    KN150583.1                                                              ||
## ||    KN150636.1                                                              ||
## ||    KN150467.1                                                              ||
## ||    KN149967.1                                                              ||
## ||    KZ116000.1                                                              ||
## ||    KN149798.1                                                              ||
## ||    KN150152.1                                                              ||
## ||    KN150205.1                                                              ||
## ||    KN149705.1                                                              ||
## ||    KN150680.1                                                              ||
## ||    KN150564.1                                                              ||
## ||    KN150617.1                                                              ||
## ||    KN150395.1                                                              ||
## ||    KN150279.1                                                              ||
## ||    KN149895.1                                                              ||
## ||    KN149948.1                                                              ||
## ||    KN150302.1                                                              ||
## ||    KN149779.1                                                              ||
## ||    KN150080.1                                                              ||
## ||    KN150133.2                                                              ||
## ||    KN150017.1                                                              ||
## ||    KZ115981.1                                                              ||
## ||    KN150661.1                                                              ||
## ||    KN150492.2                                                              ||
## ||    KN150545.1                                                              ||
## ||    KZ116055.1                                                              ||
## ||    KN149992.1                                                              ||
## ||    KN150429.1                                                              ||
## ||    KN149876.1                                                              ||
## ||    KN150230.1                                                              ||
## ||    KN150061.1                                                              ||
## ||    KN149730.1                                                              ||
## ||    KN150114.1                                                              ||
## ||    KZ115962.1                                                              ||
## ||    KN150642.1                                                              ||
## ||    KN150473.1                                                              ||
## ||    KN150526.1                                                              ||
## ||    KZ116036.1                                                              ||
## ||    KN150357.1                                                              ||
## ||    KN149973.1                                                              ||
## ||    KN149857.1                                                              ||
## ||    KN150211.1                                                              ||
## ||    KN149688.2                                                              ||
## ||    KN150042.1                                                              ||
## ||    KN149711.1                                                              ||
## ||    KN150570.1                                                              ||
## ||    KN150623.1                                                              ||
## ||    KN150454.1                                                              ||
## ||    KN150285.1                                                              ||
## ||    KN150338.1                                                              ||
## ||    KZ116017.1                                                              ||
## ||    KN149785.1                                                              ||
## ||    KN149838.1                                                              ||
## ||    KN150023.1                                                              ||
## ||    KN150697.1                                                              ||
## ||    KN150551.1                                                              ||
## ||    KZ116061.1                                                              ||
## ||    KN150382.1                                                              ||
## ||    KN149882.1                                                              ||
## ||    KN149935.1                                                              ||
## ||    KN150319.1                                                              ||
## ||    KN150097.1                                                              ||
## ||    KN149766.1                                                              ||
## ||    KN149819.1                                                              ||
## ||    KN150120.1                                                              ||
## ||    KN150004.2                                                              ||
## ||    KZ115998.1                                                              ||
## ||    KN150678.1                                                              ||
## ||    KN150701.1                                                              ||
## ||    KN150532.1                                                              ||
## ||    KZ116042.1                                                              ||
## ||    KN150363.1                                                              ||
## ||    KN150416.1                                                              ||
## ||    KN150194.1                                                              ||
## ||    KN149863.1                                                              ||
## ||    KN149916.1                                                              ||
## ||    KN150078.1                                                              ||
## ||    KN149694.1                                                              ||
## ||    KN149747.1                                                              ||
## ||    KZ115979.1                                                              ||
## ||    KN150659.1                                                              ||
## ||    KN150513.1                                                              ||
## ||    KN150291.1                                                              ||
## ||    KZ116023.1                                                              ||
## ||    KN149960.1                                                              ||
## ||    KN150344.1                                                              ||
## ||    KN149791.1                                                              ||
## ||    KN150175.1                                                              ||
## ||    KN149844.1                                                              ||
## ||    KN150228.1                                                              ||
## ||    KN150059.1                                                              ||
## ||    KN149728.1                                                              ||
## ||    KN150587.1                                                              ||
## ||    KN150610.1                                                              ||
## ||    KN150441.1                                                              ||
## ||    KN150272.1                                                              ||
## ||    KN149941.1                                                              ||
## ||    KZ116004.1                                                              ||
## ||    KN150325.1                                                              ||
## ||    KN149772.1                                                              ||
## ||    KN150156.1                                                              ||
## ||    KN149825.1                                                              ||
## ||    KN149709.1                                                              ||
## ||    KN150684.1                                                              ||
## ||    KN150568.1                                                              ||
## ||    KN150399.1                                                              ||
## ||    KN150422.1                                                              ||
## ||    KN149899.1                                                              ||
## ||    KN150253.1                                                              ||
## ||    KN149922.1                                                              ||
## ||    KN150306.1                                                              ||
## ||    KN150084.1                                                              ||
## ||    KN149753.1                                                              ||
## ||    KN150137.1                                                              ||
## ||    KN149806.1                                                              ||
## ||    KZ115985.1                                                              ||
## ||                                                                            ||
## || Global environment is initialised.                                         ||
## || Load the 1-th index block...                                               ||
## || The index block has been loaded.                                           ||
## || Start read mapping in chunk.                                               ||
## ||    2% completed, 0.1 mins elapsed, rate=101.7k reads per second            ||
## ||    9% completed, 0.1 mins elapsed, rate=106.2k reads per second            ||
## ||   15% completed, 0.1 mins elapsed, rate=107.4k reads per second            ||
## ||   22% completed, 0.1 mins elapsed, rate=107.2k reads per second            ||
## ||   29% completed, 0.1 mins elapsed, rate=107.2k reads per second            ||
## ||   36% completed, 0.2 mins elapsed, rate=107.7k reads per second            ||
## ||   42% completed, 0.2 mins elapsed, rate=107.6k reads per second            ||
## ||   49% completed, 0.2 mins elapsed, rate=107.9k reads per second            ||
## ||   56% completed, 0.2 mins elapsed, rate=108.1k reads per second            ||
## ||   63% completed, 0.2 mins elapsed, rate=108.3k reads per second            ||
## ||   69% completed, 0.3 mins elapsed, rate=59.8k reads per second             ||
## ||   73% completed, 0.3 mins elapsed, rate=60.0k reads per second             ||
## ||   76% completed, 0.3 mins elapsed, rate=60.2k reads per second             ||
## ||   79% completed, 0.4 mins elapsed, rate=60.3k reads per second             ||
## ||   83% completed, 0.4 mins elapsed, rate=60.4k reads per second             ||
## ||   86% completed, 0.4 mins elapsed, rate=60.5k reads per second             ||
## ||   89% completed, 0.4 mins elapsed, rate=60.7k reads per second             ||
## ||   93% completed, 0.4 mins elapsed, rate=60.8k reads per second             ||
## ||   96% completed, 0.4 mins elapsed, rate=60.9k reads per second             ||
## ||                                                                            ||
## ||                           Completed successfully.                          ||
## ||                                                                            ||
## \\====================================    ====================================//
## 
## //================================   Summary =================================\\
## ||                                                                            ||
## ||                 Total reads : 1,624,879                                    ||
## ||                      Mapped : 1,252,755 (77.1%)                            ||
## ||             Uniquely mapped : 878,549                                      ||
## ||               Multi-mapping : 374,206                                      ||
## ||                                                                            ||
## ||                    Unmapped : 372,124                                      ||
## ||                                                                            ||
## ||                   Junctions : 590                                          ||
## ||                      Indels : 0                                            ||
## ||                                                                            ||
## ||                Running time : 0.4 minutes                                  ||
## ||                                                                            ||
## \\============================================================================//
```

```
sortBam(file = 'Heart/bam/fish_heart_captured_sequence.BAM',
        destination = 'Heart/bam/fish_heart_captured_sequence.sorted')
```

```
## [1] "Heart/bam/fish_heart_captured_sequence.sorted.bam"
```

```
indexBam('Heart/bam/fish_heart_captured_sequence.sorted.bam')
```

```
##       Heart/bam/fish_heart_captured_sequence.sorted.bam 
## "Heart/bam/fish_heart_captured_sequence.sorted.bam.bai"
```

```
counts = featureCounts(
  files = 'Heart/bam/fish_heart_captured_sequence.sorted.bam',
#  minMQS = 10,
  annot.ext = gtf_file,
  isGTFAnnotationFile = TRUE,
  nthreads = 8)
```

```
## 
##         ==========     _____ _    _ ____  _____  ______          _____  
##         =====         / ____| |  | |  _ \|  __ \|  ____|   /\   |  __ \ 
##           =====      | (___ | |  | | |_) | |__) | |__     /  \  | |  | |
##             ====      \___ \| |  | |  _ <|  _  /|  __|   / /\ \ | |  | |
##               ====    ____) | |__| | |_) | | \ \| |____ / ____ \| |__| |
##         ==========   |_____/ \____/|____/|_|  \_\______/_/    \_\_____/
##        Rsubread 2.0.1
## 
## //========================== featureCounts setting ===========================\\
## ||                                                                            ||
## ||             Input files : 1 BAM file                                       ||
## ||                           o fish_heart_captured_sequence.sorted.bam        ||
## ||                                                                            ||
## ||              Annotation : Danio_rerio.GRCz11.93.chr.gtf (GTF)              ||
## ||      Dir for temp files : .                                                ||
## ||                 Threads : 8                                                ||
## ||                   Level : meta-feature level                               ||
## ||              Paired-end : no                                               ||
## ||      Multimapping reads : counted                                          ||
## || Multi-overlapping reads : not counted                                      ||
## ||   Min overlapping bases : 1                                                ||
## ||                                                                            ||
## \\============================================================================//
## 
## //================================= Running ==================================\\
## ||                                                                            ||
## || Load annotation file Danio_rerio.GRCz11.93.chr.gtf ...                     ||
## ||    Features : 481079                                                       ||
## ||    Meta-features : 32057                                                   ||
## ||    Chromosomes/contigs : 26                                                ||
## ||                                                                            ||
## || Process BAM file fish_heart_captured_sequence.sorted.bam...                ||
## ||    Single-end reads are included.                                          ||
## ||    Total alignments : 1624879                                              ||
## ||    Successfully assigned alignments : 593938 (36.6%)                       ||
## ||    Running time : 0.01 minutes                                             ||
## ||                                                                            ||
## || Write the final count table.                                               ||
## || Write the read assignment summary.                                         ||
## ||                                                                            ||
## \\============================================================================//
```

```
genecounts = counts$counts
sum(genecounts[, 1] > 0)
```

```
## [1] 13329
```

#### 48 hpf Heart RNA-seq Data

```
for (heartfile in list.files("/home/jth54/Timecourse/9499R/Fastq/", 
                             "^9499X(4|12|20)_.*.txt.gz$", full.names = TRUE)){
  basename <- gsub(".*//(.*?)_.*.txt.gz", "\\1", heartfile)
  bamfile <- paste0("Heart/bams/", basename, ".bam")
  
  subjunc(index = reference_genome,
          readfile1 = heartfile,
          output_file = bamfile,
          TH1 = 1,
          nthreads = 8,
          useAnnotation = TRUE,
          annot.ext = gtf_file,
          isGTF = TRUE)
  
  sortBam(bamfile, paste0("Heart/bams/", basename, ".sorted"))
  
  indexBam(paste0("Heart/bams/", basename, ".sorted.bam"))}
```

```
## 
##         ==========     _____ _    _ ____  _____  ______          _____  
##         =====         / ____| |  | |  _ \|  __ \|  ____|   /\   |  __ \ 
##           =====      | (___ | |  | | |_) | |__) | |__     /  \  | |  | |
##             ====      \___ \| |  | |  _ <|  _  /|  __|   / /\ \ | |  | |
##               ====    ____) | |__| | |_) | | \ \| |____ / ____ \| |__| |
##         ==========   |_____/ \____/|____/|_|  \_\______/_/    \_\_____/
##        Rsubread 2.0.1
## 
## //================================= setting ==================================\\
## ||                                                                            ||
## || Function      : Read alignment + Junction detection (RNA-Seq)              ||
## || Input file    : 9499X12_130405_SN141_0668_AD2154ACXX_8.txt.gz              ||
## || Output file   : 9499X12.bam (BAM)                                          ||
## || Index name    : GRCz11                                                     ||
## ||                                                                            ||
## ||                    ------------------------------------                    ||
## ||                                                                            ||
## ||                               Threads : 8                                  ||
## ||                          Phred offset : 33                                 ||
## ||                             Min votes : 1 / 14                             ||
## ||                        Max mismatches : 3                                  ||
## ||                      Max indel length : 5                                  ||
## ||            Report multi-mapping reads : yes                                ||
## || Max alignments per multi-mapping read : 1                                  ||
## ||                           Annotations : Danio_rerio.GRCz11.93.chr.gtf  ... ||
## ||                                                                            ||
## \\============================================================================//
## 
## //================ Running (24-Jun-2020 10:58:23, pid=12013) =================\\
## ||                                                                            ||
## || Check the input reads.                                                     ||
## || The input file contains base space reads.                                  ||
## || Initialise the memory objects.                                             ||
## || Estimate the mean read length.                                             ||
## || The range of Phred scores observed in the data is [2,41]                   ||
## || Create the output BAM file.                                                ||
## || Check the index.                                                           ||
## || Init the voting space.                                                     ||
## || Load the annotation file.                                                  ||
## || 481079 annotation records were loaded.                                     ||
## ||                                                                            ||
## || Chromosomes/contigs in index but not in annotation :                       ||
## ||    KN150665.1                                                              ||
## ||    KN150549.1                                                              ||
## ||    KN149996.1                                                              ||
## ||    KZ116059.1                                                              ||
## ||    KN150350.1                                                              ||
## ||    KN150403.1                                                              ||
## ||    KN150181.1                                                              ||
## ||    KN150234.1                                                              ||
## ||    KN149850.1                                                              ||
## ||    KN149903.1                                                              ||
## ||    KN150065.1                                                              ||
## ||    KN150118.1                                                              ||
## ||    KZ115966.1                                                              ||
## ||    KN150593.1                                                              ||
## ||    KN150646.1                                                              ||
## ||    KN148828.2                                                              ||
## ||    KN150477.1                                                              ||
## ||    KN150500.1                                                              ||
## ||    KN149977.2                                                              ||
## ||    KZ116010.1                                                              ||
## ||    KN150331.1                                                              ||
## ||    KN150162.1                                                              ||
## ||    KN150215.1                                                              ||
## ||    KN149715.1                                                              ||
## ||    KN150690.1                                                              ||
## ||    KN150574.1                                                              ||
## ||    KN150458.1                                                              ||
## ||    KN150289.1                                                              ||
## ||    KN149958.1                                                              ||
## ||    KN150312.1                                                              ||
## ||    KN149789.1                                                              ||
## ||    KN150090.1                                                              ||
## ||    KN149812.1                                                              ||
## ||    KN147651.2                                                              ||
## ||    KZ115991.1                                                              ||
## ||    KN150671.1                                                              ||
## ||    KN150555.1                                                              ||
## ||    KN150608.1                                                              ||
## ||    KN150386.1                                                              ||
## ||    KZ116065.1                                                              ||
## ||    KN150439.1                                                              ||
## ||    KN149939.1                                                              ||
## ||    KN150240.1                                                              ||
## ||    KN150071.1                                                              ||
## ||    KN149740.1                                                              ||
## ||    KN150008.1                                                              ||
## ||    KN147632.2                                                              ||
## ||    KZ115972.1                                                              ||
## ||    KN150652.1                                                              ||
## ||    KN150483.1                                                              ||
## ||    KN150536.1                                                              ||
## ||    KZ116046.1                                                              ||
## ||    KN149983.1                                                              ||
## ||    KN149867.1                                                              ||
## ||    KN150221.1                                                              ||
## ||    KN149698.1                                                              ||
## ||    KN150052.1                                                              ||
## ||    KN149721.1                                                              ||
## ||    KN150105.1                                                              ||
## ||    KZ115953.1                                                              ||
## ||    KN150580.1                                                              ||
## ||    KN150633.1                                                              ||
## ||    KN150464.1                                                              ||
## ||    KN150517.1                                                              ||
## ||    KZ116027.1                                                              ||
## ||    KN149795.1                                                              ||
## ||    KN150179.1                                                              ||
## ||    KN149848.1                                                              ||
## ||    KN150202.1                                                              ||
## ||    KN150033.1                                                              ||
## ||    KN149702.1                                                              ||
## ||    KN150561.1                                                              ||
## ||    KN150614.1                                                              ||
## ||    KN150392.1                                                              ||
## ||    KN150445.1                                                              ||
## ||    KN150276.1                                                              ||
## ||    KN149892.1                                                              ||
## ||    KN150329.1                                                              ||
## ||    KZ116008.1                                                              ||
## ||    KN149829.1                                                              ||
## ||    KN150130.1                                                              ||
## ||    KN150014.1                                                              ||
## ||    KN150711.1                                                              ||
## ||    KN150542.1                                                              ||
## ||    KZ116052.1                                                              ||
## ||    KN150426.1                                                              ||
## ||    KN149873.1                                                              ||
## ||    KN150257.1                                                              ||
## ||    KN149926.1                                                              ||
## ||    KN149757.1                                                              ||
## ||    KN150111.1                                                              ||
## ||    KZ115989.1                                                              ||
## ||    KN150669.1                                                              ||
## ||    KN150470.1                                                              ||
## ||    KN150523.1                                                              ||
## ||    KN150354.1                                                              ||
## ||    KN149970.1                                                              ||
## ||    KZ116033.1                                                              ||
## ||    KN150407.1                                                              ||
## ||    KN150185.1                                                              ||
## ||    KN149854.1                                                              ||
## ||    KN150238.1                                                              ||
## ||    KN149907.1                                                              ||
## ||    KN149685.1                                                              ||
## ||    KN149738.1                                                              ||
## ||    KN150620.1                                                              ||
## ||    KN150451.1                                                              ||
## ||    KN150504.1                                                              ||
## ||    KN150282.1                                                              ||
## ||    KZ116014.1                                                              ||
## ||    KN149951.1                                                              ||
## ||    KN150335.1                                                              ||
## ||    KN149782.1                                                              ||
## ||    KN150166.1                                                              ||
## ||    KN150219.1                                                              ||
## ||    KN149835.1                                                              ||
## ||    KN149719.1                                                              ||
## ||    KN150020.1                                                              ||
## ||    KN150578.1                                                              ||
## ||    KN150601.1                                                              ||
## ||    KN150432.1                                                              ||
## ||    KN150263.1                                                              ||
## ||    KN149932.1                                                              ||
## ||    KN150094.1                                                              ||
## ||    KN150147.1                                                              ||
## ||    KN149763.1                                                              ||
## ||    KN149816.1                                                              ||
## ||    KZ115995.1                                                              ||
## ||    KN150675.1                                                              ||
## ||    KN150559.1                                                              ||
## ||    KN150360.1                                                              ||
## ||    KN150413.1                                                              ||
## ||    KN150191.1                                                              ||
## ||    KN149860.1                                                              ||
## ||    KN150244.1                                                              ||
## ||    KN149913.1                                                              ||
## ||    KN149691.1                                                              ||
## ||    KN150128.1                                                              ||
## ||    KN147636.1                                                              ||
## ||    KZ115976.1                                                              ||
## ||    KN150656.2                                                              ||
## ||    KN150709.1                                                              ||
## ||    KN150487.1                                                              ||
## ||    KN150510.1                                                              ||
## ||    KZ116020.1                                                              ||
## ||    KN150341.1                                                              ||
## ||    KN150172.1                                                              ||
## ||    KN149841.1                                                              ||
## ||    KN150225.1                                                              ||
## ||    KN150056.1                                                              ||
## ||    KN149725.1                                                              ||
## ||    KZ115957.1                                                              ||
## ||    KN150468.1                                                              ||
## ||    KN150299.1                                                              ||
## ||    KN149968.1                                                              ||
## ||    KZ116001.1                                                              ||
## ||    KN150322.1                                                              ||
## ||    KN150206.1                                                              ||
## ||    KN149822.1                                                              ||
## ||    KN150037.1                                                              ||
## ||    KN149706.1                                                              ||
## ||    KN150565.1                                                              ||
## ||    KN150396.1                                                              ||
## ||    KN150449.1                                                              ||
## ||    KN149896.1                                                              ||
## ||    KN149949.1                                                              ||
## ||    KN150250.1                                                              ||
## ||    KN150303.1                                                              ||
## ||    KN150081.1                                                              ||
## ||    KN149750.1                                                              ||
## ||    KN150134.1                                                              ||
## ||    KN149803.1                                                              ||
## ||    KN150018.1                                                              ||
## ||    KN147642.2                                                              ||
## ||    KZ115982.1                                                              ||
## ||    KN150662.1                                                              ||
## ||    KN150493.1                                                              ||
## ||    KN150546.1                                                              ||
## ||    KZ116056.1                                                              ||
## ||    KN149993.1                                                              ||
## ||    KN150400.1                                                              ||
## ||    KN150231.1                                                              ||
## ||    KN149900.1                                                              ||
## ||    KN150062.1                                                              ||
## ||    KN150115.1                                                              ||
## ||    KZ115963.1                                                              ||
## ||    KN150643.1                                                              ||
## ||    KN150527.1                                                              ||
## ||    KN150358.1                                                              ||
## ||    KZ116037.1                                                              ||
## ||    KN149974.1                                                              ||
## ||    KN150189.1                                                              ||
## ||    KN150212.1                                                              ||
## ||    KN149689.2                                                              ||
## ||    KN150043.1                                                              ||
## ||    KN150571.1                                                              ||
## ||    KN150455.1                                                              ||
## ||    KN150508.1                                                              ||
## ||    KN150286.1                                                              ||
## ||    KZ116018.1                                                              ||
## ||    KN149955.1                                                              ||
## ||    KN149786.1                                                              ||
## ||    KN149839.1                                                              ||
## ||    KN150140.2                                                              ||
## ||    KN150024.1                                                              ||
## ||    KN150698.1                                                              ||
## ||    KN150552.1                                                              ||
## ||    KN150605.1                                                              ||
## ||    KZ116062.1                                                              ||
## ||    KN150383.1                                                              ||
## ||    KN150436.1                                                              ||
## ||    KN149883.1                                                              ||
## ||    KN150267.1                                                              ||
## ||    KN149936.1                                                              ||
## ||    KN150098.1                                                              ||
## ||    KN149767.1                                                              ||
## ||    KN150121.1                                                              ||
## ||    KN150005.1                                                              ||
## ||    KZ115999.1                                                              ||
## ||    KN150679.1                                                              ||
## ||    KN149980.1                                                              ||
## ||    KN150364.1                                                              ||
## ||    KZ116043.1                                                              ||
## ||    KN150195.1                                                              ||
## ||    KN149864.1                                                              ||
## ||    KN150248.1                                                              ||
## ||    KN149917.1                                                              ||
## ||    KN150079.1                                                              ||
## ||    KN149695.1                                                              ||
## ||    KN149748.1                                                              ||
## ||    KN150102.1                                                              ||
## ||    KZ115950.1                                                              ||
## ||    KN150630.1                                                              ||
## ||    KN150514.1                                                              ||
## ||    KN150292.1                                                              ||
## ||    KN150345.1                                                              ||
## ||    KN149961.1                                                              ||
## ||    KZ116024.1                                                              ||
## ||    KN149792.1                                                              ||
## ||    KN150176.1                                                              ||
## ||    KN150229.1                                                              ||
## ||    KN149729.1                                                              ||
## ||    KN150030.1                                                              ||
## ||    KN150588.1                                                              ||
## ||    KZ116005.1                                                              ||
## ||    KN150326.1                                                              ||
## ||    KN150157.1                                                              ||
## ||    KN149826.1                                                              ||
## ||    KN150011.1                                                              ||
## ||    KN150685.1                                                              ||
## ||    KN150254.1                                                              ||
## ||    KN150307.1                                                              ||
## ||    KN149923.1                                                              ||
## ||    KN150085.1                                                              ||
## ||    KN149754.1                                                              ||
## ||    KN149807.1                                                              ||
## ||    KZ115986.1                                                              ||
## ||    KN150666.1                                                              ||
## ||    KN150520.1                                                              ||
## ||    KN149997.1                                                              ||
## ||    KZ116030.1                                                              ||
## ||    KN150351.1                                                              ||
## ||    KN150404.1                                                              ||
## ||    KN150182.1                                                              ||
## ||    KN149851.1                                                              ||
## ||    KN149904.1                                                              ||
## ||    KN150066.1                                                              ||
## ||    KN149682.1                                                              ||
## ||    KN150119.1                                                              ||
## ||    KN149735.1                                                              ||
## ||    KZ115967.1                                                              ||
## ||    KN150594.1                                                              ||
## ||    KN150647.1                                                              ||
## ||    KN150478.1                                                              ||
## ||    KN150501.1                                                              ||
## ||    KN149978.1                                                              ||
## ||    KN150332.1                                                              ||
## ||    KZ116011.1                                                              ||
## ||    KN150216.1                                                              ||
## ||    KN149832.1                                                              ||
## ||    KN150047.1                                                              ||
## ||    KN149716.1                                                              ||
## ||    KN150691.1                                                              ||
## ||    KZ115948.1                                                              ||
## ||    KN150575.1                                                              ||
## ||    KN150628.1                                                              ||
## ||    KN150459.1                                                              ||
## ||    KN149959.1                                                              ||
## ||    KN150260.1                                                              ||
## ||    KN150313.1                                                              ||
## ||    KN150144.1                                                              ||
## ||    KN149760.1                                                              ||
## ||    KN149813.1                                                              ||
## ||    KN150028.1                                                              ||
## ||    KN147652.2                                                              ||
## ||    KZ115992.1                                                              ||
## ||    KN150556.1                                                              ||
## ||    KN150609.1                                                              ||
## ||    KN150387.1                                                              ||
## ||    KZ116066.1                                                              ||
## ||    KN149887.1                                                              ||
## ||    KN150241.1                                                              ||
## ||    KN150072.1                                                              ||
## ||    KN149741.1                                                              ||
## ||    KN150125.1                                                              ||
## ||    KN150009.1                                                              ||
## ||    KZ115973.1                                                              ||
## ||    KN150653.1                                                              ||
## ||    KN150484.1                                                              ||
## ||    KN150537.1                                                              ||
## ||    KZ116047.1                                                              ||
## ||    KN150368.1                                                              ||
## ||    KN149984.1                                                              ||
## ||    KN150199.1                                                              ||
## ||    KN150222.1                                                              ||
## ||    KN149722.1                                                              ||
## ||    KN150106.1                                                              ||
## ||    KZ115954.1                                                              ||
## ||    KN150581.1                                                              ||
## ||    KN150634.1                                                              ||
## ||    KN150465.1                                                              ||
## ||    KN150518.1                                                              ||
## ||    KN150296.1                                                              ||
## ||    KZ116028.1                                                              ||
## ||    KN149965.1                                                              ||
## ||    KN150349.1                                                              ||
## ||    KN149796.1                                                              ||
## ||    KN149849.1                                                              ||
## ||    KN150150.1                                                              ||
## ||    KN150203.1                                                              ||
## ||    KN150034.1                                                              ||
## ||    KN149703.1                                                              ||
## ||    KN150562.1                                                              ||
## ||    KN150615.1                                                              ||
## ||    KN150393.1                                                              ||
## ||    KN150446.1                                                              ||
## ||    KN149893.1                                                              ||
## ||    KZ116009.1                                                              ||
## ||    KN149946.1                                                              ||
## ||    KN150300.1                                                              ||
## ||    KN149777.1                                                              ||
## ||    KN150131.1                                                              ||
## ||    KN149800.1                                                              ||
## ||    KN150015.1                                                              ||
## ||    KN150689.1                                                              ||
## ||    KN150490.1                                                              ||
## ||    KN150543.1                                                              ||
## ||    KN150374.1                                                              ||
## ||    KZ116053.1                                                              ||
## ||    KN149990.1                                                              ||
## ||    KN149874.1                                                              ||
## ||    KN150258.1                                                              ||
## ||    KN149927.1                                                              ||
## ||    KN150089.1                                                              ||
## ||    KN149758.1                                                              ||
## ||    KN150112.1                                                              ||
## ||    KZ115960.1                                                              ||
## ||    KN150640.1                                                              ||
## ||    KN150471.1                                                              ||
## ||    KN150355.1                                                              ||
## ||    KN149971.1                                                              ||
## ||    KZ116034.1                                                              ||
## ||    KN150408.1                                                              ||
## ||    KN150186.1                                                              ||
## ||    KN149855.1                                                              ||
## ||    KN149908.1                                                              ||
## ||    KN149686.1                                                              ||
## ||    KN149739.1                                                              ||
## ||    KN150040.1                                                              ||
## ||    KN150598.1                                                              ||
## ||    KN150621.1                                                              ||
## ||    KN150452.1                                                              ||
## ||    KN150505.1                                                              ||
## ||    KN150283.1                                                              ||
## ||    KZ116015.1                                                              ||
## ||    KN150336.1                                                              ||
## ||    KN149783.1                                                              ||
## ||    KN150167.1                                                              ||
## ||    KN149836.1                                                              ||
## ||    KN150021.1                                                              ||
## ||    KN150579.1                                                              ||
## ||    KN150602.1                                                              ||
## ||    KN150380.1                                                              ||
## ||    KN150264.1                                                              ||
## ||    KN149880.1                                                              ||
## ||    KN149933.1                                                              ||
## ||    KN150317.1                                                              ||
## ||    KN150095.1                                                              ||
## ||    KN149764.1                                                              ||
## ||    KN150148.1                                                              ||
## ||    KN149817.1                                                              ||
## ||    KN150002.1                                                              ||
## ||    KZ115996.1                                                              ||
## ||    KN150676.1                                                              ||
## ||    KN150530.1                                                              ||
## ||    KZ116040.1                                                              ||
## ||    KN150361.1                                                              ||
## ||    KN150414.1                                                              ||
## ||    KN149861.1                                                              ||
## ||    KN149914.1                                                              ||
## ||    KN150076.1                                                              ||
## ||    KN149692.1                                                              ||
## ||    KN149745.1                                                              ||
## ||    KN147637.2                                                              ||
## ||    KZ115977.1                                                              ||
## ||    KN150488.1                                                              ||
## ||    KN150511.1                                                              ||
## ||    KN149988.1                                                              ||
## ||    KZ116021.1                                                              ||
## ||    KN150342.1                                                              ||
## ||    KN150173.1                                                              ||
## ||    KN150226.1                                                              ||
## ||    KN150057.1                                                              ||
## ||    KZ115958.1                                                              ||
## ||    KN150585.1                                                              ||
## ||    KN150469.1                                                              ||
## ||    KN149969.1                                                              ||
## ||    KN150270.1                                                              ||
## ||    KZ116002.1                                                              ||
## ||    KN150323.1                                                              ||
## ||    KN149770.1                                                              ||
## ||    KN150154.1                                                              ||
## ||    KN149823.1                                                              ||
## ||    KN150038.1                                                              ||
## ||    KN149707.2                                                              ||
## ||    KN150682.1                                                              ||
## ||    KN150619.1                                                              ||
## ||    KN150397.1                                                              ||
## ||    KN150420.1                                                              ||
## ||    KN149897.1                                                              ||
## ||    KN150251.1                                                              ||
## ||    KN150082.1                                                              ||
## ||    KN149751.1                                                              ||
## ||    KN149804.1                                                              ||
## ||    KN150019.2                                                              ||
## ||    KZ115983.1                                                              ||
## ||    KN150663.1                                                              ||
## ||    KN150494.1                                                              ||
## ||    KN150547.1                                                              ||
## ||    KZ116057.1                                                              ||
## ||    KN150401.1                                                              ||
## ||    KN149878.1                                                              ||
## ||    KN150232.1                                                              ||
## ||    KN149901.1                                                              ||
## ||    KN150063.1                                                              ||
## ||    KN149732.1                                                              ||
## ||    KZ115964.1                                                              ||
## ||    KN150591.1                                                              ||
## ||    KN150644.1                                                              ||
## ||    KN150475.1                                                              ||
## ||    KN150528.1                                                              ||
## ||    KZ116038.1                                                              ||
## ||    KN149859.1                                                              ||
## ||    KN150160.1                                                              ||
## ||    KN150213.1                                                              ||
## ||    KN150044.1                                                              ||
## ||    KN149713.1                                                              ||
## ||    KN147552.2                                                              ||
## ||    KN150572.1                                                              ||
## ||    KN150625.1                                                              ||
## ||    KN150509.1                                                              ||
## ||    KN150287.1                                                              ||
## ||    KN149956.1                                                              ||
## ||    KZ116019.1                                                              ||
## ||    KN150310.1                                                              ||
## ||    KN149787.1                                                              ||
## ||    KN150141.1                                                              ||
## ||    KN149810.1                                                              ||
## ||    KN150025.1                                                              ||
## ||    KN148038.2                                                              ||
## ||    KN150699.1                                                              ||
## ||    KN150553.1                                                              ||
## ||    KN150606.1                                                              ||
## ||    KZ116063.1                                                              ||
## ||    KN150384.1                                                              ||
## ||    KN150437.1                                                              ||
## ||    KN150268.1                                                              ||
## ||    KN149884.1                                                              ||
## ||    KN149937.1                                                              ||
## ||    KN149768.1                                                              ||
## ||    KN150122.1                                                              ||
## ||    KN150006.1                                                              ||
## ||    KZ115970.1                                                              ||
## ||    KN150650.1                                                              ||
## ||    KN150703.1                                                              ||
## ||    KN150481.1                                                              ||
## ||    KN150534.1                                                              ||
## ||    KN150365.1                                                              ||
## ||    KN149981.1                                                              ||
## ||    KZ116044.1                                                              ||
## ||    KN150418.1                                                              ||
## ||    KN150196.1                                                              ||
## ||    KN150249.1                                                              ||
## ||    KN149865.1                                                              ||
## ||    KN149696.2                                                              ||
## ||    KN149749.1                                                              ||
## ||    KN150103.1                                                              ||
## ||    KZ115951.1                                                              ||
## ||    KN150631.1                                                              ||
## ||    KN150462.1                                                              ||
## ||    KN150515.1                                                              ||
## ||    KN150293.1                                                              ||
## ||    KZ116025.1                                                              ||
## ||    KN149793.1                                                              ||
## ||    KN150177.1                                                              ||
## ||    KN149846.1                                                              ||
## ||    KN150200.1                                                              ||
## ||    KN150031.1                                                              ||
## ||    KN149700.1                                                              ||
## ||    KN150589.1                                                              ||
## ||    KN150390.1                                                              ||
## ||    KN150443.1                                                              ||
## ||    KN149890.1                                                              ||
## ||    KN150274.1                                                              ||
## ||    KZ116006.1                                                              ||
## ||    KN150327.1                                                              ||
## ||    KN149943.1                                                              ||
## ||    KN149774.1                                                              ||
## ||    KN150158.1                                                              ||
## ||    KN149827.1                                                              ||
## ||    KN150012.1                                                              ||
## ||    KN150540.1                                                              ||
## ||    KZ116050.1                                                              ||
## ||    KN150371.1                                                              ||
## ||    KN149871.1                                                              ||
## ||    KN150255.2                                                              ||
## ||    KN150308.1                                                              ||
## ||    KN149924.1                                                              ||
## ||    KN150086.1                                                              ||
## ||    KN149755.1                                                              ||
## ||    KZ115987.1                                                              ||
## ||    KN150667.1                                                              ||
## ||    KN150521.1                                                              ||
## ||    KN149998.1                                                              ||
## ||    KZ116031.1                                                              ||
## ||    KN150352.2                                                              ||
## ||    KN150405.1                                                              ||
## ||    KN150183.1                                                              ||
## ||    KN150236.1                                                              ||
## ||    KN149852.1                                                              ||
## ||    KN149905.1                                                              ||
## ||    KN150067.1                                                              ||
## ||    KN149683.1                                                              ||
## ||    KZ115968.1                                                              ||
## ||    KN150595.1                                                              ||
## ||    KN150479.1                                                              ||
## ||    KN150502.1                                                              ||
## ||    KN149979.1                                                              ||
## ||    KN150280.1                                                              ||
## ||    KZ116012.1                                                              ||
## ||    KN150333.1                                                              ||
## ||    KN150164.1                                                              ||
## ||    KN149780.1                                                              ||
## ||    KN150217.1                                                              ||
## ||    KN150048.1                                                              ||
## ||    KN149717.1                                                              ||
## ||    KZ115949.1                                                              ||
## ||    KN150576.1                                                              ||
## ||    KN150629.1                                                              ||
## ||    KN150430.1                                                              ||
## ||    KN150314.1                                                              ||
## ||    KN150092.1                                                              ||
## ||    KN150145.1                                                              ||
## ||    KN149761.1                                                              ||
## ||    KN149814.1                                                              ||
## ||    KN150029.1                                                              ||
## ||    KZ115993.1                                                              ||
## ||    KN150557.1                                                              ||
## ||    KN150388.1                                                              ||
## ||    KZ116067.1                                                              ||
## ||    KN150411.1                                                              ||
## ||    KN149888.1                                                              ||
## ||    KN150242.1                                                              ||
## ||    KN150073.1                                                              ||
## ||    KN149742.1                                                              ||
## ||    KN150126.1                                                              ||
## ||    KZ115974.1                                                              ||
## ||    KN150654.1                                                              ||
## ||    KN150485.1                                                              ||
## ||    KN150538.1                                                              ||
## ||    KZ116048.1                                                              ||
## ||    KN150369.1                                                              ||
## ||    KN150170.1                                                              ||
## ||    KN150054.1                                                              ||
## ||    KN149723.1                                                              ||
## ||    KZ115955.1                                                              ||
## ||    KN150582.1                                                              ||
## ||    KN150635.1                                                              ||
## ||    KN150466.1                                                              ||
## ||    KN150519.1                                                              ||
## ||    KN150297.1                                                              ||
## ||    KZ116029.1                                                              ||
## ||    KN150320.1                                                              ||
## ||    KN149797.1                                                              ||
## ||    KN150204.1                                                              ||
## ||    KN150035.1                                                              ||
## ||    KN150616.1                                                              ||
## ||    KN150394.1                                                              ||
## ||    KN150278.1                                                              ||
## ||    KN150301.1                                                              ||
## ||    KN149778.1                                                              ||
## ||    KN150132.1                                                              ||
## ||    KN149801.1                                                              ||
## ||    KN150016.1                                                              ||
## ||    KZ115980.1                                                              ||
## ||    KN150713.1                                                              ||
## ||    KN150491.1                                                              ||
## ||    KN150544.1                                                              ||
## ||    KN149991.1                                                              ||
## ||    KZ116054.1                                                              ||
## ||    KN150375.1                                                              ||
## ||    KN150428.1                                                              ||
## ||    KN150259.1                                                              ||
## ||    KN149928.1                                                              ||
## ||    KN150060.1                                                              ||
## ||    KN150113.1                                                              ||
## ||    KZ115961.1                                                              ||
## ||    KN150641.1                                                              ||
## ||    KN150472.1                                                              ||
## ||    KN150525.1                                                              ||
## ||    KZ116035.1                                                              ||
## ||    KN150187.1                                                              ||
## ||    KN149856.1                                                              ||
## ||    KN149909.1                                                              ||
## ||    KN150210.1                                                              ||
## ||    KN149687.1                                                              ||
## ||    KN150041.2                                                              ||
## ||    KN149710.1                                                              ||
## ||    KN150599.1                                                              ||
## ||    KN150622.1                                                              ||
## ||    KN150453.1                                                              ||
## ||    KN150506.1                                                              ||
## ||    KN150284.1                                                              ||
## ||    KZ116016.1                                                              ||
## ||    KN150337.1                                                              ||
## ||    KN149953.1                                                              ||
## ||    KN150168.1                                                              ||
## ||    KN149784.1                                                              ||
## ||    KN149837.1                                                              ||
## ||    KN150022.1                                                              ||
## ||    KN150550.1                                                              ||
## ||    KN150603.1                                                              ||
## ||    KZ116060.1                                                              ||
## ||    KN150434.1                                                              ||
## ||    KN149881.1                                                              ||
## ||    KN150265.1                                                              ||
## ||    KN150318.1                                                              ||
## ||    KN149765.1                                                              ||
## ||    KN149818.1                                                              ||
## ||    KZ115997.1                                                              ||
## ||    KN150677.1                                                              ||
## ||    KN150700.2                                                              ||
## ||    KZ116041.1                                                              ||
## ||    KN150362.1                                                              ||
## ||    KN150193.1                                                              ||
## ||    KN149862.1                                                              ||
## ||    KN150246.1                                                              ||
## ||    KN149915.1                                                              ||
## ||    KN149693.1                                                              ||
## ||    KN150077.1                                                              ||
## ||    KN149746.1                                                              ||
## ||    KN150100.1                                                              ||
## ||    KZ115978.1                                                              ||
## ||    KN150489.1                                                              ||
## ||    KN149989.1                                                              ||
## ||    KN150290.1                                                              ||
## ||    KZ116022.1                                                              ||
## ||    KN150343.1                                                              ||
## ||    KN150174.1                                                              ||
## ||    KN149790.1                                                              ||
## ||    KN150227.1                                                              ||
## ||    KN149843.1                                                              ||
## ||    KN150058.1                                                              ||
## ||    KN149727.1                                                              ||
## ||    KZ115959.1                                                              ||
## ||    KN150586.1                                                              ||
## ||    KN150639.1                                                              ||
## ||    KN150440.1                                                              ||
## ||    KN150271.1                                                              ||
## ||    KN149940.1                                                              ||
## ||    KZ116003.1                                                              ||
## ||    KN150324.1                                                              ||
## ||    KN150155.1                                                              ||
## ||    KN149771.1                                                              ||
## ||    KN149824.1                                                              ||
## ||    KN150039.1                                                              ||
## ||    KN149708.1                                                              ||
## ||    KN150683.1                                                              ||
## ||    KN150567.1                                                              ||
## ||    KN150398.1                                                              ||
## ||    KN150421.1                                                              ||
## ||    KN150252.1                                                              ||
## ||    KN149921.1                                                              ||
## ||    KN150083.1                                                              ||
## ||    KN150136.1                                                              ||
## ||    KN149805.1                                                              ||
## ||    KZ115984.1                                                              ||
## ||    KN150664.1                                                              ||
## ||    KN150495.1                                                              ||
## ||    KN150548.1                                                              ||
## ||    KZ116058.1                                                              ||
## ||    KN149995.1                                                              ||
## ||    KN150379.1                                                              ||
## ||    KN149879.1                                                              ||
## ||    KN150180.1                                                              ||
## ||    KN150233.1                                                              ||
## ||    KN150064.1                                                              ||
## ||    KN149680.1                                                              ||
## ||    KN149733.1                                                              ||
## ||    KN150117.1                                                              ||
## ||    KZ115965.1                                                              ||
## ||    KN150592.1                                                              ||
## ||    KN150645.1                                                              ||
## ||    KN150476.1                                                              ||
## ||    KN150529.1                                                              ||
## ||    KN149976.1                                                              ||
## ||    KZ116039.1                                                              ||
## ||    KN150330.1                                                              ||
## ||    KN150161.1                                                              ||
## ||    KN150214.1                                                              ||
## ||    KN150045.2                                                              ||
## ||    KN149714.1                                                              ||
## ||    KN150573.1                                                              ||
## ||    KN150626.1                                                              ||
## ||    KN149957.1                                                              ||
## ||    KN150311.1                                                              ||
## ||    KN149788.1                                                              ||
## ||    KN149811.1                                                              ||
## ||    KZ115990.1                                                              ||
## ||    KN150670.1                                                              ||
## ||    KN150554.1                                                              ||
## ||    KN150607.1                                                              ||
## ||    KN150385.1                                                              ||
## ||    KZ116064.1                                                              ||
## ||    KN149885.1                                                              ||
## ||    KN150269.1                                                              ||
## ||    KN149938.1                                                              ||
## ||    KN149769.1                                                              ||
## ||    KN150070.1                                                              ||
## ||    KN150123.1                                                              ||
## ||    KN150007.1                                                              ||
## ||    KZ115971.1                                                              ||
## ||    KN150651.1                                                              ||
## ||    KN150535.1                                                              ||
## ||    KZ116045.1                                                              ||
## ||    KN150366.1                                                              ||
## ||    KN149982.1                                                              ||
## ||    KN150419.1                                                              ||
## ||    KN150197.1                                                              ||
## ||    KN149866.1                                                              ||
## ||    KN149919.1                                                              ||
## ||    KN150220.1                                                              ||
## ||    KN149697.1                                                              ||
## ||    KN150051.1                                                              ||
## ||    KN149720.1                                                              ||
## ||    KN150104.1                                                              ||
## ||    KZ115952.1                                                              ||
## ||    KN150632.1                                                              ||
## ||    KN150516.1                                                              ||
## ||    KZ116026.1                                                              ||
## ||    KN150347.1                                                              ||
## ||    KN149794.1                                                              ||
## ||    KN149847.1                                                              ||
## ||    KN150201.1                                                              ||
## ||    KN150032.1                                                              ||
## ||    KN149701.1                                                              ||
## ||    KN150560.1                                                              ||
## ||    KN150613.1                                                              ||
## ||    KN150444.1                                                              ||
## ||    KN150275.1                                                              ||
## ||    KZ116007.1                                                              ||
## ||    KN150328.1                                                              ||
## ||    KN149944.1                                                              ||
## ||    KN150159.1                                                              ||
## ||    KN149775.2                                                              ||
## ||    KN149828.1                                                              ||
## ||    KN150013.1                                                              ||
## ||    KN150687.1                                                              ||
## ||    KN148869.2                                                              ||
## ||    KN150710.1                                                              ||
## ||    KN150541.1                                                              ||
## ||    KZ116051.1                                                              ||
## ||    KN150372.1                                                              ||
## ||    KN150425.1                                                              ||
## ||    KN149872.1                                                              ||
## ||    KN150309.1                                                              ||
## ||    KN149925.1                                                              ||
## ||    KN149756.1                                                              ||
## ||    KN149809.1                                                              ||
## ||    KN150110.1                                                              ||
## ||    KZ115988.1                                                              ||
## ||    KN150668.1                                                              ||
## ||    KN150499.1                                                              ||
## ||    KN150522.1                                                              ||
## ||    KN149999.1                                                              ||
## ||    KZ116032.1                                                              ||
## ||    KN150353.1                                                              ||
## ||    KN150406.1                                                              ||
## ||    KN150184.1                                                              ||
## ||    KN150237.1                                                              ||
## ||    KN149853.1                                                              ||
## ||    KN149906.1                                                              ||
## ||    KN150068.1                                                              ||
## ||    KN149684.1                                                              ||
## ||    KN149737.1                                                              ||
## ||    KZ115969.1                                                              ||
## ||    KN150596.1                                                              ||
## ||    KN150649.1                                                              ||
## ||    KN150450.1                                                              ||
## ||    KN150503.1                                                              ||
## ||    KN149950.1                                                              ||
## ||    KZ116013.1                                                              ||
## ||    KN150334.1                                                              ||
## ||    KN150165.1                                                              ||
## ||    KN150218.1                                                              ||
## ||    KN149834.1                                                              ||
## ||    KN150049.1                                                              ||
## ||    KN149718.1                                                              ||
## ||    KN150577.1                                                              ||
## ||    KN150431.1                                                              ||
## ||    KN149931.1                                                              ||
## ||    KN150315.1                                                              ||
## ||    KN150093.1                                                              ||
## ||    KN150146.1                                                              ||
## ||    KN149762.1                                                              ||
## ||    KN149815.1                                                              ||
## ||    KN150000.1                                                              ||
## ||    KZ115994.1                                                              ||
## ||    KN150558.1                                                              ||
## ||    KN150389.1                                                              ||
## ||    KN150412.1                                                              ||
## ||    KN149889.1                                                              ||
## ||    KN150190.1                                                              ||
## ||    KN150243.2                                                              ||
## ||    KN149912.1                                                              ||
## ||    KN150074.1                                                              ||
## ||    KN149690.1                                                              ||
## ||    KN149743.1                                                              ||
## ||    KN150127.1                                                              ||
## ||    KZ115975.1                                                              ||
## ||    KN150655.1                                                              ||
## ||    KN150708.1                                                              ||
## ||    KN150486.1                                                              ||
## ||    KZ116049.1                                                              ||
## ||    KN150340.1                                                              ||
## ||    KN150171.1                                                              ||
## ||    KN150224.1                                                              ||
## ||    KN149840.1                                                              ||
## ||    KN150055.1                                                              ||
## ||    KN149724.1                                                              ||
## ||    KZ115956.1                                                              ||
## ||    KN150583.1                                                              ||
## ||    KN150636.1                                                              ||
## ||    KN150467.1                                                              ||
## ||    KN149967.1                                                              ||
## ||    KZ116000.1                                                              ||
## ||    KN149798.1                                                              ||
## ||    KN150152.1                                                              ||
## ||    KN150205.1                                                              ||
## ||    KN149705.1                                                              ||
## ||    KN150680.1                                                              ||
## ||    KN150564.1                                                              ||
## ||    KN150617.1                                                              ||
## ||    KN150395.1                                                              ||
## ||    KN150279.1                                                              ||
## ||    KN149895.1                                                              ||
## ||    KN149948.1                                                              ||
## ||    KN150302.1                                                              ||
## ||    KN149779.1                                                              ||
## ||    KN150080.1                                                              ||
## ||    KN150133.2                                                              ||
## ||    KN150017.1                                                              ||
## ||    KZ115981.1                                                              ||
## ||    KN150661.1                                                              ||
## ||    KN150492.2                                                              ||
## ||    KN150545.1                                                              ||
## ||    KZ116055.1                                                              ||
## ||    KN149992.1                                                              ||
## ||    KN150429.1                                                              ||
## ||    KN149876.1                                                              ||
## ||    KN150230.1                                                              ||
## ||    KN150061.1                                                              ||
## ||    KN149730.1                                                              ||
## ||    KN150114.1                                                              ||
## ||    KZ115962.1                                                              ||
## ||    KN150642.1                                                              ||
## ||    KN150473.1                                                              ||
## ||    KN150526.1                                                              ||
## ||    KZ116036.1                                                              ||
## ||    KN150357.1                                                              ||
## ||    KN149973.1                                                              ||
## ||    KN149857.1                                                              ||
## ||    KN150211.1                                                              ||
## ||    KN149688.2                                                              ||
## ||    KN150042.1                                                              ||
## ||    KN149711.1                                                              ||
## ||    KN150570.1                                                              ||
## ||    KN150623.1                                                              ||
## ||    KN150454.1                                                              ||
## ||    KN150285.1                                                              ||
## ||    KN150338.1                                                              ||
## ||    KZ116017.1                                                              ||
## ||    KN149785.1                                                              ||
## ||    KN149838.1                                                              ||
## ||    KN150023.1                                                              ||
## ||    KN150697.1                                                              ||
## ||    KN150551.1                                                              ||
## ||    KZ116061.1                                                              ||
## ||    KN150382.1                                                              ||
## ||    KN149882.1                                                              ||
## ||    KN149935.1                                                              ||
## ||    KN150319.1                                                              ||
## ||    KN150097.1                                                              ||
## ||    KN149766.1                                                              ||
## ||    KN149819.1                                                              ||
## ||    KN150120.1                                                              ||
## ||    KN150004.2                                                              ||
## ||    KZ115998.1                                                              ||
## ||    KN150678.1                                                              ||
## ||    KN150701.1                                                              ||
## ||    KN150532.1                                                              ||
## ||    KZ116042.1                                                              ||
## ||    KN150363.1                                                              ||
## ||    KN150416.1                                                              ||
## ||    KN150194.1                                                              ||
## ||    KN149863.1                                                              ||
## ||    KN149916.1                                                              ||
## ||    KN150078.1                                                              ||
## ||    KN149694.1                                                              ||
## ||    KN149747.1                                                              ||
## ||    KZ115979.1                                                              ||
## ||    KN150659.1                                                              ||
## ||    KN150513.1                                                              ||
## ||    KN150291.1                                                              ||
## ||    KZ116023.1                                                              ||
## ||    KN149960.1                                                              ||
## ||    KN150344.1                                                              ||
## ||    KN149791.1                                                              ||
## ||    KN150175.1                                                              ||
## ||    KN149844.1                                                              ||
## ||    KN150228.1                                                              ||
## ||    KN150059.1                                                              ||
## ||    KN149728.1                                                              ||
## ||    KN150587.1                                                              ||
## ||    KN150610.1                                                              ||
## ||    KN150441.1                                                              ||
## ||    KN150272.1                                                              ||
## ||    KN149941.1                                                              ||
## ||    KZ116004.1                                                              ||
## ||    KN150325.1                                                              ||
## ||    KN149772.1                                                              ||
## ||    KN150156.1                                                              ||
## ||    KN149825.1                                                              ||
## ||    KN149709.1                                                              ||
## ||    KN150684.1                                                              ||
## ||    KN150568.1                                                              ||
## ||    KN150399.1                                                              ||
## ||    KN150422.1                                                              ||
## ||    KN149899.1                                                              ||
## ||    KN150253.1                                                              ||
## ||    KN149922.1                                                              ||
## ||    KN150306.1                                                              ||
## ||    KN150084.1                                                              ||
## ||    KN149753.1                                                              ||
## ||    KN150137.1                                                              ||
## ||    KN149806.1                                                              ||
## ||    KZ115985.1                                                              ||
## ||                                                                            ||
## || Global environment is initialised.                                         ||
## || Load the 1-th index block...                                               ||
## || The index block has been loaded.                                           ||
## || Start read mapping in chunk.                                               ||
## ||    0% completed, 0.1 mins elapsed, rate=73.7k reads per second             ||
## ||    6% completed, 0.4 mins elapsed, rate=84.4k reads per second             ||
## ||   13% completed, 0.8 mins elapsed, rate=84.8k reads per second             ||
## ||   18% completed, 1.1 mins elapsed, rate=72.5k reads per second             ||
## ||   18% completed, 1.1 mins elapsed, rate=73.6k reads per second             ||
## ||   19% completed, 1.2 mins elapsed, rate=74.8k reads per second             ||
## ||   20% completed, 1.2 mins elapsed, rate=75.9k reads per second             ||
## ||   21% completed, 1.2 mins elapsed, rate=77.3k reads per second             ||
## ||   22% completed, 1.3 mins elapsed, rate=78.5k reads per second             ||
## ||   23% completed, 1.3 mins elapsed, rate=79.7k reads per second             ||
## ||   23% completed, 1.3 mins elapsed, rate=80.7k reads per second             ||
## ||   24% completed, 1.4 mins elapsed, rate=81.7k reads per second             ||
## ||   25% completed, 1.4 mins elapsed, rate=82.6k reads per second             ||
## || Start read mapping in chunk.                                               ||
## ||   26% completed, 1.4 mins elapsed, rate=87.7k reads per second             ||
## ||   33% completed, 1.7 mins elapsed, rate=88.7k reads per second             ||
## ||   39% completed, 2.1 mins elapsed, rate=89.7k reads per second             ||
## ||   44% completed, 2.3 mins elapsed, rate=84.4k reads per second             ||
## ||   45% completed, 2.4 mins elapsed, rate=84.8k reads per second             ||
## ||   45% completed, 2.4 mins elapsed, rate=85.2k reads per second             ||
## ||   46% completed, 2.4 mins elapsed, rate=85.6k reads per second             ||
## ||   47% completed, 2.5 mins elapsed, rate=85.9k reads per second             ||
## ||   48% completed, 2.5 mins elapsed, rate=86.3k reads per second             ||
## ||   49% completed, 2.5 mins elapsed, rate=86.6k reads per second             ||
## ||   50% completed, 2.6 mins elapsed, rate=87.0k reads per second             ||
## ||   51% completed, 2.6 mins elapsed, rate=87.3k reads per second             ||
## ||   51% completed, 2.6 mins elapsed, rate=87.6k reads per second             ||
## || Start read mapping in chunk.                                               ||
## ||   52% completed, 2.7 mins elapsed, rate=90.3k reads per second             ||
## ||   58% completed, 3.0 mins elapsed, rate=91.0k reads per second             ||
## ||   65% completed, 3.3 mins elapsed, rate=91.6k reads per second             ||
## ||   70% completed, 3.6 mins elapsed, rate=88.0k reads per second             ||
## ||   71% completed, 3.6 mins elapsed, rate=88.1k reads per second             ||
## ||   72% completed, 3.7 mins elapsed, rate=88.3k reads per second             ||
## ||   72% completed, 3.7 mins elapsed, rate=88.4k reads per second             ||
## ||   73% completed, 3.7 mins elapsed, rate=88.5k reads per second             ||
## ||   74% completed, 3.8 mins elapsed, rate=88.7k reads per second             ||
## ||   75% completed, 3.8 mins elapsed, rate=88.9k reads per second             ||
## ||   76% completed, 3.9 mins elapsed, rate=89.0k reads per second             ||
## ||   77% completed, 3.9 mins elapsed, rate=89.1k reads per second             ||
## || Start read mapping in chunk.                                               ||
## ||   78% completed, 3.9 mins elapsed, rate=90.7k reads per second             ||
## ||   84% completed, 4.3 mins elapsed, rate=91.0k reads per second             ||
## ||   91% completed, 4.6 mins elapsed, rate=91.4k reads per second             ||
## ||   93% completed, 4.7 mins elapsed, rate=88.6k reads per second             ||
## ||   94% completed, 4.8 mins elapsed, rate=88.7k reads per second             ||
## ||   94% completed, 4.8 mins elapsed, rate=88.7k reads per second             ||
## ||   95% completed, 4.8 mins elapsed, rate=88.7k reads per second             ||
## ||   96% completed, 4.9 mins elapsed, rate=88.8k reads per second             ||
## ||   96% completed, 4.9 mins elapsed, rate=88.8k reads per second             ||
## ||   97% completed, 4.9 mins elapsed, rate=88.9k reads per second             ||
## ||   98% completed, 5.0 mins elapsed, rate=88.9k reads per second             ||
## ||   99% completed, 5.0 mins elapsed, rate=89.0k reads per second             ||
## ||   99% completed, 5.0 mins elapsed, rate=89.0k reads per second             ||
## ||                                                                            ||
## ||                           Completed successfully.                          ||
## ||                                                                            ||
## \\====================================    ====================================//
## 
## //================================   Summary =================================\\
## ||                                                                            ||
## ||                 Total reads : 26,877,266                                   ||
## ||                      Mapped : 25,392,455 (94.5%)                           ||
## ||             Uniquely mapped : 16,064,116                                   ||
## ||               Multi-mapping : 9,328,339                                    ||
## ||                                                                            ||
## ||                    Unmapped : 1,484,811                                    ||
## ||                                                                            ||
## ||                   Junctions : 135,968                                      ||
## ||                      Indels : 68,092                                       ||
## ||                                                                            ||
## ||                Running time : 5.0 minutes                                  ||
## ||                                                                            ||
## \\============================================================================//
## 
## 
##         ==========     _____ _    _ ____  _____  ______          _____  
##         =====         / ____| |  | |  _ \|  __ \|  ____|   /\   |  __ \ 
##           =====      | (___ | |  | | |_) | |__) | |__     /  \  | |  | |
##             ====      \___ \| |  | |  _ <|  _  /|  __|   / /\ \ | |  | |
##               ====    ____) | |__| | |_) | | \ \| |____ / ____ \| |__| |
##         ==========   |_____/ \____/|____/|_|  \_\______/_/    \_\_____/
##        Rsubread 2.0.1
## 
## //================================= setting ==================================\\
## ||                                                                            ||
## || Function      : Read alignment + Junction detection (RNA-Seq)              ||
## || Input file    : 9499X4_130405_SN141_0668_AD2154ACXX_7.txt.gz               ||
## || Output file   : 9499X4.bam (BAM)                                           ||
## || Index name    : GRCz11                                                     ||
## ||                                                                            ||
## ||                    ------------------------------------                    ||
## ||                                                                            ||
## ||                               Threads : 8                                  ||
## ||                          Phred offset : 33                                 ||
## ||                             Min votes : 1 / 14                             ||
## ||                        Max mismatches : 3                                  ||
## ||                      Max indel length : 5                                  ||
## ||            Report multi-mapping reads : yes                                ||
## || Max alignments per multi-mapping read : 1                                  ||
## ||                           Annotations : Danio_rerio.GRCz11.93.chr.gtf  ... ||
## ||                                                                            ||
## \\============================================================================//
## 
## //================ Running (24-Jun-2020 11:08:09, pid=12013) =================\\
## ||                                                                            ||
## || Check the input reads.                                                     ||
## || The input file contains base space reads.                                  ||
## || Initialise the memory objects.                                             ||
## || Estimate the mean read length.                                             ||
## || The range of Phred scores observed in the data is [2,41]                   ||
## || Create the output BAM file.                                                ||
## || Check the index.                                                           ||
## || Init the voting space.                                                     ||
## || Load the annotation file.                                                  ||
## || 481079 annotation records were loaded.                                     ||
## ||                                                                            ||
## || Chromosomes/contigs in index but not in annotation :                       ||
## ||    KN150665.1                                                              ||
## ||    KN150549.1                                                              ||
## ||    KN149996.1                                                              ||
## ||    KZ116059.1                                                              ||
## ||    KN150350.1                                                              ||
## ||    KN150403.1                                                              ||
## ||    KN150181.1                                                              ||
## ||    KN150234.1                                                              ||
## ||    KN149850.1                                                              ||
## ||    KN149903.1                                                              ||
## ||    KN150065.1                                                              ||
## ||    KN150118.1                                                              ||
## ||    KZ115966.1                                                              ||
## ||    KN150593.1                                                              ||
## ||    KN150646.1                                                              ||
## ||    KN148828.2                                                              ||
## ||    KN150477.1                                                              ||
## ||    KN150500.1                                                              ||
## ||    KN149977.2                                                              ||
## ||    KZ116010.1                                                              ||
## ||    KN150331.1                                                              ||
## ||    KN150162.1                                                              ||
## ||    KN150215.1                                                              ||
## ||    KN149715.1                                                              ||
## ||    KN150690.1                                                              ||
## ||    KN150574.1                                                              ||
## ||    KN150458.1                                                              ||
## ||    KN150289.1                                                              ||
## ||    KN149958.1                                                              ||
## ||    KN150312.1                                                              ||
## ||    KN149789.1                                                              ||
## ||    KN150090.1                                                              ||
## ||    KN149812.1                                                              ||
## ||    KN147651.2                                                              ||
## ||    KZ115991.1                                                              ||
## ||    KN150671.1                                                              ||
## ||    KN150555.1                                                              ||
## ||    KN150608.1                                                              ||
## ||    KN150386.1                                                              ||
## ||    KZ116065.1                                                              ||
## ||    KN150439.1                                                              ||
## ||    KN149939.1                                                              ||
## ||    KN150240.1                                                              ||
## ||    KN150071.1                                                              ||
## ||    KN149740.1                                                              ||
## ||    KN150008.1                                                              ||
## ||    KN147632.2                                                              ||
## ||    KZ115972.1                                                              ||
## ||    KN150652.1                                                              ||
## ||    KN150483.1                                                              ||
## ||    KN150536.1                                                              ||
## ||    KZ116046.1                                                              ||
## ||    KN149983.1                                                              ||
## ||    KN149867.1                                                              ||
## ||    KN150221.1                                                              ||
## ||    KN149698.1                                                              ||
## ||    KN150052.1                                                              ||
## ||    KN149721.1                                                              ||
## ||    KN150105.1                                                              ||
## ||    KZ115953.1                                                              ||
## ||    KN150580.1                                                              ||
## ||    KN150633.1                                                              ||
## ||    KN150464.1                                                              ||
## ||    KN150517.1                                                              ||
## ||    KZ116027.1                                                              ||
## ||    KN149795.1                                                              ||
## ||    KN150179.1                                                              ||
## ||    KN149848.1                                                              ||
## ||    KN150202.1                                                              ||
## ||    KN150033.1                                                              ||
## ||    KN149702.1                                                              ||
## ||    KN150561.1                                                              ||
## ||    KN150614.1                                                              ||
## ||    KN150392.1                                                              ||
## ||    KN150445.1                                                              ||
## ||    KN150276.1                                                              ||
## ||    KN149892.1                                                              ||
## ||    KN150329.1                                                              ||
## ||    KZ116008.1                                                              ||
## ||    KN149829.1                                                              ||
## ||    KN150130.1                                                              ||
## ||    KN150014.1                                                              ||
## ||    KN150711.1                                                              ||
## ||    KN150542.1                                                              ||
## ||    KZ116052.1                                                              ||
## ||    KN150426.1                                                              ||
## ||    KN149873.1                                                              ||
## ||    KN150257.1                                                              ||
## ||    KN149926.1                                                              ||
## ||    KN149757.1                                                              ||
## ||    KN150111.1                                                              ||
## ||    KZ115989.1                                                              ||
## ||    KN150669.1                                                              ||
## ||    KN150470.1                                                              ||
## ||    KN150523.1                                                              ||
## ||    KN150354.1                                                              ||
## ||    KN149970.1                                                              ||
## ||    KZ116033.1                                                              ||
## ||    KN150407.1                                                              ||
## ||    KN150185.1                                                              ||
## ||    KN149854.1                                                              ||
## ||    KN150238.1                                                              ||
## ||    KN149907.1                                                              ||
## ||    KN149685.1                                                              ||
## ||    KN149738.1                                                              ||
## ||    KN150620.1                                                              ||
## ||    KN150451.1                                                              ||
## ||    KN150504.1                                                              ||
## ||    KN150282.1                                                              ||
## ||    KZ116014.1                                                              ||
## ||    KN149951.1                                                              ||
## ||    KN150335.1                                                              ||
## ||    KN149782.1                                                              ||
## ||    KN150166.1                                                              ||
## ||    KN150219.1                                                              ||
## ||    KN149835.1                                                              ||
## ||    KN149719.1                                                              ||
## ||    KN150020.1                                                              ||
## ||    KN150578.1                                                              ||
## ||    KN150601.1                                                              ||
## ||    KN150432.1                                                              ||
## ||    KN150263.1                                                              ||
## ||    KN149932.1                                                              ||
## ||    KN150094.1                                                              ||
## ||    KN150147.1                                                              ||
## ||    KN149763.1                                                              ||
## ||    KN149816.1                                                              ||
## ||    KZ115995.1                                                              ||
## ||    KN150675.1                                                              ||
## ||    KN150559.1                                                              ||
## ||    KN150360.1                                                              ||
## ||    KN150413.1                                                              ||
## ||    KN150191.1                                                              ||
## ||    KN149860.1                                                              ||
## ||    KN150244.1                                                              ||
## ||    KN149913.1                                                              ||
## ||    KN149691.1                                                              ||
## ||    KN150128.1                                                              ||
## ||    KN147636.1                                                              ||
## ||    KZ115976.1                                                              ||
## ||    KN150656.2                                                              ||
## ||    KN150709.1                                                              ||
## ||    KN150487.1                                                              ||
## ||    KN150510.1                                                              ||
## ||    KZ116020.1                                                              ||
## ||    KN150341.1                                                              ||
## ||    KN150172.1                                                              ||
## ||    KN149841.1                                                              ||
## ||    KN150225.1                                                              ||
## ||    KN150056.1                                                              ||
## ||    KN149725.1                                                              ||
## ||    KZ115957.1                                                              ||
## ||    KN150468.1                                                              ||
## ||    KN150299.1                                                              ||
## ||    KN149968.1                                                              ||
## ||    KZ116001.1                                                              ||
## ||    KN150322.1                                                              ||
## ||    KN150206.1                                                              ||
## ||    KN149822.1                                                              ||
## ||    KN150037.1                                                              ||
## ||    KN149706.1                                                              ||
## ||    KN150565.1                                                              ||
## ||    KN150396.1                                                              ||
## ||    KN150449.1                                                              ||
## ||    KN149896.1                                                              ||
## ||    KN149949.1                                                              ||
## ||    KN150250.1                                                              ||
## ||    KN150303.1                                                              ||
## ||    KN150081.1                                                              ||
## ||    KN149750.1                                                              ||
## ||    KN150134.1                                                              ||
## ||    KN149803.1                                                              ||
## ||    KN150018.1                                                              ||
## ||    KN147642.2                                                              ||
## ||    KZ115982.1                                                              ||
## ||    KN150662.1                                                              ||
## ||    KN150493.1                                                              ||
## ||    KN150546.1                                                              ||
## ||    KZ116056.1                                                              ||
## ||    KN149993.1                                                              ||
## ||    KN150400.1                                                              ||
## ||    KN150231.1                                                              ||
## ||    KN149900.1                                                              ||
## ||    KN150062.1                                                              ||
## ||    KN150115.1                                                              ||
## ||    KZ115963.1                                                              ||
## ||    KN150643.1                                                              ||
## ||    KN150527.1                                                              ||
## ||    KN150358.1                                                              ||
## ||    KZ116037.1                                                              ||
## ||    KN149974.1                                                              ||
## ||    KN150189.1                                                              ||
## ||    KN150212.1                                                              ||
## ||    KN149689.2                                                              ||
## ||    KN150043.1                                                              ||
## ||    KN150571.1                                                              ||
## ||    KN150455.1                                                              ||
## ||    KN150508.1                                                              ||
## ||    KN150286.1                                                              ||
## ||    KZ116018.1                                                              ||
## ||    KN149955.1                                                              ||
## ||    KN149786.1                                                              ||
## ||    KN149839.1                                                              ||
## ||    KN150140.2                                                              ||
## ||    KN150024.1                                                              ||
## ||    KN150698.1                                                              ||
## ||    KN150552.1                                                              ||
## ||    KN150605.1                                                              ||
## ||    KZ116062.1                                                              ||
## ||    KN150383.1                                                              ||
## ||    KN150436.1                                                              ||
## ||    KN149883.1                                                              ||
## ||    KN150267.1                                                              ||
## ||    KN149936.1                                                              ||
## ||    KN150098.1                                                              ||
## ||    KN149767.1                                                              ||
## ||    KN150121.1                                                              ||
## ||    KN150005.1                                                              ||
## ||    KZ115999.1                                                              ||
## ||    KN150679.1                                                              ||
## ||    KN149980.1                                                              ||
## ||    KN150364.1                                                              ||
## ||    KZ116043.1                                                              ||
## ||    KN150195.1                                                              ||
## ||    KN149864.1                                                              ||
## ||    KN150248.1                                                              ||
## ||    KN149917.1                                                              ||
## ||    KN150079.1                                                              ||
## ||    KN149695.1                                                              ||
## ||    KN149748.1                                                              ||
## ||    KN150102.1                                                              ||
## ||    KZ115950.1                                                              ||
## ||    KN150630.1                                                              ||
## ||    KN150514.1                                                              ||
## ||    KN150292.1                                                              ||
## ||    KN150345.1                                                              ||
## ||    KN149961.1                                                              ||
## ||    KZ116024.1                                                              ||
## ||    KN149792.1                                                              ||
## ||    KN150176.1                                                              ||
## ||    KN150229.1                                                              ||
## ||    KN149729.1                                                              ||
## ||    KN150030.1                                                              ||
## ||    KN150588.1                                                              ||
## ||    KZ116005.1                                                              ||
## ||    KN150326.1                                                              ||
## ||    KN150157.1                                                              ||
## ||    KN149826.1                                                              ||
## ||    KN150011.1                                                              ||
## ||    KN150685.1                                                              ||
## ||    KN150254.1                                                              ||
## ||    KN150307.1                                                              ||
## ||    KN149923.1                                                              ||
## ||    KN150085.1                                                              ||
## ||    KN149754.1                                                              ||
## ||    KN149807.1                                                              ||
## ||    KZ115986.1                                                              ||
## ||    KN150666.1                                                              ||
## ||    KN150520.1                                                              ||
## ||    KN149997.1                                                              ||
## ||    KZ116030.1                                                              ||
## ||    KN150351.1                                                              ||
## ||    KN150404.1                                                              ||
## ||    KN150182.1                                                              ||
## ||    KN149851.1                                                              ||
## ||    KN149904.1                                                              ||
## ||    KN150066.1                                                              ||
## ||    KN149682.1                                                              ||
## ||    KN150119.1                                                              ||
## ||    KN149735.1                                                              ||
## ||    KZ115967.1                                                              ||
## ||    KN150594.1                                                              ||
## ||    KN150647.1                                                              ||
## ||    KN150478.1                                                              ||
## ||    KN150501.1                                                              ||
## ||    KN149978.1                                                              ||
## ||    KN150332.1                                                              ||
## ||    KZ116011.1                                                              ||
## ||    KN150216.1                                                              ||
## ||    KN149832.1                                                              ||
## ||    KN150047.1                                                              ||
## ||    KN149716.1                                                              ||
## ||    KN150691.1                                                              ||
## ||    KZ115948.1                                                              ||
## ||    KN150575.1                                                              ||
## ||    KN150628.1                                                              ||
## ||    KN150459.1                                                              ||
## ||    KN149959.1                                                              ||
## ||    KN150260.1                                                              ||
## ||    KN150313.1                                                              ||
## ||    KN150144.1                                                              ||
## ||    KN149760.1                                                              ||
## ||    KN149813.1                                                              ||
## ||    KN150028.1                                                              ||
## ||    KN147652.2                                                              ||
## ||    KZ115992.1                                                              ||
## ||    KN150556.1                                                              ||
## ||    KN150609.1                                                              ||
## ||    KN150387.1                                                              ||
## ||    KZ116066.1                                                              ||
## ||    KN149887.1                                                              ||
## ||    KN150241.1                                                              ||
## ||    KN150072.1                                                              ||
## ||    KN149741.1                                                              ||
## ||    KN150125.1                                                              ||
## ||    KN150009.1                                                              ||
## ||    KZ115973.1                                                              ||
## ||    KN150653.1                                                              ||
## ||    KN150484.1                                                              ||
## ||    KN150537.1                                                              ||
## ||    KZ116047.1                                                              ||
## ||    KN150368.1                                                              ||
## ||    KN149984.1                                                              ||
## ||    KN150199.1                                                              ||
## ||    KN150222.1                                                              ||
## ||    KN149722.1                                                              ||
## ||    KN150106.1                                                              ||
## ||    KZ115954.1                                                              ||
## ||    KN150581.1                                                              ||
## ||    KN150634.1                                                              ||
## ||    KN150465.1                                                              ||
## ||    KN150518.1                                                              ||
## ||    KN150296.1                                                              ||
## ||    KZ116028.1                                                              ||
## ||    KN149965.1                                                              ||
## ||    KN150349.1                                                              ||
## ||    KN149796.1                                                              ||
## ||    KN149849.1                                                              ||
## ||    KN150150.1                                                              ||
## ||    KN150203.1                                                              ||
## ||    KN150034.1                                                              ||
## ||    KN149703.1                                                              ||
## ||    KN150562.1                                                              ||
## ||    KN150615.1                                                              ||
## ||    KN150393.1                                                              ||
## ||    KN150446.1                                                              ||
## ||    KN149893.1                                                              ||
## ||    KZ116009.1                                                              ||
## ||    KN149946.1                                                              ||
## ||    KN150300.1                                                              ||
## ||    KN149777.1                                                              ||
## ||    KN150131.1                                                              ||
## ||    KN149800.1                                                              ||
## ||    KN150015.1                                                              ||
## ||    KN150689.1                                                              ||
## ||    KN150490.1                                                              ||
## ||    KN150543.1                                                              ||
## ||    KN150374.1                                                              ||
## ||    KZ116053.1                                                              ||
## ||    KN149990.1                                                              ||
## ||    KN149874.1                                                              ||
## ||    KN150258.1                                                              ||
## ||    KN149927.1                                                              ||
## ||    KN150089.1                                                              ||
## ||    KN149758.1                                                              ||
## ||    KN150112.1                                                              ||
## ||    KZ115960.1                                                              ||
## ||    KN150640.1                                                              ||
## ||    KN150471.1                                                              ||
## ||    KN150355.1                                                              ||
## ||    KN149971.1                                                              ||
## ||    KZ116034.1                                                              ||
## ||    KN150408.1                                                              ||
## ||    KN150186.1                                                              ||
## ||    KN149855.1                                                              ||
## ||    KN149908.1                                                              ||
## ||    KN149686.1                                                              ||
## ||    KN149739.1                                                              ||
## ||    KN150040.1                                                              ||
## ||    KN150598.1                                                              ||
## ||    KN150621.1                                                              ||
## ||    KN150452.1                                                              ||
## ||    KN150505.1                                                              ||
## ||    KN150283.1                                                              ||
## ||    KZ116015.1                                                              ||
## ||    KN150336.1                                                              ||
## ||    KN149783.1                                                              ||
## ||    KN150167.1                                                              ||
## ||    KN149836.1                                                              ||
## ||    KN150021.1                                                              ||
## ||    KN150579.1                                                              ||
## ||    KN150602.1                                                              ||
## ||    KN150380.1                                                              ||
## ||    KN150264.1                                                              ||
## ||    KN149880.1                                                              ||
## ||    KN149933.1                                                              ||
## ||    KN150317.1                                                              ||
## ||    KN150095.1                                                              ||
## ||    KN149764.1                                                              ||
## ||    KN150148.1                                                              ||
## ||    KN149817.1                                                              ||
## ||    KN150002.1                                                              ||
## ||    KZ115996.1                                                              ||
## ||    KN150676.1                                                              ||
## ||    KN150530.1                                                              ||
## ||    KZ116040.1                                                              ||
## ||    KN150361.1                                                              ||
## ||    KN150414.1                                                              ||
## ||    KN149861.1                                                              ||
## ||    KN149914.1                                                              ||
## ||    KN150076.1                                                              ||
## ||    KN149692.1                                                              ||
## ||    KN149745.1                                                              ||
## ||    KN147637.2                                                              ||
## ||    KZ115977.1                                                              ||
## ||    KN150488.1                                                              ||
## ||    KN150511.1                                                              ||
## ||    KN149988.1                                                              ||
## ||    KZ116021.1                                                              ||
## ||    KN150342.1                                                              ||
## ||    KN150173.1                                                              ||
## ||    KN150226.1                                                              ||
## ||    KN150057.1                                                              ||
## ||    KZ115958.1                                                              ||
## ||    KN150585.1                                                              ||
## ||    KN150469.1                                                              ||
## ||    KN149969.1                                                              ||
## ||    KN150270.1                                                              ||
## ||    KZ116002.1                                                              ||
## ||    KN150323.1                                                              ||
## ||    KN149770.1                                                              ||
## ||    KN150154.1                                                              ||
## ||    KN149823.1                                                              ||
## ||    KN150038.1                                                              ||
## ||    KN149707.2                                                              ||
## ||    KN150682.1                                                              ||
## ||    KN150619.1                                                              ||
## ||    KN150397.1                                                              ||
## ||    KN150420.1                                                              ||
## ||    KN149897.1                                                              ||
## ||    KN150251.1                                                              ||
## ||    KN150082.1                                                              ||
## ||    KN149751.1                                                              ||
## ||    KN149804.1                                                              ||
## ||    KN150019.2                                                              ||
## ||    KZ115983.1                                                              ||
## ||    KN150663.1                                                              ||
## ||    KN150494.1                                                              ||
## ||    KN150547.1                                                              ||
## ||    KZ116057.1                                                              ||
## ||    KN150401.1                                                              ||
## ||    KN149878.1                                                              ||
## ||    KN150232.1                                                              ||
## ||    KN149901.1                                                              ||
## ||    KN150063.1                                                              ||
## ||    KN149732.1                                                              ||
## ||    KZ115964.1                                                              ||
## ||    KN150591.1                                                              ||
## ||    KN150644.1                                                              ||
## ||    KN150475.1                                                              ||
## ||    KN150528.1                                                              ||
## ||    KZ116038.1                                                              ||
## ||    KN149859.1                                                              ||
## ||    KN150160.1                                                              ||
## ||    KN150213.1                                                              ||
## ||    KN150044.1                                                              ||
## ||    KN149713.1                                                              ||
## ||    KN147552.2                                                              ||
## ||    KN150572.1                                                              ||
## ||    KN150625.1                                                              ||
## ||    KN150509.1                                                              ||
## ||    KN150287.1                                                              ||
## ||    KN149956.1                                                              ||
## ||    KZ116019.1                                                              ||
## ||    KN150310.1                                                              ||
## ||    KN149787.1                                                              ||
## ||    KN150141.1                                                              ||
## ||    KN149810.1                                                              ||
## ||    KN150025.1                                                              ||
## ||    KN148038.2                                                              ||
## ||    KN150699.1                                                              ||
## ||    KN150553.1                                                              ||
## ||    KN150606.1                                                              ||
## ||    KZ116063.1                                                              ||
## ||    KN150384.1                                                              ||
## ||    KN150437.1                                                              ||
## ||    KN150268.1                                                              ||
## ||    KN149884.1                                                              ||
## ||    KN149937.1                                                              ||
## ||    KN149768.1                                                              ||
## ||    KN150122.1                                                              ||
## ||    KN150006.1                                                              ||
## ||    KZ115970.1                                                              ||
## ||    KN150650.1                                                              ||
## ||    KN150703.1                                                              ||
## ||    KN150481.1                                                              ||
## ||    KN150534.1                                                              ||
## ||    KN150365.1                                                              ||
## ||    KN149981.1                                                              ||
## ||    KZ116044.1                                                              ||
## ||    KN150418.1                                                              ||
## ||    KN150196.1                                                              ||
## ||    KN150249.1                                                              ||
## ||    KN149865.1                                                              ||
## ||    KN149696.2                                                              ||
## ||    KN149749.1                                                              ||
## ||    KN150103.1                                                              ||
## ||    KZ115951.1                                                              ||
## ||    KN150631.1                                                              ||
## ||    KN150462.1                                                              ||
## ||    KN150515.1                                                              ||
## ||    KN150293.1                                                              ||
## ||    KZ116025.1                                                              ||
## ||    KN149793.1                                                              ||
## ||    KN150177.1                                                              ||
## ||    KN149846.1                                                              ||
## ||    KN150200.1                                                              ||
## ||    KN150031.1                                                              ||
## ||    KN149700.1                                                              ||
## ||    KN150589.1                                                              ||
## ||    KN150390.1                                                              ||
## ||    KN150443.1                                                              ||
## ||    KN149890.1                                                              ||
## ||    KN150274.1                                                              ||
## ||    KZ116006.1                                                              ||
## ||    KN150327.1                                                              ||
## ||    KN149943.1                                                              ||
## ||    KN149774.1                                                              ||
## ||    KN150158.1                                                              ||
## ||    KN149827.1                                                              ||
## ||    KN150012.1                                                              ||
## ||    KN150540.1                                                              ||
## ||    KZ116050.1                                                              ||
## ||    KN150371.1                                                              ||
## ||    KN149871.1                                                              ||
## ||    KN150255.2                                                              ||
## ||    KN150308.1                                                              ||
## ||    KN149924.1                                                              ||
## ||    KN150086.1                                                              ||
## ||    KN149755.1                                                              ||
## ||    KZ115987.1                                                              ||
## ||    KN150667.1                                                              ||
## ||    KN150521.1                                                              ||
## ||    KN149998.1                                                              ||
## ||    KZ116031.1                                                              ||
## ||    KN150352.2                                                              ||
## ||    KN150405.1                                                              ||
## ||    KN150183.1                                                              ||
## ||    KN150236.1                                                              ||
## ||    KN149852.1                                                              ||
## ||    KN149905.1                                                              ||
## ||    KN150067.1                                                              ||
## ||    KN149683.1                                                              ||
## ||    KZ115968.1                                                              ||
## ||    KN150595.1                                                              ||
## ||    KN150479.1                                                              ||
## ||    KN150502.1                                                              ||
## ||    KN149979.1                                                              ||
## ||    KN150280.1                                                              ||
## ||    KZ116012.1                                                              ||
## ||    KN150333.1                                                              ||
## ||    KN150164.1                                                              ||
## ||    KN149780.1                                                              ||
## ||    KN150217.1                                                              ||
## ||    KN150048.1                                                              ||
## ||    KN149717.1                                                              ||
## ||    KZ115949.1                                                              ||
## ||    KN150576.1                                                              ||
## ||    KN150629.1                                                              ||
## ||    KN150430.1                                                              ||
## ||    KN150314.1                                                              ||
## ||    KN150092.1                                                              ||
## ||    KN150145.1                                                              ||
## ||    KN149761.1                                                              ||
## ||    KN149814.1                                                              ||
## ||    KN150029.1                                                              ||
## ||    KZ115993.1                                                              ||
## ||    KN150557.1                                                              ||
## ||    KN150388.1                                                              ||
## ||    KZ116067.1                                                              ||
## ||    KN150411.1                                                              ||
## ||    KN149888.1                                                              ||
## ||    KN150242.1                                                              ||
## ||    KN150073.1                                                              ||
## ||    KN149742.1                                                              ||
## ||    KN150126.1                                                              ||
## ||    KZ115974.1                                                              ||
## ||    KN150654.1                                                              ||
## ||    KN150485.1                                                              ||
## ||    KN150538.1                                                              ||
## ||    KZ116048.1                                                              ||
## ||    KN150369.1                                                              ||
## ||    KN150170.1                                                              ||
## ||    KN150054.1                                                              ||
## ||    KN149723.1                                                              ||
## ||    KZ115955.1                                                              ||
## ||    KN150582.1                                                              ||
## ||    KN150635.1                                                              ||
## ||    KN150466.1                                                              ||
## ||    KN150519.1                                                              ||
## ||    KN150297.1                                                              ||
## ||    KZ116029.1                                                              ||
## ||    KN150320.1                                                              ||
## ||    KN149797.1                                                              ||
## ||    KN150204.1                                                              ||
## ||    KN150035.1                                                              ||
## ||    KN150616.1                                                              ||
## ||    KN150394.1                                                              ||
## ||    KN150278.1                                                              ||
## ||    KN150301.1                                                              ||
## ||    KN149778.1                                                              ||
## ||    KN150132.1                                                              ||
## ||    KN149801.1                                                              ||
## ||    KN150016.1                                                              ||
## ||    KZ115980.1                                                              ||
## ||    KN150713.1                                                              ||
## ||    KN150491.1                                                              ||
## ||    KN150544.1                                                              ||
## ||    KN149991.1                                                              ||
## ||    KZ116054.1                                                              ||
## ||    KN150375.1                                                              ||
## ||    KN150428.1                                                              ||
## ||    KN150259.1                                                              ||
## ||    KN149928.1                                                              ||
## ||    KN150060.1                                                              ||
## ||    KN150113.1                                                              ||
## ||    KZ115961.1                                                              ||
## ||    KN150641.1                                                              ||
## ||    KN150472.1                                                              ||
## ||    KN150525.1                                                              ||
## ||    KZ116035.1                                                              ||
## ||    KN150187.1                                                              ||
## ||    KN149856.1                                                              ||
## ||    KN149909.1                                                              ||
## ||    KN150210.1                                                              ||
## ||    KN149687.1                                                              ||
## ||    KN150041.2                                                              ||
## ||    KN149710.1                                                              ||
## ||    KN150599.1                                                              ||
## ||    KN150622.1                                                              ||
## ||    KN150453.1                                                              ||
## ||    KN150506.1                                                              ||
## ||    KN150284.1                                                              ||
## ||    KZ116016.1                                                              ||
## ||    KN150337.1                                                              ||
## ||    KN149953.1                                                              ||
## ||    KN150168.1                                                              ||
## ||    KN149784.1                                                              ||
## ||    KN149837.1                                                              ||
## ||    KN150022.1                                                              ||
## ||    KN150550.1                                                              ||
## ||    KN150603.1                                                              ||
## ||    KZ116060.1                                                              ||
## ||    KN150434.1                                                              ||
## ||    KN149881.1                                                              ||
## ||    KN150265.1                                                              ||
## ||    KN150318.1                                                              ||
## ||    KN149765.1                                                              ||
## ||    KN149818.1                                                              ||
## ||    KZ115997.1                                                              ||
## ||    KN150677.1                                                              ||
## ||    KN150700.2                                                              ||
## ||    KZ116041.1                                                              ||
## ||    KN150362.1                                                              ||
## ||    KN150193.1                                                              ||
## ||    KN149862.1                                                              ||
## ||    KN150246.1                                                              ||
## ||    KN149915.1                                                              ||
## ||    KN149693.1                                                              ||
## ||    KN150077.1                                                              ||
## ||    KN149746.1                                                              ||
## ||    KN150100.1                                                              ||
## ||    KZ115978.1                                                              ||
## ||    KN150489.1                                                              ||
## ||    KN149989.1                                                              ||
## ||    KN150290.1                                                              ||
## ||    KZ116022.1                                                              ||
## ||    KN150343.1                                                              ||
## ||    KN150174.1                                                              ||
## ||    KN149790.1                                                              ||
## ||    KN150227.1                                                              ||
## ||    KN149843.1                                                              ||
## ||    KN150058.1                                                              ||
## ||    KN149727.1                                                              ||
## ||    KZ115959.1                                                              ||
## ||    KN150586.1                                                              ||
## ||    KN150639.1                                                              ||
## ||    KN150440.1                                                              ||
## ||    KN150271.1                                                              ||
## ||    KN149940.1                                                              ||
## ||    KZ116003.1                                                              ||
## ||    KN150324.1                                                              ||
## ||    KN150155.1                                                              ||
## ||    KN149771.1                                                              ||
## ||    KN149824.1                                                              ||
## ||    KN150039.1                                                              ||
## ||    KN149708.1                                                              ||
## ||    KN150683.1                                                              ||
## ||    KN150567.1                                                              ||
## ||    KN150398.1                                                              ||
## ||    KN150421.1                                                              ||
## ||    KN150252.1                                                              ||
## ||    KN149921.1                                                              ||
## ||    KN150083.1                                                              ||
## ||    KN150136.1                                                              ||
## ||    KN149805.1                                                              ||
## ||    KZ115984.1                                                              ||
## ||    KN150664.1                                                              ||
## ||    KN150495.1                                                              ||
## ||    KN150548.1                                                              ||
## ||    KZ116058.1                                                              ||
## ||    KN149995.1                                                              ||
## ||    KN150379.1                                                              ||
## ||    KN149879.1                                                              ||
## ||    KN150180.1                                                              ||
## ||    KN150233.1                                                              ||
## ||    KN150064.1                                                              ||
## ||    KN149680.1                                                              ||
## ||    KN149733.1                                                              ||
## ||    KN150117.1                                                              ||
## ||    KZ115965.1                                                              ||
## ||    KN150592.1                                                              ||
## ||    KN150645.1                                                              ||
## ||    KN150476.1                                                              ||
## ||    KN150529.1                                                              ||
## ||    KN149976.1                                                              ||
## ||    KZ116039.1                                                              ||
## ||    KN150330.1                                                              ||
## ||    KN150161.1                                                              ||
## ||    KN150214.1                                                              ||
## ||    KN150045.2                                                              ||
## ||    KN149714.1                                                              ||
## ||    KN150573.1                                                              ||
## ||    KN150626.1                                                              ||
## ||    KN149957.1                                                              ||
## ||    KN150311.1                                                              ||
## ||    KN149788.1                                                              ||
## ||    KN149811.1                                                              ||
## ||    KZ115990.1                                                              ||
## ||    KN150670.1                                                              ||
## ||    KN150554.1                                                              ||
## ||    KN150607.1                                                              ||
## ||    KN150385.1                                                              ||
## ||    KZ116064.1                                                              ||
## ||    KN149885.1                                                              ||
## ||    KN150269.1                                                              ||
## ||    KN149938.1                                                              ||
## ||    KN149769.1                                                              ||
## ||    KN150070.1                                                              ||
## ||    KN150123.1                                                              ||
## ||    KN150007.1                                                              ||
## ||    KZ115971.1                                                              ||
## ||    KN150651.1                                                              ||
## ||    KN150535.1                                                              ||
## ||    KZ116045.1                                                              ||
## ||    KN150366.1                                                              ||
## ||    KN149982.1                                                              ||
## ||    KN150419.1                                                              ||
## ||    KN150197.1                                                              ||
## ||    KN149866.1                                                              ||
## ||    KN149919.1                                                              ||
## ||    KN150220.1                                                              ||
## ||    KN149697.1                                                              ||
## ||    KN150051.1                                                              ||
## ||    KN149720.1                                                              ||
## ||    KN150104.1                                                              ||
## ||    KZ115952.1                                                              ||
## ||    KN150632.1                                                              ||
## ||    KN150516.1                                                              ||
## ||    KZ116026.1                                                              ||
## ||    KN150347.1                                                              ||
## ||    KN149794.1                                                              ||
## ||    KN149847.1                                                              ||
## ||    KN150201.1                                                              ||
## ||    KN150032.1                                                              ||
## ||    KN149701.1                                                              ||
## ||    KN150560.1                                                              ||
## ||    KN150613.1                                                              ||
## ||    KN150444.1                                                              ||
## ||    KN150275.1                                                              ||
## ||    KZ116007.1                                                              ||
## ||    KN150328.1                                                              ||
## ||    KN149944.1                                                              ||
## ||    KN150159.1                                                              ||
## ||    KN149775.2                                                              ||
## ||    KN149828.1                                                              ||
## ||    KN150013.1                                                              ||
## ||    KN150687.1                                                              ||
## ||    KN148869.2                                                              ||
## ||    KN150710.1                                                              ||
## ||    KN150541.1                                                              ||
## ||    KZ116051.1                                                              ||
## ||    KN150372.1                                                              ||
## ||    KN150425.1                                                              ||
## ||    KN149872.1                                                              ||
## ||    KN150309.1                                                              ||
## ||    KN149925.1                                                              ||
## ||    KN149756.1                                                              ||
## ||    KN149809.1                                                              ||
## ||    KN150110.1                                                              ||
## ||    KZ115988.1                                                              ||
## ||    KN150668.1                                                              ||
## ||    KN150499.1                                                              ||
## ||    KN150522.1                                                              ||
## ||    KN149999.1                                                              ||
## ||    KZ116032.1                                                              ||
## ||    KN150353.1                                                              ||
## ||    KN150406.1                                                              ||
## ||    KN150184.1                                                              ||
## ||    KN150237.1                                                              ||
## ||    KN149853.1                                                              ||
## ||    KN149906.1                                                              ||
## ||    KN150068.1                                                              ||
## ||    KN149684.1                                                              ||
## ||    KN149737.1                                                              ||
## ||    KZ115969.1                                                              ||
## ||    KN150596.1                                                              ||
## ||    KN150649.1                                                              ||
## ||    KN150450.1                                                              ||
## ||    KN150503.1                                                              ||
## ||    KN149950.1                                                              ||
## ||    KZ116013.1                                                              ||
## ||    KN150334.1                                                              ||
## ||    KN150165.1                                                              ||
## ||    KN150218.1                                                              ||
## ||    KN149834.1                                                              ||
## ||    KN150049.1                                                              ||
## ||    KN149718.1                                                              ||
## ||    KN150577.1                                                              ||
## ||    KN150431.1                                                              ||
## ||    KN149931.1                                                              ||
## ||    KN150315.1                                                              ||
## ||    KN150093.1                                                              ||
## ||    KN150146.1                                                              ||
## ||    KN149762.1                                                              ||
## ||    KN149815.1                                                              ||
## ||    KN150000.1                                                              ||
## ||    KZ115994.1                                                              ||
## ||    KN150558.1                                                              ||
## ||    KN150389.1                                                              ||
## ||    KN150412.1                                                              ||
## ||    KN149889.1                                                              ||
## ||    KN150190.1                                                              ||
## ||    KN150243.2                                                              ||
## ||    KN149912.1                                                              ||
## ||    KN150074.1                                                              ||
## ||    KN149690.1                                                              ||
## ||    KN149743.1                                                              ||
## ||    KN150127.1                                                              ||
## ||    KZ115975.1                                                              ||
## ||    KN150655.1                                                              ||
## ||    KN150708.1                                                              ||
## ||    KN150486.1                                                              ||
## ||    KZ116049.1                                                              ||
## ||    KN150340.1                                                              ||
## ||    KN150171.1                                                              ||
## ||    KN150224.1                                                              ||
## ||    KN149840.1                                                              ||
## ||    KN150055.1                                                              ||
## ||    KN149724.1                                                              ||
## ||    KZ115956.1                                                              ||
## ||    KN150583.1                                                              ||
## ||    KN150636.1                                                              ||
## ||    KN150467.1                                                              ||
## ||    KN149967.1                                                              ||
## ||    KZ116000.1                                                              ||
## ||    KN149798.1                                                              ||
## ||    KN150152.1                                                              ||
## ||    KN150205.1                                                              ||
## ||    KN149705.1                                                              ||
## ||    KN150680.1                                                              ||
## ||    KN150564.1                                                              ||
## ||    KN150617.1                                                              ||
## ||    KN150395.1                                                              ||
## ||    KN150279.1                                                              ||
## ||    KN149895.1                                                              ||
## ||    KN149948.1                                                              ||
## ||    KN150302.1                                                              ||
## ||    KN149779.1                                                              ||
## ||    KN150080.1                                                              ||
## ||    KN150133.2                                                              ||
## ||    KN150017.1                                                              ||
## ||    KZ115981.1                                                              ||
## ||    KN150661.1                                                              ||
## ||    KN150492.2                                                              ||
## ||    KN150545.1                                                              ||
## ||    KZ116055.1                                                              ||
## ||    KN149992.1                                                              ||
## ||    KN150429.1                                                              ||
## ||    KN149876.1                                                              ||
## ||    KN150230.1                                                              ||
## ||    KN150061.1                                                              ||
## ||    KN149730.1                                                              ||
## ||    KN150114.1                                                              ||
## ||    KZ115962.1                                                              ||
## ||    KN150642.1                                                              ||
## ||    KN150473.1                                                              ||
## ||    KN150526.1                                                              ||
## ||    KZ116036.1                                                              ||
## ||    KN150357.1                                                              ||
## ||    KN149973.1                                                              ||
## ||    KN149857.1                                                              ||
## ||    KN150211.1                                                              ||
## ||    KN149688.2                                                              ||
## ||    KN150042.1                                                              ||
## ||    KN149711.1                                                              ||
## ||    KN150570.1                                                              ||
## ||    KN150623.1                                                              ||
## ||    KN150454.1                                                              ||
## ||    KN150285.1                                                              ||
## ||    KN150338.1                                                              ||
## ||    KZ116017.1                                                              ||
## ||    KN149785.1                                                              ||
## ||    KN149838.1                                                              ||
## ||    KN150023.1                                                              ||
## ||    KN150697.1                                                              ||
## ||    KN150551.1                                                              ||
## ||    KZ116061.1                                                              ||
## ||    KN150382.1                                                              ||
## ||    KN149882.1                                                              ||
## ||    KN149935.1                                                              ||
## ||    KN150319.1                                                              ||
## ||    KN150097.1                                                              ||
## ||    KN149766.1                                                              ||
## ||    KN149819.1                                                              ||
## ||    KN150120.1                                                              ||
## ||    KN150004.2                                                              ||
## ||    KZ115998.1                                                              ||
## ||    KN150678.1                                                              ||
## ||    KN150701.1                                                              ||
## ||    KN150532.1                                                              ||
## ||    KZ116042.1                                                              ||
## ||    KN150363.1                                                              ||
## ||    KN150416.1                                                              ||
## ||    KN150194.1                                                              ||
## ||    KN149863.1                                                              ||
## ||    KN149916.1                                                              ||
## ||    KN150078.1                                                              ||
## ||    KN149694.1                                                              ||
## ||    KN149747.1                                                              ||
## ||    KZ115979.1                                                              ||
## ||    KN150659.1                                                              ||
## ||    KN150513.1                                                              ||
## ||    KN150291.1                                                              ||
## ||    KZ116023.1                                                              ||
## ||    KN149960.1                                                              ||
## ||    KN150344.1                                                              ||
## ||    KN149791.1                                                              ||
## ||    KN150175.1                                                              ||
## ||    KN149844.1                                                              ||
## ||    KN150228.1                                                              ||
## ||    KN150059.1                                                              ||
## ||    KN149728.1                                                              ||
## ||    KN150587.1                                                              ||
## ||    KN150610.1                                                              ||
## ||    KN150441.1                                                              ||
## ||    KN150272.1                                                              ||
## ||    KN149941.1                                                              ||
## ||    KZ116004.1                                                              ||
## ||    KN150325.1                                                              ||
## ||    KN149772.1                                                              ||
## ||    KN150156.1                                                              ||
## ||    KN149825.1                                                              ||
## ||    KN149709.1                                                              ||
## ||    KN150684.1                                                              ||
## ||    KN150568.1                                                              ||
## ||    KN150399.1                                                              ||
## ||    KN150422.1                                                              ||
## ||    KN149899.1                                                              ||
## ||    KN150253.1                                                              ||
## ||    KN149922.1                                                              ||
## ||    KN150306.1                                                              ||
## ||    KN150084.1                                                              ||
## ||    KN149753.1                                                              ||
## ||    KN150137.1                                                              ||
## ||    KN149806.1                                                              ||
## ||    KZ115985.1                                                              ||
## ||                                                                            ||
## || Global environment is initialised.                                         ||
## || Load the 1-th index block...                                               ||
## || The index block has been loaded.                                           ||
## || Start read mapping in chunk.                                               ||
## ||    0% completed, 0.1 mins elapsed, rate=99.6k reads per second             ||
## ||    6% completed, 0.3 mins elapsed, rate=112.5k reads per second            ||
## ||   13% completed, 0.5 mins elapsed, rate=112.3k reads per second            ||
## ||   20% completed, 0.9 mins elapsed, rate=90.6k reads per second             ||
## ||   21% completed, 0.9 mins elapsed, rate=92.0k reads per second             ||
## ||   22% completed, 1.0 mins elapsed, rate=93.2k reads per second             ||
## ||   23% completed, 1.0 mins elapsed, rate=94.5k reads per second             ||
## ||   25% completed, 1.0 mins elapsed, rate=95.8k reads per second             ||
## ||   26% completed, 1.0 mins elapsed, rate=96.9k reads per second             ||
## ||   27% completed, 1.1 mins elapsed, rate=98.0k reads per second             ||
## ||   28% completed, 1.1 mins elapsed, rate=99.0k reads per second             ||
## ||   29% completed, 1.1 mins elapsed, rate=99.9k reads per second             ||
## || Start read mapping in chunk.                                               ||
## ||   30% completed, 1.2 mins elapsed, rate=107.5k reads per second            ||
## ||   37% completed, 1.4 mins elapsed, rate=109.9k reads per second            ||
## ||   43% completed, 1.6 mins elapsed, rate=111.3k reads per second            ||
## ||   51% completed, 1.9 mins elapsed, rate=103.2k reads per second            ||
## ||   52% completed, 1.9 mins elapsed, rate=103.5k reads per second            ||
## ||   53% completed, 2.0 mins elapsed, rate=103.7k reads per second            ||
## ||   54% completed, 2.0 mins elapsed, rate=103.9k reads per second            ||
## ||   55% completed, 2.0 mins elapsed, rate=104.2k reads per second            ||
## ||   56% completed, 2.1 mins elapsed, rate=104.4k reads per second            ||
## ||   57% completed, 2.1 mins elapsed, rate=104.7k reads per second            ||
## ||   58% completed, 2.1 mins elapsed, rate=104.8k reads per second            ||
## ||   59% completed, 2.2 mins elapsed, rate=105.0k reads per second            ||
## ||   60% completed, 2.2 mins elapsed, rate=105.1k reads per second            ||
## || Start read mapping in chunk.                                               ||
## ||   60% completed, 2.2 mins elapsed, rate=109.1k reads per second            ||
## ||   67% completed, 2.4 mins elapsed, rate=110.1k reads per second            ||
## ||   73% completed, 2.6 mins elapsed, rate=111.1k reads per second            ||
## ||   81% completed, 3.0 mins elapsed, rate=105.8k reads per second            ||
## ||   82% completed, 3.0 mins elapsed, rate=105.9k reads per second            ||
## ||   83% completed, 3.1 mins elapsed, rate=105.8k reads per second            ||
## ||   84% completed, 3.1 mins elapsed, rate=105.7k reads per second            ||
## ||   85% completed, 3.1 mins elapsed, rate=105.7k reads per second            ||
## ||   86% completed, 3.2 mins elapsed, rate=105.6k reads per second            ||
## ||   87% completed, 3.2 mins elapsed, rate=105.6k reads per second            ||
## ||   88% completed, 3.2 mins elapsed, rate=105.5k reads per second            ||
## ||   89% completed, 3.3 mins elapsed, rate=105.5k reads per second            ||
## ||   90% completed, 3.3 mins elapsed, rate=105.5k reads per second            ||
## || Start read mapping in chunk.                                               ||
## ||   90% completed, 3.3 mins elapsed, rate=107.8k reads per second            ||
## ||   97% completed, 3.6 mins elapsed, rate=105.1k reads per second            ||
## ||   97% completed, 3.6 mins elapsed, rate=105.0k reads per second            ||
## ||   97% completed, 3.6 mins elapsed, rate=105.0k reads per second            ||
## ||   98% completed, 3.6 mins elapsed, rate=105.0k reads per second            ||
## ||   98% completed, 3.6 mins elapsed, rate=105.0k reads per second            ||
## ||   98% completed, 3.7 mins elapsed, rate=105.0k reads per second            ||
## ||   99% completed, 3.7 mins elapsed, rate=104.9k reads per second            ||
## ||   99% completed, 3.7 mins elapsed, rate=104.9k reads per second            ||
## ||   99% completed, 3.7 mins elapsed, rate=104.9k reads per second            ||
## ||                                                                            ||
## ||                           Completed successfully.                          ||
## ||                                                                            ||
## \\====================================    ====================================//
## 
## //================================   Summary =================================\\
## ||                                                                            ||
## ||                 Total reads : 23,279,297                                   ||
## ||                      Mapped : 15,235,962 (65.4%)                           ||
## ||             Uniquely mapped : 6,650,958                                    ||
## ||               Multi-mapping : 8,585,004                                    ||
## ||                                                                            ||
## ||                    Unmapped : 8,043,335                                    ||
## ||                                                                            ||
## ||                   Junctions : 85,589                                       ||
## ||                      Indels : 28,826                                       ||
## ||                                                                            ||
## ||                Running time : 3.7 minutes                                  ||
## ||                                                                            ||
## \\============================================================================//
```

```
rnacounts = featureCounts(files = list.files(pattern = "^9499X(4|12|20).*.sorted.bam$", 
                                             path = "Heart/bams/", full.names = TRUE),
                          minMQS = 30,
                          annot.ext = gtf_file,
                          isGTFAnnotationFile = TRUE,
                          nthreads = 8)
```

```
## 
##         ==========     _____ _    _ ____  _____  ______          _____  
##         =====         / ____| |  | |  _ \|  __ \|  ____|   /\   |  __ \ 
##           =====      | (___ | |  | | |_) | |__) | |__     /  \  | |  | |
##             ====      \___ \| |  | |  _ <|  _  /|  __|   / /\ \ | |  | |
##               ====    ____) | |__| | |_) | | \ \| |____ / ____ \| |__| |
##         ==========   |_____/ \____/|____/|_|  \_\______/_/    \_\_____/
##        Rsubread 2.0.1
## 
## //========================== featureCounts setting ===========================\\
## ||                                                                            ||
## ||             Input files : 2 BAM files                                      ||
## ||                           o 9499X12.sorted.bam                             ||
## ||                           o 9499X4.sorted.bam                              ||
## ||                                                                            ||
## ||              Annotation : Danio_rerio.GRCz11.93.chr.gtf (GTF)              ||
## ||      Dir for temp files : .                                                ||
## ||                 Threads : 8                                                ||
## ||                   Level : meta-feature level                               ||
## ||              Paired-end : no                                               ||
## ||      Multimapping reads : counted                                          ||
## || Multi-overlapping reads : not counted                                      ||
## ||   Min overlapping bases : 1                                                ||
## ||                                                                            ||
## \\============================================================================//
## 
## //================================= Running ==================================\\
## ||                                                                            ||
## || Load annotation file Danio_rerio.GRCz11.93.chr.gtf ...                     ||
## ||    Features : 481079                                                       ||
## ||    Meta-features : 32057                                                   ||
## ||    Chromosomes/contigs : 26                                                ||
## ||                                                                            ||
## || Process BAM file 9499X12.sorted.bam...                                     ||
## ||    Single-end reads are included.                                          ||
## ||    Total alignments : 26877266                                             ||
## ||    Successfully assigned alignments : 8750882 (32.6%)                      ||
## ||    Running time : 0.07 minutes                                             ||
## ||                                                                            ||
## || Process BAM file 9499X4.sorted.bam...                                      ||
## ||    Single-end reads are included.                                          ||
## ||    Total alignments : 23279297                                             ||
## ||    Successfully assigned alignments : 3468083 (14.9%)                      ||
## ||    Running time : 0.06 minutes                                             ||
## ||                                                                            ||
## || Write the final count table.                                               ||
## || Write the read assignment summary.                                         ||
## ||                                                                            ||
## \\============================================================================//
```

```
head(rnacounts$counts)
```

```
##                    9499X12.sorted.bam 9499X4.sorted.bam
## ENSDARG00000102141                317               152
## ENSDARG00000102123                  8                 0
## ENSDARG00000114503                 76                18
## ENSDARG00000115971                 13                 3
## ENSDARG00000098311                139                11
## ENSDARG00000104839                346                56
```

```
total_rnacounts <- apply(rnacounts$counts, 1, min)
head(total_rnacounts)
```

```
## ENSDARG00000102141 ENSDARG00000102123 ENSDARG00000114503 ENSDARG00000115971 
##                152                  0                 18                  3 
## ENSDARG00000098311 ENSDARG00000104839 
##                 11                 56
```

```
genecounts <- data.frame(genecounts)
genecounts$rnacounts = total_rnacounts[row.names(genecounts)]
colnames(genecounts) <- c("sgRNAs", "rna")
inboth <- sum(genecounts$rna > 0 & genecounts$sgRNAs > 0)
incrispr <-  sum(genecounts$rna == 0 & genecounts$sgRNAs > 0)
inheart <- sum(genecounts$rna > 0 & genecounts$sgRNAs == 0)


cor(genecounts$sgRNAs, genecounts$rna, method = "pearson")
```

```
## [1] 0.09232218
```

```
plot(log2(genecounts$sgRNAs) ~ log2(genecounts$rna))
```

```
qplot(log2(genecounts$rna), log2(genecounts$sgRNAs))
```

```
ggplot(genecounts[genecounts$rna > 0 & genecounts$sgRNAs > 0, ], 
       aes(x = log2(rna), y = log2(sgRNAs))) + 
  geom_point(alpha = .3, color = "royalblue4") +
  theme(text = element_text(size = 24))
```

#### Session Information

```
sessionInfo()
```

```
## R version 3.6.3 (2020-02-29)
## Platform: x86_64-pc-linux-gnu (64-bit)
## Running under: Debian GNU/Linux 9 (stretch)
## 
## Matrix products: default
## BLAS:   /usr/lib/openblas-base/libblas.so.3
## LAPACK: /usr/lib/libopenblasp-r0.2.19.so
## 
## locale:
##  [1] LC_CTYPE=en_US.UTF-8       LC_NUMERIC=en_US.UTF-8    
##  [3] LC_TIME=en_US.UTF-8        LC_COLLATE=en_US.UTF-8    
##  [5] LC_MONETARY=en_US.UTF-8    LC_MESSAGES=en_US.UTF-8   
##  [7] LC_PAPER=en_US.UTF-8       LC_NAME=C                 
##  [9] LC_ADDRESS=C               LC_TELEPHONE=C            
## [11] LC_MEASUREMENT=en_US.UTF-8 LC_IDENTIFICATION=C       
## 
## attached base packages:
##  [1] grid      stats4    parallel  stats     graphics  grDevices utils    
##  [8] datasets  methods   base     
## 
## other attached packages:
##  [1] ggplot2_3.3.0        VennDiagram_1.6.20   futile.logger_1.4.3 
##  [4] Rsamtools_2.2.3      Biostrings_2.54.0    XVector_0.26.0      
##  [7] GenomicRanges_1.38.0 GenomeInfoDb_1.22.1  IRanges_2.20.2      
## [10] S4Vectors_0.24.4     BiocGenerics_0.32.0  Rsubread_2.0.1      
## 
## loaded via a namespace (and not attached):
##  [1] Rcpp_1.0.4.6           pillar_1.4.3           compiler_3.6.3        
##  [4] formatR_1.7            bitops_1.0-6           futile.options_1.0.1  
##  [7] tools_3.6.3            zlibbioc_1.32.0        digest_0.6.25         
## [10] jsonlite_1.6.1         evaluate_0.14          lifecycle_0.2.0       
## [13] tibble_3.0.1           gtable_0.3.0           pkgconfig_2.0.3       
## [16] rlang_0.4.5            yaml_2.2.1             xfun_0.13             
## [19] GenomeInfoDbData_1.2.2 withr_2.2.0            dplyr_0.8.5           
## [22] stringr_1.4.0          knitr_1.28             vctrs_0.2.4           
## [25] tidyselect_1.0.0       glue_1.4.0             R6_2.4.1              
## [28] BiocParallel_1.20.1    rmarkdown_2.1          farver_2.0.3          
## [31] purrr_0.3.4            lambda.r_1.2.4         magrittr_1.5          
## [34] ellipsis_0.3.0         scales_1.1.0           htmltools_0.4.0       
## [37] assertthat_0.2.1       colorspace_1.4-1       labeling_0.3          
## [40] stringi_1.4.6          RCurl_1.98-1.2         munsell_0.5.0         
## [43] crayon_1.3.4
```
