## Supplemental Note 4 for "SLALOM: A Simple and Rapid Method for Enzymatic Synthesis of CRISPR-Cas9 sgRNA Libraries": SupplementalNote4.html

CrispyR analysis of GFP Screen: First 3 Sites


### CrispyR analysis of GFP Screen: First 3 Sites

###### Jonathon Hill

Cris-py is designed to analyze indels from CRISPR mutagenesis. However, it is struggling with my files and saying none have any valid sequences, but I can find them when I look manually. I will re-create the code here to do my own analysis.

Cris-py steps: 1. Identify sequences and reverse compliment if necessary. 2. Measure lengths to get overall Indel percentage. 3. Look for deletion of specific guides to attribute Indels.

One issue is that Cris-py is not designed for close sites where deletions may cover more than one sgRNA site. I will address that here.

#### Inputs

```
start_seq <- toupper("cgcaaatgggcggtaggcgtgt")
end_seq <- toupper("tcgtgaccaccctgacctacggc")
sites <- toupper(c(site1 = "cgaggagctgttcaccggggtggtgcccat", 
                   site2 = "caagttcagcgtgtccggcgagggcgaggg",
                   site3 = "gttcatctgcaccaccggcaagctgcccgt")) 
wt_seq <- toupper(gsub("[ \t\r\n]", "", 
                       "CGCAAATGGGCGGTAGGCGTGTACGGTGGGAGGTCTATATAAGCAGAGCT
                        CTCTGGCTAACTAGAGAACCCACTGCTTACTGGCTTATCGAAATTAATAC
                        GACTCACTATAGGGAGACCCAAGCTTGGTACCGAGCTCGGATCGATATCT
                        GCGGCCTAGCTAGCGCTACCGGTCGCCACCATGGTGAGCAAGGGCGAGGA
                        GCTGTTCACCGGGGTGGTGCCCATCCTGGTCGAGCTGGACGGCGACGTAA
                        GCGGCCACAAGTTCAGCGTGTCCGGCGAGGGCGAGGGCGATGCCACCTAC
                        TGCAAGCTGACCCTGAAGTTCATCTGCACCACCGGCAAGCTGCCCGTGCC
                        CTGGCCCACCCTCGTGACCACCCTGACCTACGGC"))
pcr_size <- nchar(wt_seq)
con_fastq <- "Con3.extendedFrags.fastq"
pos_fastq <- "Pos3.extendedFrags.fastq"
neg_fastq <- "Neg3.extendedFrags.fastq"
```

#### Functions

```
library(data.table, quietly = TRUE)
library(msa)
```

```
## Loading required package: Biostrings
```

```
## Loading required package: BiocGenerics
```

```
## Loading required package: parallel
```

```
## 
## Attaching package: 'BiocGenerics'
```

```
## The following objects are masked from 'package:parallel':
## 
##     clusterApply, clusterApplyLB, clusterCall, clusterEvalQ,
##     clusterExport, clusterMap, parApply, parCapply, parLapply,
##     parLapplyLB, parRapply, parSapply, parSapplyLB
```

```
## The following objects are masked from 'package:stats':
## 
##     IQR, mad, sd, var, xtabs
```

```
## The following objects are masked from 'package:base':
## 
##     anyDuplicated, append, as.data.frame, basename, cbind, colnames,
##     dirname, do.call, duplicated, eval, evalq, Filter, Find, get, grep,
##     grepl, intersect, is.unsorted, lapply, Map, mapply, match, mget,
##     order, paste, pmax, pmax.int, pmin, pmin.int, Position, rank,
##     rbind, Reduce, rownames, sapply, setdiff, sort, table, tapply,
##     union, unique, unsplit, which, which.max, which.min
```

```
## Loading required package: S4Vectors
```

```
## Loading required package: stats4
```

```
## 
## Attaching package: 'S4Vectors'
```

```
## The following objects are masked from 'package:data.table':
## 
##     first, second
```

```
## The following object is masked from 'package:base':
## 
##     expand.grid
```

```
## Loading required package: IRanges
```

```
## 
## Attaching package: 'IRanges'
```

```
## The following object is masked from 'package:data.table':
## 
##     shift
```

```
## Loading required package: XVector
```

```
## 
## Attaching package: 'Biostrings'
```

```
## The following object is masked from 'package:base':
## 
##     strsplit
```

```
library(Biostrings)

read_fastq <- function(fastq_name) {
  # Read fastq
  fastq <- fread(fastq_name, 
                 header = FALSE, 
                 sep = NULL, 
                 col.names = "Values", 
                 stringsAsFactors = FALSE)
  
  # Fastq has four lines per record, so need to process
  fastq[, "Type" := rep(c("Name", "Seq", "Plus", "Qual"), 
                        length.out = nrow(fastq))] 
  fastq[, "ID" := rep(1:(nrow(fastq)/4), each = 4)]
  fastq <- dcast(fastq, ID ~ Type, value.var = "Values")
  
  return(fastq)
}

process_seqs <- function(seqs, start_seq, end_seq) {
  # Set up search for complete PCR products
  re <- paste0(start_seq, ".*", end_seq)
  
  # Convert to DNAStringSet for faster processing
  seqs <- DNAStringSet(seqs)
  
  # Get complete PCR products (rev comp when needed)
  forward_seqs <- seqs[grepl(re, seqs)]
  reverse_seqs <- reverseComplement(seqs)
  reverse_seqs <- reverse_seqs[grepl(re, reverse_seqs)]
  total_valid <- length(forward_seqs) + length(reverse_seqs)
  cat(paste("Number Valid/Total:", total_valid, "/", length(seqs)), sep = "\n")
  
  return(c(forward_seqs, reverse_seqs))
}

find_muts <- function(site, seqs, pcr_size) {
  # Find mutations in target site
  muts <- !grepl(site, seqs)
  
  # find which of those is an indel
  indels <- width(seqs) != pcr_size & muts
  
  return(list(muts = muts, indels = indels))
}

get_mut_df <- function(seqs, sites, pcr_size) {
  # Find sites affected in each sequence
  mut_list <- lapply(sites, find_muts, seqs, pcr_size)
  mut_df <- data.frame(mut_list)
  mut_df$total_sites_mut <- rowSums(data.frame(mut_df[, grepl("muts", 
                                                   colnames(mut_df))]))
  mut_df$total_sites_indel <- rowSums(data.frame(mut_df[, grepl("indels", 
                                                     colnames(mut_df))]))
  mut_df <- cbind(seqs, mut_df)
  return(mut_df)
}

print_stats <- function(mut_df, mut_sites, 
                        min_sites = 1, max_sites = 1, common = FALSE, n = 10) {
  re <- paste0(mut_sites, collapse = "|")
  
  # Mutants
  re_mut <- paste0("[", re, "].muts")
  mutated <- rowSums(data.frame(mut_df[, grepl(re_mut, colnames(mut_df))]))
  valid_mut <- mutated >= min_sites & mut_df$total_sites_mut <= max_sites
  mut_rate <- paste0("Mutation Rate: ", sum(valid_mut), " / ", nrow(mut_df), 
                     " (", sum(valid_mut)/nrow(mut_df)*100, "%)\n")
  
  # Indels
  re_indel <- paste0("[", re, "].indels")
  indel <- rowSums(data.frame(mut_df[, grepl(re_indel, colnames(mut_df))]))
  valid_indel <- indel >= min_sites & mut_df$total_sites_indel <= max_sites
  indel_rate <- paste0("Indel Rate: ", sum(valid_indel), " / ", nrow(mut_df), 
                       " (", sum(valid_indel)/nrow(mut_df)*100, "%)\n")
  
  results <- c(mut_rate, indel_rate)
  
  # Common alleles
  if (common) {
    common_muts <- table(mut_df$seqs[valid_mut])
    common_muts <- common_muts[order(common_muts, decreasing = TRUE)]
    common_muts <- data.frame(common_muts)
    alignments <- lapply(head(as.character(common_muts$Var1), n = n), 
                         pairwiseAlignment, 
                         subject = wt_seq, 
                         gapOpening = 10, 
                         gapExtension = 0)
    alignments <- lapply(alignments, function(x) toString(alignedPattern(x)))
    results <- c(results, paste0(unlist(alignments), collapse = "\n"))
  }
  return(results)
}
```

#### Control

```
seqs <- read_fastq(con_fastq)$Seq
filtered_seqs <- process_seqs(seqs, start_seq, end_seq)
```

```
## Number Valid/Total: 27201 / 34065
```

```
mut_df <- get_mut_df(filtered_seqs, sites, pcr_size)
cat("Site 1:\n",
print_stats(mut_df, mut_sites = "site1"))
```

```
## Site 1:
##  Mutation Rate: 260 / 27201 (0.955847211499577%)
##  Indel Rate: 8 / 27201 (0.0294106834307562%)
```

```
cat("Site 2:\n",
print_stats(mut_df, mut_sites = "site2"))
```

```
## Site 2:
##  Mutation Rate: 331 / 27201 (1.21686702694754%)
##  Indel Rate: 4 / 27201 (0.0147053417153781%)
```

```
cat("Site 3:\n",
print_stats(mut_df, mut_sites = "site3"))
```

```
## Site 3:
##  Mutation Rate: 371 / 27201 (1.36392044410132%)
##  Indel Rate: 8 / 27201 (0.0294106834307562%)
```

```
cat("Multiple Sites:\n",
print_stats(mut_df, mut_sites = c("site1", "site2", "site3"), 
            min_sites = 2, max_sites = 3))
```

```
## Multiple Sites:
##  Mutation Rate: 79 / 27201 (0.290430498878718%)
##  Indel Rate: 4 / 27201 (0.0147053417153781%)
```

#### GFP Positive Pool

```
seqs <- read_fastq(pos_fastq)$Seq
filtered_seqs <- process_seqs(seqs, start_seq, end_seq)
```

```
## Number Valid/Total: 33468 / 40512
```

```
mut_df <- get_mut_df(filtered_seqs, sites, pcr_size)
cat("Site 1:\n",
print_stats(mut_df, mut_sites = "site1"))
```

```
## Site 1:
##  Mutation Rate: 379 / 33468 (1.13242500298793%)
##  Indel Rate: 83 / 33468 (0.247998087725589%)
```

```
cat("Site 2:\n",
print_stats(mut_df, mut_sites = "site2"))
```

```
## Site 2:
##  Mutation Rate: 675 / 33468 (2.01685191825027%)
##  Indel Rate: 358 / 33468 (1.06967849886459%)
```

```
cat("Site 3:\n",
print_stats(mut_df, mut_sites = "site3"))
```

```
## Site 3:
##  Mutation Rate: 430 / 33468 (1.28480937014462%)
##  Indel Rate: 70 / 33468 (0.209155013744472%)
```

```
cat("Multiple Sites:\n",
print_stats(mut_df, mut_sites = c("site1", "site2", "site3"), 
            min_sites = 2, max_sites = 3))
```

```
## Multiple Sites:
##  Mutation Rate: 75 / 33468 (0.224094657583363%)
##  Indel Rate: 10 / 33468 (0.0298792876777818%)
```

#### GFP Negative Pool

```
seqs <- read_fastq(neg_fastq)$Seq
filtered_seqs <- process_seqs(seqs, start_seq, end_seq)
```

```
## Number Valid/Total: 41187 / 51220
```

```
mut_df <- get_mut_df(filtered_seqs, sites, pcr_size)
cat("Site 1:\n",
print_stats(mut_df, mut_sites = "site1", common = TRUE))
```

```
## Site 1:
##  Mutation Rate: 3383 / 41187 (8.2137567679122%)
##  Indel Rate: 3259 / 41187 (7.91269089761332%)
##  CGCAAATGGGCGGTAGGCGTGTACGGTGGGAGGTCTATATAAGCAGAGCTCTCTGGCTAACTAGAGAACCCACTGCTTACTGGCTTATCGAAATTAATACGACTCACTATAGGGAGACCCAAGCTTGGTACCGAGCTCGGATCGATATCTGCGGCCTAGCTAGCGCTACCGGTCGCCACCATGGTGAGCAAGGGCGAGGAGCTGTTCACCGGGG-GGTGCCCATCCTGGTCGAGCTGGACGGCGACGTAAACGGCCACAAGTTCAGCGTGTCCGGCGAGGGCGAGGGCGATGCCACCTACGGCAAGCTGACCCTGAAGTTCATCTGCACCACCGGCAAGCTGCCCGTGCCCTGGCCCACCCTCGTGACCACCCTGACCTACGGC
## CGCAAATGGGCGGTAGGCGTGTACGGTGGGAGGTCTATATAAGCAGAGCTCTCTGGCTAACTAGAGAACCCACTGCTTACTGGCTTATCGAAATTAATACGACTCACTATAGGGAGACCCAAGCTTGGTACCGAGCTCGGATCGATATCTGCGGCCTAGCTAGCGCTACCGGTCGCCACCATGGTGAGCAAGGGCGAGGAGC------------------CCATCCTGGTCGAGCTGGACGGCGACGTAAACGGCCACAAGTTCAGCGTGTCCGGCGAGGGCGAGGGCGATGCCACCTACGGCAAGCTGACCCTGAAGTTCATCTGCACCACCGGCAAGCTGCCCGTGCCCTGGCCCACCCTCGTGACCACCCTGACCTACGGC
## CGCAAATGGGCGGTAGGCGTGTACGGTGGGAGGTCTATATAAGCAGAGCTCTCTGGCTAACTAGAGAACCCACTGCTTACTGGCTTATCGAAATTAATACGACTCACTATAGGGAGACCCAAGCTTGGTACCGAGCTCGGATCGATATCTGCGGCCTAGCTAGCGCTACCGGTCGCCACCATGGTGAGCAAGGGCG---------------GGGTGGTGCCCATCCTGGTCGAGCTGGACGGCGACGTAAACGGCCACAAGTTCAGCGTGTCCGGCGAGGGCGAGGGCGATGCCACCTACGGCAAGCTGACCCTGAAGTTCATCTGCACCACCGGCAAGCTGCCCGTGCCCTGGCCCACCCTCGTGACCACCCTGACCTACGGC
## CGCAAATGGGCGGTAGGCGTGTACGGTGGGAGGTCTATATAAGCAGAGCTCTCTGGCTAACTAGAGAACCCACTGCTTACTGGCTTATCGAAATTAATACGACTCACTATAGGGAGACCCAAGCTTGGTACCGAGCTCGGATCGATATCTGCGGCCTAGCTAGCGCTACCGGTCGCCACCAT------------------------------------------CCTGGTCGAGCTGGACGGCGACGTAAACGGCCACAAGTTCAGCGTGTCCGGCGAGGGCGAGGGCGATGCCACCTACGGCAAGCTGACCCTGAAGTTCATCTGCACCACCGGCAAGCTGCCCGTGCCCTGGCCCACCCTCGTGACCACCCTGACCTACGGC
## CGCAAATGGGCGGTAGGCGTGTACGGTGGGAGGTCTATATAAGCAGAGCTCTCTGGCTAACTAGAGAACCCACTGCTTACTGGCTTATCGAAATTAATACGACTCACTATAGGGAGACCCAAGCTTGGTACCGAGCTCGGATCGATATCTGCGGCCTAGCTAGCGCTACCGGTCGCCACCATGGTGAGCAAGGGC------------ACCGGGGTGGTGCCCATCCTGGTCGAGCTGGACGGCGACGTAAACGGCCACAAGTTCAGCGTGTCCGGCGAGGGCGAGGGCGATGCCACCTACGGCAAGCTGACCCTGAAGTTCATCTGCACCACCGGCAAGCTGCCCGTGCCCTGGCCCACCCTCGTGACCACCCTGACCTACGGC
## CGCAAATGGGCGGTAGGCGTGTACGGTGGGAGGTCTATATAAGCAGAGCTCTCTGGCTAACTAGAGAACCCACTGCTTACTGGCTTATCGAAATTAATACGACTCACTATAGGGAGACCCAAGCTTGGTACCGAGCTCGGATCGATATCTGCGGCCTAGCTAGCGCTACCGGTCGCCACCATGGTGAGCAAGGG-------------------GTGGTGCCCATCCTGGTCGAGCTGGACGGCGACGTAAACGGCCACAAGTTCAGCGTGTCCGGCGAGGGCGAGGGCGATGCCACCTACGGCAAGCTGACCCTGAAGTTCATCTGCACCACCGGCAAGCTGCCCGTGCCCTGGCCCACCCTCGTGACCACCCTGACCTACGGC
## CGCAAATGGGCGGTAGGCGTGTACGGTGGGAGGTCTATATAAGCAGAGCTCTCTGGCTAACTAGAGAACCCACTGCTTACTGGCTTATCGAAATTAATACGACTCACTATAGGGAGACCCAAGCTTGGTACCGAGCTCGGATCGATATCTGCGGCCTAGCTAGCGCTACCGGTCGCCACCATGGTGAGCAAGGGCGAGGAGCTGT----------GGTGCCCATCCTGGTCGAGCTGGACGGCGACGTAAACGGCCACAAGTTCAGCGTGTCCGGCGAGGGCGAGGGCGATGCCACCTACGGCAAGCTGACCCTGAAGTTCATCTGCACCACCGGCAAGCTGCCCGTGCCCTGGCCCACCCTCGTGACCACCCTGACCTACGGC
## CGCAAATGGGCGGTAGGCGTGTACGGTGGGAGGTCTATATAAGCAGAGCTCTCTGGCTAACTAGAGAACCCACTGC----------------------------------------------------------------------------------------------------------------------------------------GGTGGTGCCCATCCTGGTCGAGCTGGACGGCGACGTAAACGGCCACAAGTTCAGCGTGTCCGGCGAGGGCGAGGGCGATGCCACCTACGGCAAGCTGACCCTGAAGTTCATCTGCACCACCGGCAAGCTGCCCGTGCCCTGGCCCACCCTCGTGACCACCCTGACCTACGGC
## CGCAAATGGGCGGTAGGCGTGTACGGTGGGAGGTCTATATAAGCAGAGCTCTCTGGCTAACTAGAGAACCCACTGCTTACTGGCTTATCGAAATTAATACGACTCACTATAGGGAGACCCAAGCTTGGTACCGAGCTCGGATCGATATCTGCGGCCTAGCTAGCGCTACCGGTCGCCACCATGGTGAGCAAGGGCGAGGAGCTG---------------------------------GACGGCGACGTAAACGGCCACAAGTTCAGCGTGTCCGGCGAGGGCGAGGGCGATGCCACCTACGGCAAGCTGACCCTGAAGTTCATCTGCACCACCGGCAAGCTGCCCGTGCCCTGGCCCACCCTCGTGACCACCCTGACCTACGGC
## CGCAAATGGGCGGTAGGCGTGTACGGTGGGAGGTCTATATAAGCAGAGCTCTCTGGCTAACTAGAGAACCCACTGCTTACTGGCTTATCGAAATTAATACGACTCACTATAGGGAGACCCAAGCTTGGTACCGAGCTCGGATCGATATCTGCGGCCTAGCTAGCGCTACCGGTCGCCACCATGGTGAGCAAGGGCGAGGAGCTGTTCACCGGGGTG---CCCATCCTGGTCGAGCTGGACGGCGACGTAAACGGCCACAAGTTCAGCGTGTCCGGCGAGGGCGAGGGCGATGCCACCTACGGCAAGCTGACCCTGAAGTTCATCTGCACCACCGGCAAGCTGCCCGTGCCCTGGCCCACCCTCGTGACCACCCTGACCTACGGC
```

```
cat("Site 2:\n",
print_stats(mut_df, mut_sites = "site2", common = TRUE))
```

```
## Site 2:
##  Mutation Rate: 21796 / 41187 (52.9196105567291%)
##  Indel Rate: 21606 / 41187 (52.458299949013%)
##  CGCAAATGGGCGGTAGGCGTGTACGGTGGGAGGTCTATATAAGCAGAGCTCTCTGGCTAACTAGAGAACCCACTGCTTACTGGCTTATCGAAATTAATACGACTCACTATAGGGAGACCCAAGCTTGGTACCGAGCTCGGATCGATATCTGCGGCCTAGCTAGCGCTACCGGTCGCCACCATGGTGAGCAAGGGCGAGGAGCTGTTCACCGGGGTGGTGCCCATCCTGGTCGAGCTGGACGGCGACGTAAACGGCCACAAGTTCAGCGTGTCCGGCGAGGGCGA------TGCCACCTACGGCAAGCTGACCCTGAAGTTCATCTGCACCACCGGCAAGCTGCCCGTGCCCTGGCCCACCCTCGTGACCACCCTGACCTACGGC
## CGCAAATGGGCGGTAGGCGTGTACGGTGGGAGGTCTATATAAGCAGAGCTCTCTGGCTAACTAGAGAACCCACTGCTTACTGGCTTATCGAAATTAATACGACTCACTATAGGGAGACCCAAGCTTGGTACCGAGCTCGGATCGATATCTGCGGCCTAGCTAGCGCTACCGGTCGCCACCATGGTGAGCAAGGGCGAGGAGCTGTTCACCGGGGTGGTGCCCATCCTGGTCGAGCTGGACGGCGACGTAAACGGCCACAAGTTCAGCGTGTCCGGCGA------------TGCCACCTACGGCAAGCTGACCCTGAAGTTCATCTGCACCACCGGCAAGCTGCCCGTGCCCTGGCCCACCCTCGTGACCACCCTGACCTACGGC
## CGCAAATGGGCGGTAGGCGTGTACGGTGGGAGGTCTATATAAGCAGAGCTCTCTGGCTAACTAGAGAACCCACTGCTTACTGGCTTATCGAAATTAATACGACTCACTATAGGGAGACCCAAGCTTGGTACCGAGCTCGGATCGATATCTGCGGCCTAGCTAGCGCTACCGGTCGCCACCATGGTGAGCAAGGGCGAGGAGCTGTTCACCGGGGTGGTGCCCATCCTGGTCGAGCTGGACGGCGACGTAAACGGCCACAAGTTCAGCGTGTCCGGCGAAGGGCGAGGGCGATGCCACCTACGGCAAGCTGACCCTGAAGTTCATCTGCACCACCGGCAAGCTGCCCGTGCCCTGGCCCACCCTCGTGACCACCCTGACCTACGGC
## CGCAAATGGGCGGTAGGCGTGTACGGTGGGAGGTCTATATAAGCAGAGCTCTCTGGCTAACTAGAGAACCCACTGCTTACTGGCTTATCGAAATTAATACGACTCACTATAGGGAGACCCAAGCTTGGTACCGAGCTCGGATCGATATCTGCGGCCTAGCTAGCGCTACCGGTCGCCACCATGGTGAGCAAGGGCGAGGAGCTGTTCACCGGGGTGGTGCCCATCCTGGTCGAGCTGGACGGCGACGTAAACGGCCACAAGTTCAGCG---------------AGGGCGATGCCACCTACGGCAAGCTGACCCTGAAGTTCATCTGCACCACCGGCAAGCTGCCCGTGCCCTGGCCCACCCTCGTGACCACCCTGACCTACGGC
## CGCAAATGGGCGGTAGGCGTGTACGGTGGGAGGTCTATATAAGCAGAGCTCTCTGGCTAACTAGAGAACCCACTGCTTACTGGCTTATCGAAATTAATACGACTCACTATAGGGAGACCCAAGCTTGGTACCGAGCTCGGATCGATATCTGCGGCCTAGCTAGCGCTACCGGTCGCCACCATGGTGAGCAAGGGCGAGGAGCTGTTCACCGGGGTGGTGCCCATCCTGGTCGAGCTGGACGGCGACGTAAACGGCCACAAGTTCAG-------------GGCGAGGGCGATGCCACCTACGGCAAGCTGACCCTGAAGTTCATCTGCACCACCGGCAAGCTGCCCGTGCCCTGGCCCACCCTCGTGACCACCCTGACCTACGGC
## CGCAAATGGGCGGTAGGCGTGTACGGTGGGAGGTCTATATAAGCAGAGCTCTCTGGCTAACTAGAGAACCCACTGCTTACTGGCTTATCGAAATTAATACGACTCACTATAGGGAGACCCAAGCTTGGTACCGAGCTCGGATCGATATCTGCGGCCTAGCTAGCGCTACCGGTCGCCACCATGGTGAGCAAGGGCGAGGAGCTGTTCACCGGGGTGGTGCCCATCCTGGTCGAGCTGGACGGCGACGTAAACGGCCACAAG------------------GGCGAGGGCGATGCCACCTACGGCAAGCTGACCCTGAAGTTCATCTGCACCACCGGCAAGCTGCCCGTGCCCTGGCCCACCCTCGTGACCACCCTGACCTACGGC
## CGCAAATGGGCGGTAGGCGTGTACGGTGGGAGGTCTATATAAGCAGAGCTCTCTGGCTAACTAGAGAACCCACTGCTTACTGGCTTATCGAAATTAATACGACTCACTATAGGGAGACCCAAGCTTGGTACCGAGCTCGGATCGATATCTGCGGCCTAGCTAGCGCTACCGGTCGCCACCATGGTGAGCAAGGGCGAGGAGCTGTTCACCGGGGTGGTGCCCATCCTGGTCGAGCTGGACGGCGACGTAAACGGCCACAAGTTCAGCGTGTCCGGC-AGGGCGAGGGCGATGCCACCTACGGCAAGCTGACCCTGAAGTTCATCTGCACCACCGGCAAGCTGCCCGTGCCCTGGCCCACCCTCGTGACCACCCTGACCTACGGC
## CGCAAATGGGCGGTAGGCGTGTACGGTGGGAGGTCTATATAAGCAGAGCTCTCTGGCTAACTAGAGAACCCACTGCTTACTGGCTTATCGAAATTAATACGACTCACTATAGGGAGACCCAAGCTTGGTACCGAGCTCGGATCGATATCTGCGGCCTAGCTAGCGCTACCGGTCGCCACCATGGTGAGCAAGGGCGAGGAGCTGTTCACCGGGGTGGTGCCCATCCTGGTCGAGCTGGACGGCGACGTAAACGGCCACAAGTTCAGCGTGTCCGGCG-GGGCGAGGGCGATGCCACCTACGGCAAGCTGACCCTGAAGTTCATCTGCACCACCGGCAAGCTGCCCGTGCCCTGGCCCACCCTCGTGACCACCCTGACCTACGGC
## CGCAAATGGGCGGTAGGCGTGTACGGTGGGAGGTCTATATAAGCAGAGCTCTCTGGCTAACTAGAGAACCCACTGCTTACTGGCTTATCGAAATTAATACGACTCACTATAGGGAGACCCAAGCTTGGTACCGAGCTCGGATCGATATCTGCGGCCTAGCTAGCGCTACCGGTCGCCACCATGGTGAGCAAGGGCGAGGAGCTGTTCACCGGGGTGGTGCCCATCCTGGTCGAGCTGGACGGCGACGTAAACGGCCACAAGTTCAGCGTGTCCGGCG---GCGAGGGCGATGCCACCTACGGCAAGCTGACCCTGAAGTTCATCTGCACCACCGGCAAGCTGCCCGTGCCCTGGCCCACCCTCGTGACCACCCTGACCTACGGC
## CGCAAATGGGCGGTAGGCGTGTACGGTGGGAGGTCTATATAAGCAGAGCTCTCTGGCTAACTAGAGAACCCACTGCTTACTGGCTTATCGAAATTAATACGACTCACTATAGGGAGACCCAAGCTTGGTACCGAGCTCGGATCGATATCTGCGGCCTAGCTAGCGCTACCGGTCGCCACCATGGTGAGCAAGGGCGAGGAGCTGTTCACCGGGGTGGTGCCCATCCTGGTCGAGCTGGACGGCGACGTAAACGGCCACAAGTTCAGCG---------------------ATGCCACCTACGGCAAGCTGACCCTGAAGTTCATCTGCACCACCGGCAAGCTGCCCGTGCCCTGGCCCACCCTCGTGACCACCCTGACCTACGGC
```

```
cat("Site 3:\n",
print_stats(mut_df, mut_sites = "site3", common = TRUE))
```

```
## Site 3:
##  Mutation Rate: 1880 / 41187 (4.56454706582174%)
##  Indel Rate: 1737 / 41187 (4.21735013475126%)
##  CGCAAATGGGCGGTAGGCGTGTACGGTGGGAGGTCTATATAAGCAGAGCTCTCTGGCTAACTAGAGAACCCACTGCTTACTGGCTTATCGAAATTAATACGACTCACTATAGGGAGACCCAAGCTTGGTACCGAGCTCGGATCGATATCTGCGGCCTAGCTAGCGCTACCGGTCGCCACCATGGTGAGCAAGGGCGAGGAGCTGTTCACCGGGGTGGTGCCCATCCTGGTCGAGCTGGACGGCGACGTAAACGGCCACAAGTTCAGCGTGTCCGGCGAGGGCGAGGGCGATGCCACCTACGGCAAGCTGACCCTGAAGTTCATCTGCACCACCGGCAAAGCTGCCCGTGCCCTGGCCCACCCTCGTGACCACCCTGACCTACGGC
## CGCAAATGGGCGGTAGGCGTGTACGGTGGGAGGTCTATATAAGCAGAGCTCTCTGGCTAACTAGAGAACCCACTGCTTACTGGCTTATCGAAATTAATACGACTCACTATAGGGAGACCCAAGCTTGGTACCGAGCTCGGATCGATATCTGCGGCCTAGCTAGCGCTACCGGTCGCCACCATGGTGAGCAAGGGCGAGGAGCTGTTCACCGGGGTGGTGCCCATCCTGGTCGAGCTGGACGGCGACGTAAACGGCCACAAGTTCAGCGTGTCCGGCGAGGGCGAGGGCGATGCCACCTACGGCAAGCTGACCCTGAAGTTCATCTGCACCACCG-------------GCCCTGGCCCACCCTCGTGACCACCCTGACCTACGGC
## CGCAAATGGGCGGTAGGCGTGTACGGTGGGAGGTCTATATAAGCAGAGCTCTCTGGCTAACTAGAGAACCCACTGCTTACTGGCTTATCGAAATTAATACGACTCACTATAGGGAGACCCAAGCTTGGTACCGAGCTCGGATCGATATCTGCGGCCTAGCTAGCGCTACCGGTCGCCACCATGGTGAGCAAGGGCGAGGAGCTGTTCACCGGGGTGGTGCCCATCCTGGTCGAGCTGGACGGCGACGTAAACGGCCACAAGTTCAGCGTGTCCGGCGAGGGCGAGGGCGATGCCACCTACGGCAAGCTGACCCTGAAGTTCATCTGCACCACCGGCA-GCTGCCCGTGCCCTGGCCCACCCTCGTGACCACCCTGACCTACGGC
## CGCAAATGGGCGGTAGGCGTGTACGGTGGGAGGTCTATATAAGCAGAGCTCTCTGGCTAACTAGAGAACCCACTGCTTACTGGCTTATCGAAATTAATACGACTCACTATAGGGAGACCCAAGCTTGGTACCGAGCTCGGATCGATATCTGCGGCCTAGCTAGCGCTACCGGTCGCCACCATGGTGAGCAAGGGCGAGGAGCTGTTCACCGGGGTGGTGCCCATCCTGGTCGAGCTGGACGGCGACGTAAACGGCCACAAGTTCAGCGTGTCCGGCGAGGGCGAGGGCGATGCCACCTACGGCAAGCTGACCCTGAAGTTCATCTGC----------------CCGTGCCCTGGCCCACCCTCGTGACCACCCTGACCTACGGC
## CGCAAATGGGCGGTAGGCGTGTACGGTGGGAGGTCTATATAAGCAGAGCTCTCTGGCTAACTAGAGAACCCACTGCTTACTGGCTTATCGAAATTAATACGACTCACTATAGGGAGACCCAAGCTTGGTACCGAGCTCGGATCGATATCTGCGGCCTAGCTAGCGCTACCGGTCGCCACCATGGTGAGCAAGGGCGAGGAGCTGTTCACCGGGGTGGTGCCCATCCTGGTCGAGCTGGACGGCGACGTAAACGGCCACAAGTTCAGCGTGTCCGGCGAGGGCGAGGGCGATGCCACCTACGGCAAGCTGACCCTGAAGTTCATCTGCACCACCGGC----TGCCCGTGCCCTGGCCCACCCTCGTGACCACCCTGACCTACGGC
## CGCAAATGGGCGGTAGGCGTGTACGGTGGGAGGTCTATATAAGCAGAGCTCTCTGGCTAACTAGAGAACCCACTGCTTACTGGCTTATCGAAATTAATACGACTCACTATAGGGAGACCCAAGCTTGGTACCGAGCTCGGATCGATATCTGCGGCCTAGCTAGCGCTACCGGTCGCCACCATGGTGAGCAAGGGCGAGGAGCTGTTCACCGGGGTGGTGCCCATCCTGGTCGAGCTGGACGGCGACGTAAACGGCCACAAGTTCAGCGTGTCCGGCGAGGGCGAGGGCGATGCCACCTACGGCAAGCTGACCCTGAAGTTCATCTGCACCACC-----------CGTGCCCTGGCCCACCCTCGTGACCACCCTGACCTACGGC
## CGCAAATGGGCGGTAGGCGTGTACGGTGGGAGGTCTATATAAGCAGAGCTCTCTGGCTAACTAGAGAACCCACTGCTTACTGGCTTATCGAAATTAATACGACTCACTATAGGGAGACCCAAGCTTGGTACCGAGCTCGGATCGATATCTGCGGCCTAGCTAGCGCTACCGGTCGCCACCATGGTGAGCAAGGGCGAGGAGCTGTTCACCGGGGTGGTGCCCATCCTGGTCGAGCTGGACGGCGACGTAAACGGCCACAAGTTCAGCGTGTCCGGCGAGGGCGAGGGCGATGCCACCTACGGCAAGCTGACCCTGAAGTTCATCTGCACCACCGGC-------------------CCACCCTCGTGACCACCCTGACCTACGGC
## CGCAAATGGGCGGTAGGCGTGTACGGTGGGAGGTCTATATAAGCAGAGCTCTCTGGCTAACTAGAGAACCCACTGCTTACTGGCTTATCGAAATTAATACGACTCACTATAGGGAGACCCAAGCTTGGTACCGAGCTCGGATCGATATCTGCGGCCTAGCTAGCGCTACCGGTCGCCACCATGGTGAGCAAGGGCGAGGAGCTGTTCACCGGGGTGGTGCCCATCCTGGTCGAGCTGGACGGCGACGTAAACGGCCACAAGTTCAGCGTGTCCGGCGAGGGCGAGGGCGATGCCACCTACGGCAAGCTGACCCTGAAGTTCATCTGCACCACCGGCA---------------GGCCCACCCTCGTGACCACCCTGACCTACGGC
## CGCAAATGGGCGGTAGGCGTGTACGGTGGGAGGTCTATATAAGCAGAGCTCTCTGGCTAACTAGAGAACCCACTGCTTACTGGCTTATCGAAATTAATACGACTCACTATAGGGAGACCCAAGCTTGGTACCGAGCTCGGATCGATATCTGCGGCCTAGCTAGCGCTACCGGTCGCCACCATGGTGAGCAAGGGCGAGGAGCTGTTCACCGGGGTGGTGCCCATCCTGGTCGAGCTGGACGGCGACGTAAACGGCCACAAGTTCAGCGTGTCCGGCGAGGGCGAGGGCGATGCCACCTACGGCAAGCTGACCCTGAAGTTCATCTGCACCACCGGC-------CCGTGCCCTGGCCCACCCTCGTGACCACCCTGACCTACGGC
## CGCAAATGGGCGGTAGGCGTGTACGGTGGGAGGTCTATATAAGCAGAGCTCTCTGGCTAACTAGAGAACCCACTGCTTACTGGCTTATCGAAATTAATACGACTCACTATAGGGAGACCCAAGCTTGGTACCGAGCTCGGATCGATATCTGCGGCCTAGCTAGCGCTACCGGTCGCCACCATGGTGAGCAAGGGCGAGGAGCTGTTCACCGGGGTGGTGCCCATCCTGGTCGAGCTGGACGGCGACGTAAACGGCCACAAGTTCAGCGTGTCCGGCGAGGGCGAGGGCGATGCCACCTACGGCAAGCTGACCCTGAAGTTCATCTGCAC--------AGCTGCCCGTGCCCTGGCCCACCCTCGTGACCACCCTGACCTACGGC
```

```
cat("Multiple Sites:\n",
print_stats(mut_df, mut_sites = c("site1", "site2", "site3"), 
            min_sites = 2, max_sites = 3, common = TRUE))
```

```
## Multiple Sites:
##  Mutation Rate: 2917 / 41187 (7.08233180372448%)
##  Indel Rate: 2861 / 41187 (6.9463665719766%)
##  CGCAAATGGGCGGTAGGCGTGTACGGTGGGAGGTCTATATAAGCAGAGCTCTCTGGCTAACTAGAGAACCCACTGCTTACTGGCTTATCGAAATTAATACGACTCACTATAGGGAGACCCAAGCTTGGTACCGAGCTCGGATCGATATCTGCGGCCTAGCTAGCGCTACCGGTCGCCACCATGGTGAGCAAGGGCGAGGAGCTGTTCACCGGGGT------------------------------------------------------------------------------------------------------------------------------------GCCCTGGCCCACCCTCGTGACCACCCTGACCTACGGC
## CGCAAATGGGCGGTAGGCGTGTACGGTGGGAGGTCTATATAAGCAGAGCTCTCTGGCTAACTAGAGAACCCACTGCTTACTGGCTTATCGAAATTAATACGACTCACTATAGGGAGACCCAAGCTTGGTACCGAGCTCGGATCGATATCTGCGGCCTAGCTAGCGCTACCGGTCGCCACCATGGTGAGCAAGGGCGAGGAGCTGTTCACCGGGGTGGTGCCCA---------------------------------------------------------------------------------------------------------------------------------------CCCTCGTGACCACCCTGACCTACGGC
## CGCAAATGGGCGGTAGGCGTGTACGGTGGGAGGTCTATATAAGCAGAGCTCTCTGGCTAACTAGAGAACCCACTGCTTACTGGCTTATCGAAATTAATACGACTCACTATAGGGAGACCCAAGCTTGGTACCGAGCTCGGATCGATATCTGCGGCCTAGCTAGCGCTACCGGTCGCCACCATGGTGAGCAAGGGCGAGG---------------------------------------------------------------------------------------GCGATGCCACCTACGGCAAGCTGACCCTGAAGTTCATCTGCACCACCGGCAAGCTGCCCGTGCCCTGGCCCACCCTCGTGACCACCCTGACCTACGGC
## CGCAAATGGGCGGTAGGCGTGTACGGTGGGAGGTCTATATAAGCAGAGCTCTCTGGCTAACTAGAGAACCCACTGCTTACTGGCTTATCGAAATTAATACGACTCACTATAGGGAGACCCAAGCTTGGTACCGAGCTCGGATCGATATCTGCGGCCTAGCTAGCGCTACCGGTCGCCACCATGGTGAGCAAGGGCGAGGAGCTGTTCACCGGGGTGGTGCCCATCCTGGTCGAGCTGGACGGCGACGTAAACGGCCACAAGTTCAGCGTGTCC---------------------------------------------------------ACCGGCAAGCTGCCCGTGCCCTGGCCCACCCTCGTGACCACCCTGACCTACGGC
## CGCAAATGGGCGGTAGGCGTGTACGGTGGGAGGTCTATATAAGCAGAGCTCTCTGGCTAACTAGAGAACCCACC---------------------------------------------------------------------------------------------------------------------------------------------------------------------------------------------------------------------CGATGCCACCTACGGCAAGCTGACCCTGAAGTTCATCTGCACCACCGGCAAGCTGCCCGTGCCCTGGCCCACCCTCGTGACCACCCTGACCTACGGC
## CGCAAATGGGCGGTAGGCGTGTACGGTGGGAGGTCTATATAAGCAGAGCTCTCTGGCTAACTAGAGAACCCACTGCTTACTGGCTTATCGAAATTAATACGACTCACTATAGGGAGACCCAAGCTTGGTACCGAGCTCGGATCGATATCTGCGGCCTAGCTAGCGCTACCGGTCGCCACCATGGTGAGCAAGGGCGAGGAGCTGTTCACCGGGGTGGTGCCCATCCTGGTCGAGCTGGACGGCGACGTAAACGGCCACAAGTTCAGCGTGTCCGGCGAGGGCGA------TGCCACCTACGGCAAGCTGACCCTGAAGTTCATCTGCACCACCGGCAAAGCTGCCCGTGCCCTGGCCCACCCTCGTGACCACCCTGACCTACGGC
## CGCAAATGGGCGGTAGGCGTGTACGGTGGGAGGTCTATATAAGCAGAGCTCTCTGGCTAACTAGAGAACCCACTGCTTACTGGCTTATCGAAATTAATACGACTCACTATAGGGAGACCCAAGCTTGGTACCGAGCTCGGATCGATATCTGCGGCCTAGCTAGCGCTACCGGTCGCCACCATGGTGAGCAAGGGCGAGGAGCTGTTCACCGGGGTGGTGCCCATCCTGGTCGAGCTGGACGGCGACGTAAACGGCCACAAGTTCA------------------------------------------------------TCATCTGCACCACCGGCAAGCTGCCCGTGCCCTGGCCCACCCTCGTGACCACCCTGACCTACGGC
## CGCAAATGGGCGGTAGGCGTGTACGGTGGGAGGTCTATATAAGCAGAGCTCTCTGGCTAACTAGAGAACCCACTGCTTACTGGCTTATCGAAATTAATACGACTCACTATAGGGAGACCCAAGCTTGGTACCGAGCTCGGATCGATATCTGCGGCCTAGCTAGCGCTACCGGTCGCCACCATGGTGAGCAAGGGCGAGGAGCTGTTCACCGGGGTGGTGCCCATCCTGGTCGAGCTGGACGGCGACGTAAACGGCCACAAGTTCAGCGTGTCCGGCGAGG-CGAGGGCGATGCCACCTACGGCAAGCTGACCCTGAAGTTCATCTGCACCACCGGCA-GCTGCCCGTGCCCTGGCCCACCCTCGTGACCACCCTGACCTACGGC
## CGCAAATGGGCGGTAGGCGTGTACGGTGGGAGGTCTATATAAGCAGAGCTCTCTGGCTAACTAGAGAACCCACTGCTTACTGGCTTATCGAAATTAATACGACTCACTATAGGGAGACCCAAGCTTGGTACCGAGCTCGGATCGATATCTGCGGCCTAGCTAGCGCTACC-------------------------------------------------------------------------------------------------------------------------------------------------------------ACCACCGGCAAGCTGCCCGTGCCCTGGCCCACCCTCGTGACCACCCTGACCTACGGC
## CGCAAATGGGCGGTAGGCGTGTACGGTGGGAGGTCTATATAAGCAGAGCTCTCTGGCTAACTAGAGAACCCACTGCTTACTGGCTTATCGAA-----------------------------------------------------------------------------------------------------------------------------------------------------------------------------------------------------------------------------------------------------------------CCTGGCCCACCCTCGTGACCACCCTGACCTACGGC
```
