## Supplemental Note 5 for "SLALOM: A Simple and Rapid Method for Enzymatic Synthesis of CRISPR-Cas9 sgRNA Libraries": SupplementalNote5.html

CrispyR analysis of GFP Screen: Site 4


### CrispyR analysis of GFP Screen: Site 4

###### Jonathon Hill

Cris-py is designed to analyze indels from CRISPR mutagenesis. However, it is struggling with my files and saying none have any valid sequences, but I can find them when I look manually. I will re-create the code here to do my own analysis.

Cris-py steps: 1. Identify sequences and reverse compliment if necessary. 2. Measure lengths to get overall Indel percentage. 3. Look for deletion of specific guides to attribute Indels.

One issue is that Cris-py is not designed for close sites where deletions may cover more than one sgRNA site. I will address that here.

#### Inputs

```
start_seq <- toupper("gccctgagcaaagaccccaacg")
end_seq <- toupper("tcatccgggagcagcaccccac")
sites <- toupper(c(site4 = "gttcgtgaccgccgccgggatcactctcgg")) 
wt_seq <- toupper(gsub("[ \t\r\n]", "", 
                       "gccctgagcaaagaccccaacgagaagcgcgatcacatggtcctgctggagttcgt
                        gaccgccgccgggatcactctcggcatggacgagctgtacaagtccggactcagat
                        ctagaccgtccgagaagaccttcaaacagcgccggagcttcgaacaaagagtggaa
                        gatgtccggctcatccgggagcagcaccccac"))
pcr_size <- nchar(wt_seq)
con_fastq <- "L2Con.extendedFrags.fastq"
pos_fastq <- "L2Plus.extendedFrags.fastq"
neg_fastq <- "L2Minus.extendedFrags.fastq"
```

#### Control

```
seqs <- read_fastq(con_fastq)$Seq
filtered_seqs <- process_seqs(seqs, start_seq, end_seq)
```

```
## Number Valid/Total: 16127 / 18599
```

```
mut_df <- get_mut_df(filtered_seqs, sites, pcr_size)
cat("Site 4:\n",
print_stats(mut_df, mut_sites = "site4"))
```

```
## Site 4:
##  Mutation Rate: 153 / 16127 (0.948719538661872%)
##  Indel Rate: 3 / 16127 (0.0186023438953308%)
```

#### GFP Positive Pool

```
seqs <- read_fastq(pos_fastq)$Seq
filtered_seqs <- process_seqs(seqs, start_seq, end_seq)
```

```
## Number Valid/Total: 14494 / 16011
```

```
mut_df <- get_mut_df(filtered_seqs, sites, pcr_size)
cat("Site 4:\n",
print_stats(mut_df, mut_sites = "site4"))
```

```
## Site 4:
##  Mutation Rate: 197 / 14494 (1.35918311025252%)
##  Indel Rate: 73 / 14494 (0.503656685525045%)
```

#### GFP Negative Pool

```
seqs <- read_fastq(neg_fastq)$Seq
filtered_seqs <- process_seqs(seqs, start_seq, end_seq)
```

```
## Number Valid/Total: 16707 / 18507
```

```
mut_df <- get_mut_df(filtered_seqs, sites, pcr_size)
cat("Site 4:\n",
print_stats(mut_df, mut_sites = "site4", common = TRUE))
```

```
## Site 4:
##  Mutation Rate: 486 / 16707 (2.90896031603519%)
##  Indel Rate: 333 / 16707 (1.99317651283893%)
##  GCCCTGAGCAAAGACCCCAACGAGAAGCGCGATCACATGGTCCTGCTGGAGTTCGTGAC-----------TCACTCTCGGCATGGACGAGCTGTACAAGTCCGGACTCAGATCTAGACCGTCCGAGAAGACCTTCAAACAGCGCCGGAGCTTCGAACAAAGAGTGGAAGATGTCCGGCTCATCCGGGAGCAGCACCCCAC
## GCCCTGAGCAAAGACCCCAACGAGAAGCGCGATCACATGGTCCTGCTGGAGTTC------------------ACTCTCGGCATGGACGAGCTGTACAAGTCCGGACTCAGATCTAGACCGTCCGAGAAGACCTTCAAACAGCGCCGGAGCTTCGAACAAAGAGTGGAAGATGTCCGGCTCATCCGGGAGCAGCACCCCAC
## GCCCTGAGCAAAGACCCCAACGAGAAGCGCGATCACATGGTCCTGCTGGA-----------------------CTCTCGGCATGGACGAGCTGTACAAGTCCGGACTCAGATCTAGACCGTCCGAGAAGACCTTCAAACAGCGCCGGAGCTTCGAACAAAGAGTGGAAGATGTCCGGCTCATCCGGGAGCAGCACCCCAC
## GCCCTGAGCAAAGACCCCAACGAGAAGCGCGATCACATGGTCCTGCTGGAGTTC----------------------TCGGCATGGACGAGCTGTACAAGTCCGGACTCAGATCTAGACCGTCCGAGAAGACCTTCAAACAGCGCCGGAGCTTCGAACAAAGAGTGGAAGATGTCCGGCTCATCCGGGAGCAGCACCCCAC
## GCCCTGAGCAAAGACCCCAACGAGAAGCGCGATCACATGGTCCTGCTGGAGTTCGCGACCGCCGCCGGGATCACTCTCGGCATGGACGAGCTGTACAAGTCCGGACTCAGATCTAGACCGTCCGAGAAGACCTTCAAACAGCGCCGGAGCTTCGAACAAAGAGTGGAAGATGTCCGGCTCATCCGGGAGCAGCACCCCAC
## GCCCTGAGCAAAGACCCCAACGAGAAGCGCGATCACATGGTCCTGCTGGAGTTCGTGACCGCCGCCGGGAT------------GGACGAGCTGTACAAGTCCGGACTCAGATCTAGACCGTCCGAGAAGACCTTCAAACAGCGCCGGAGCTTCGAACAAAGAGTGGAAGATGTCCGGCTCATCCGGGAGCAGCACCCCAC
## GCCCTGAGCAAAGACCCCAACGAGAAGCGCGATCACATGGTCCTGCTGGAGTTCGTGACCGCCGCCGGGATTCACTCTCGGCATGGACGAGCTGTACAAGTCCGGACTCAGATCTAGACCGTCCGAGAAGACCTTCAAACAGCGCCGGAGCTTCGAACAAAGAGTGGAAGATGTCCGGCTCATCCGGGAGCAGCACCCCAC
## GCCCTGAGCAAAGACCCCAACGAGAAGCGCGATCACATGGTCCTGCTGGAGTTCGTGAC---------------TCTCGGCATGGACGAGCTGTACAAGTCCGGACTCAGATCTAGACCGTCCGAGAAGACCTTCAAACAGCGCCGGAGCTTCGAACAAAGAGTGGAAGATGTCCGGCTCATCCGGGAGCAGCACCCCAC
## GCCCTGAGCAAAGACCCCAACGAGAAGCGCGATCACATGGTCCTGCTGGAGTTCGTGAC-------------ACTCTCGGCATGGACGAGCTGTACAAGTCCGGACTCAGATCTAGACCGTCCGAGAAGACCTTCAAACAGCGCCGGAGCTTCGAACAAAGAGTGGAAGATGTCCGGCTCATCCGGGAGCAGCACCCCAC
## GCCCTGAGCAAAGACCCCAACGAGAAGCGCGATCACATGGTCCTGCTGGAGTTCGTGACCGCCGCCGGGGT------------GGACGAGCTGTACAAGTCCGGACTCAGATCTAGACCGTCCGAGAAGACCTTCAAACAGCGCCGGAGCTTCGAACAAAGAGTGGAAGATGTCCGGCTCATCCGGGAGCAGCACCCCAC
```
